# Master of none: *GPRC6A* gene loss is more widespread than previously known

**DOI:** 10.1101/2025.11.14.688574

**Authors:** Saumya Gupta, Ajinkya Bharatraj Patil, Aswin S Soman, Nagarjun Vijay

**Affiliations:** Computational Evolutionary Genomics Lab, Department of Biological Sciences, IISER Bhopal, Bhauri, Madhya Pradesh, India

**Keywords:** *GPRC6A*, gene loss, dispensable, diet, pseudogenisation, conditional dispensability

## Abstract

*GPRC6A* encodes a class C GPCR that can be activated by multiple ligands and potentially acts as a central regulator of diverse metabolic processes by modulating endocrine pathways. Experimental studies have reported numerous distinct functions for *GPRC6A*, suggesting it may be a key drug target for several metabolic disorders. Yet, the actual function of *GPRC6A* has been the focus of considerable debate due to contradictory results and the prevalence of loss-of-function mutations in human populations, leading to the perception of *GPRC6A* as a “Master of none”. Interestingly, a genome-wide screen for gene loss events in vertebrate species identified the disruption of the *GPRC6A* gene in toothed whales, in contrast to widespread conservation in the closely related Bovidae family. We employ a synteny-informed comparative genomic approach to demonstrate that the loss of the *GPRC6A* gene among mammalian species is more widespread than previously reported, encompassing the entire Bovidae group within Artiodactyla and other fully aquatic mammals, including those belonging to Sirenia. An in-depth search of the genomes and short and long-read sequencing datasets of monotremes, hystricomorphs, rhinolophoid bats, pika, koala, and two shrews (white-toothed pygmy shrew and Asian house shrew) reveals at least nine independent *GPRC6A* gene loss events in vertebrates, highlighting its lineage-specific dispensability and raising questions regarding its ubiquitous functionality. The evolutionary loss of *GPRC6A* likely represents a lineage-specific response to specialised diets and ecological niches, reshaping metabolic regulation and taste perception and illuminating how niche specialisation influences gene retention or loss within the GPCR landscape across species.

## Introduction

G protein-coupled receptors (GPCRs) are located on the cell membrane and sense specific molecules before mediating a downstream cellular response (Rosenbaum et al. 2009; Yang et al. 2021). While GPCRs are 7TM (seven transmembrane) receptors that serve as communication nodes between the exterior and interior of cells, the specific ligands that mediate this signalling, their mode of action, and the downstream functions are varied (Pierce et al. 2002; Xue et al. 2008; Sladek and Song 2012; Heng et al. 2013). GPCRs are the largest family of membrane proteins, comprising hundreds of members classified into four major classes based on amino acid sequence identity (Yang et al. 2021). *GPRC6A* is grouped under Class C (glutamate) along with one calcium-sensing receptor (CaSR), three taste-1 receptors (TS1R1), two gamma-aminobutyric acid (GABA) type B receptors, eight metabotropic glutamate receptors, and other orphan GPCRs (Wellendorph and Bräuner-Osborne 2004).

*GPRC6A* serves as a master regulator of multiple ligands, with osteocalcin being a crucial activator that influences its regulatory roles in metabolic processes. Osteocalcin, primarily released from bone, activates *GPRC6A* across various tissues, particularly in adipose tissues and muscles, where it is essential for lipid metabolism, energy homeostasis, and glucose regulation through the modulation of insulin secretion and sensitivity (Ferron et al. 2008; Pi et al. 2017; Mukai et al. 2021; Karsenty 2024). *GPRC6A* knockout (KO) mice exhibit metabolic dysregulation, including glucose intolerance and insulin resistance, which mirrors the metabolic and skeletal defects observed in osteocalcin KO mice (Pi et al. 2008, 2015; Wei et al. 2014). Testosterone functions as another significant ligand for *GPRC6A*, modulating metabolic and reproductive functions, particularly in the testes and prostate, where it influences testosterone synthesis and secretion, linking *GPRC6A* to androgen-related pathways in male reproductive physiology (Pi et al. 2010; De Toni et al. 2016b). *GPRC6A* is also activated by basic amino acids, such as L-arginine and L-lysine, underscoring its role in nutrient signalling (Kuang et al. 2005; Oya et al. 2013), while divalent cations, including calcium and magnesium, highlight its importance in mineral ion homeostasis (Pi et al. 2005; Christiansen et al. 2007).

The coding region of the longest isoform of the human *GPRC6A* gene comprises six exons, with the Venus Fly Trap domain spanning exons 1 to 3 and the transmembrane and C-terminal domains located in exon 6 (Pi et al. 2017). Shorter isoforms skip in-frame exon 4 or partially include exon 3. Knockout studies reveal isoform-specific functionalities in *GPRC6A*, with natural polymorphisms linked to insulin resistance and testis-related phenotypes (De Toni et al. 2016a; Di Nisio et al. 2017; Jørgensen and Bräuner-Osborne 2020; Jawich et al. 2022). *GPRC6A* also plays immunological roles such as the expansion and function of innate lymphoid cells type 3 (ILC3s) and boosts interleukin-22 (IL-22) production, thereby modulating immune responses and promoting mucosal healing through L-arginine activation (Hou et al. 2022). Recently, *GPRC6A* has also been implicated in taste perception, specifically in the kokumi taste, suggesting its broader role in sensory and metabolic signalling (Yamamoto et al. 2024). Due to such diverse functions, *GPRC6A* has been described as the “Jack of all metabolism” (Pi et al. 2017). However, the validity of such ubiquitous functionality has been questioned (Jacobsen et al. 2013, 2017; Booth et al. 2014; Rueda et al. 2016; Jørgensen et al. 2019; Yanagisawa 2023), especially in light of the *GPRC6A* loss-of-function gene variant (rs6907580) identified in human populations (MacArthur et al. 2012; Jørgensen et al. 2017; Jørgensen and Bräuner-Osborne 2020). Hence, the alternative view is that *GPRC6A* is dispensable with minimal clinical relevance, a “Master of none” (Pi et al. 2017).

The extreme views of ubiquitous vs. rare or non-existent functions attributed to *GPRC6A* could result from differing roles of the gene in disparate species or different genomic backgrounds (Pi et al. 2020). Lineage-specific differences in *GPRC6A* functionality are potentially mediated by paralogs, distant structural homologs or alternative pathways with a similar role (Clemmensen et al. 2013, 2014; Pi et al. 2021). For instance, the discovery of *GPRC6A* gene loss in toothed whales is surprising and either supports the dispensability of this gene or could be a result of pathway rewiring, leading to the evolution of blubber or a higher testis-to-body mass ratio in cetaceans relative to other mammals (Turakhia et al. 2020). Species-specific changes in exon usage patterns and the evolution of novel splice isoforms are also possibilities, given the presence of multiple isoforms with skippable in-frame exons (Desai et al. 2022). Hence, comparative genomics can provide crucial clues to understanding the experimental evidence supporting contrasting views about the functionality of *GPRC6A*.

In this study, we performed an in-depth screen of mammalian genomes to evaluate the status of the *GPRC6A* gene to (1) Reconstruct the chronology of coding frame disrupting events across all cetacean species and verify the previously reported instances of *GPRC6A* gene loss in dolphin and killer whale, (2) Evaluate whether the claims of conservation of *GPRC6A* in outgroup bovidae (domestic goat, sheep, Tibetan antelope, and cow) species is supported by genomic datasets, (3) Identify other instances of *GPRC6A* gene loss among mammalian species and correct genome assembly or annotation errors, (4) Quantify the selection regimes acting on the *GPRC6A* gene in various mammalian clades. Our computational analysis of diverse genomic datasets suggests that the loss of the *GPRC6A* gene is more widespread than previously known, with important implications for understanding its biology in commercially important species such as cattle.

## RESULTS

### Conservation of the *GPRC6A* syntenic region across mammals

The genes flanking *GPRC6A* are conserved across mammals and have been unambiguously identified in the high-quality genomes of the representative species considered (see **Fig. 1**, **Supplementary Figures S1-S27**, **Supplementary Table S1-S13**). In most species, the genes *ZUP1*, *KPNA5*, and *FAM162B* are located upstream of *GPRC6A* and are separated by approximately 200 kilobases (kb). With a few exceptions, the genes downstream from *GPRC6A* are *RFX6*, *VGLL2*, and *ROS1*. While the *RFX6* gene is adjacent to *GPRC6A*, the *VGLL2* and *ROS1* genes are approximately 1Mb away in the human genome, and the intervening region contains pseudogenes such as ribosomal protein S29 pseudogene 13, 5S ribosomal pseudogene 214, *RN7SK* pseudogene 18 and *RN7SK* pseudogene 51 (**Supplementary Figure S1**). Although the same gene order is largely conserved in other mammalian species, *FAM162B* is missing from the genomes of the Norwegian rat (*Rattus norvegicus*), European hedgehog (*Erinaceus europaeus*), and wombat (*Vombatus ursinus*), and is potentially lost (**Supplementary Figures S2-S4**). Similarly, the *KPNA5* gene is absent in the house mouse (*Mus musculus*) and several Mus species, including the Ryukyu mouse (*Mus caroli*) and Gairdner’s shrewmouse (*Mus pahari*) (**Supplementary Figures S5-S8**). The entirety of the syntenic region encompassing *GPRC6A* and *FAM162B* appears absent in Monotremata (**Supplementary Figures S9-S10**). The gene order was conserved in the two closely related outgroup species, chicken (*Gallus gallus*) and European common frog (*Rana temporaria*), except for *KPNA5*, which did not yield a BLAST hit in the frog (**Supplementary Figures S11-S12**).

**Fig. 1.**
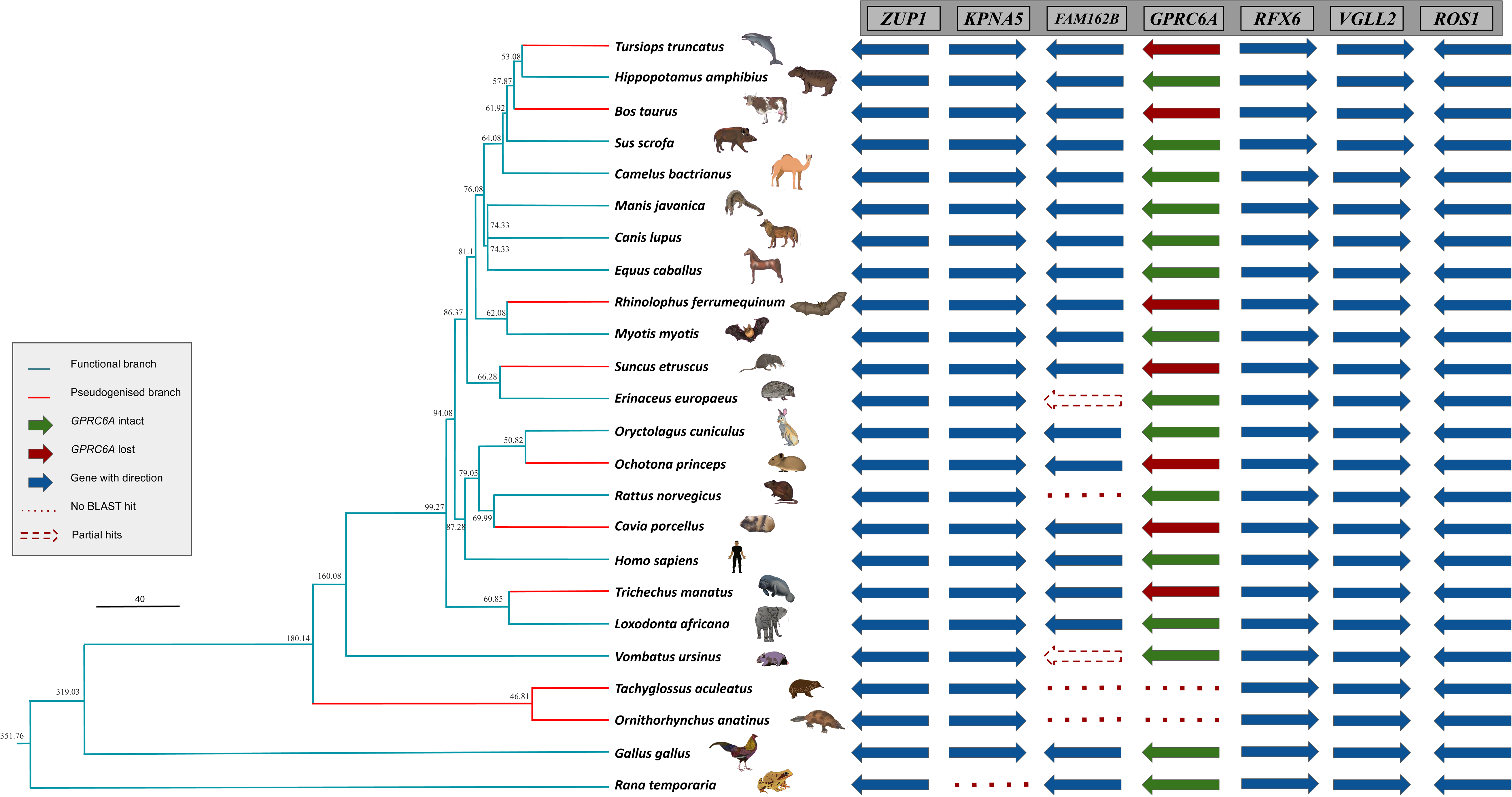
Conservation of synteny establishes orthology of *GPRC6A* in mammals. The syntenic region flanking the *GPRC6A* gene is illustrated, with arrowheads indicating the direction of transcription. Genes absent from the syntenic location, as determined by lack of BLAST hits, are represented by red dotted lines. The green arrow indicates an intact *GPRC6A* gene, whereas the red arrow signifies the presence of a premature stop codon within the gene. Partially recovered genes from genome BLAST are shown with red dotted arrows. The conserved upstream genes include *RFX6* (Regulatory Factor X6), *VGLL2* (Vestigial Like Family Member 2), and *ROS1* (ROS Proto-Oncogene 1, Receptor Tyrosine Kinase). The downstream genes are *FAM162B* (Family With Sequence Similarity 162 Member B), *KPNA5* (Karyopherin Subunit Alpha 5), and *ZUP1* (Zinc Finger Containing Ubiquitin Peptidase 1). The functional branches of *GPRC6A* are indicated by green-colored branches in the phylogenetic tree, while red-colored branches denote a loss of function. This visual representation highlights the evolutionary changes and functional status of the *GPRC6A* gene across different species.

We also analysed GC content across annotated mammalian *GPRC6A* coding sequences to evaluate its impact on sequencing accuracy. Our findings revealed that all annotated members of Caviomorpha exhibit high GC content. Additionally, three insectivorous bat species within the Vespertilionidae family and several non-Hystricomorph rodents also displayed elevated GC content, characterised by a proportion of high-GC regions exceeding 0.3 and a mean GC content above 3.45 (**Supplementary Figure S33**). RNA-seq expression data across tissues from forty-three representative species were analysed to evaluate gene expression (**Supplementary Figures S34-S77)**.

### Chronology of events leading to the loss of *GPRC6A* in Cetacea

A previous genome-wide screen identified the loss of the *GPRC6A* gene in Odontoceti (toothed whales) based on the analysis of dolphin and killer whale genomes (Turakhia et al. 2020). However, the status of the *GPRC6A* gene in other cetacean species and the chronology of events involved in gene loss are unknown. We screened the genomes of 24 cetacean species, comprising both toothed (Odontoceti) and baleen (Mysticeti) whales, to evaluate the presence of gene-disrupting mutations, identify events shared across species, and estimate the putative timeline of gene loss. All cetacean genomes analysed in this study shared two coding-frame-altering events in exon 3 of the gene (see **Fig. 2**, the positions are relative to the *Hippopotamus amphibius* reference sequence). The first is a single-base pair deletion of cytosine (C) at the 269^th^ codon (position 806 bp). The second is a two–base pair insertion of “GC” after the 361^st^ codon (positions 1084-1085 bp). Apart from these shared events, consistent with an ancient loss followed by the gradual degradation of the gene, many other gene-disrupting changes have occurred independently in various cetacean species. Several cetacean species were found to have entire exons missing from their genomes. While other gene-disrupting changes already confirm the loss of the *GPRC6A* gene, we used long-read sequencing as an additional validation step to verify the complete loss of these exons in the Common minke whale (*Balaenoptera acutorostrata*), Blue whale (*Balaenoptera musculus*), Blainville’s beaked whale (*Mesoplodon densirostris*), Narrow-ridged finless porpoise (*Neophocaena asiaorientalis*) and killer whale (*Orcinus orcas*) genomes (**Supplementary Figure S78-S82)**. Outgroup species, such as the Bactrian camel (*Camelus bactrianus*), wild boar (*Sus scrofa*) and the hippopotamus (*Hippopotamus amphibius*), lack gene-disrupting mutations and suggest the loss of *GPRC6A* in cetacean species after diverging from hippopotamids ∼55 MYA (Meredith et al. 2011; Tsagkogeorga et al. 2015).

**Fig. 2.**
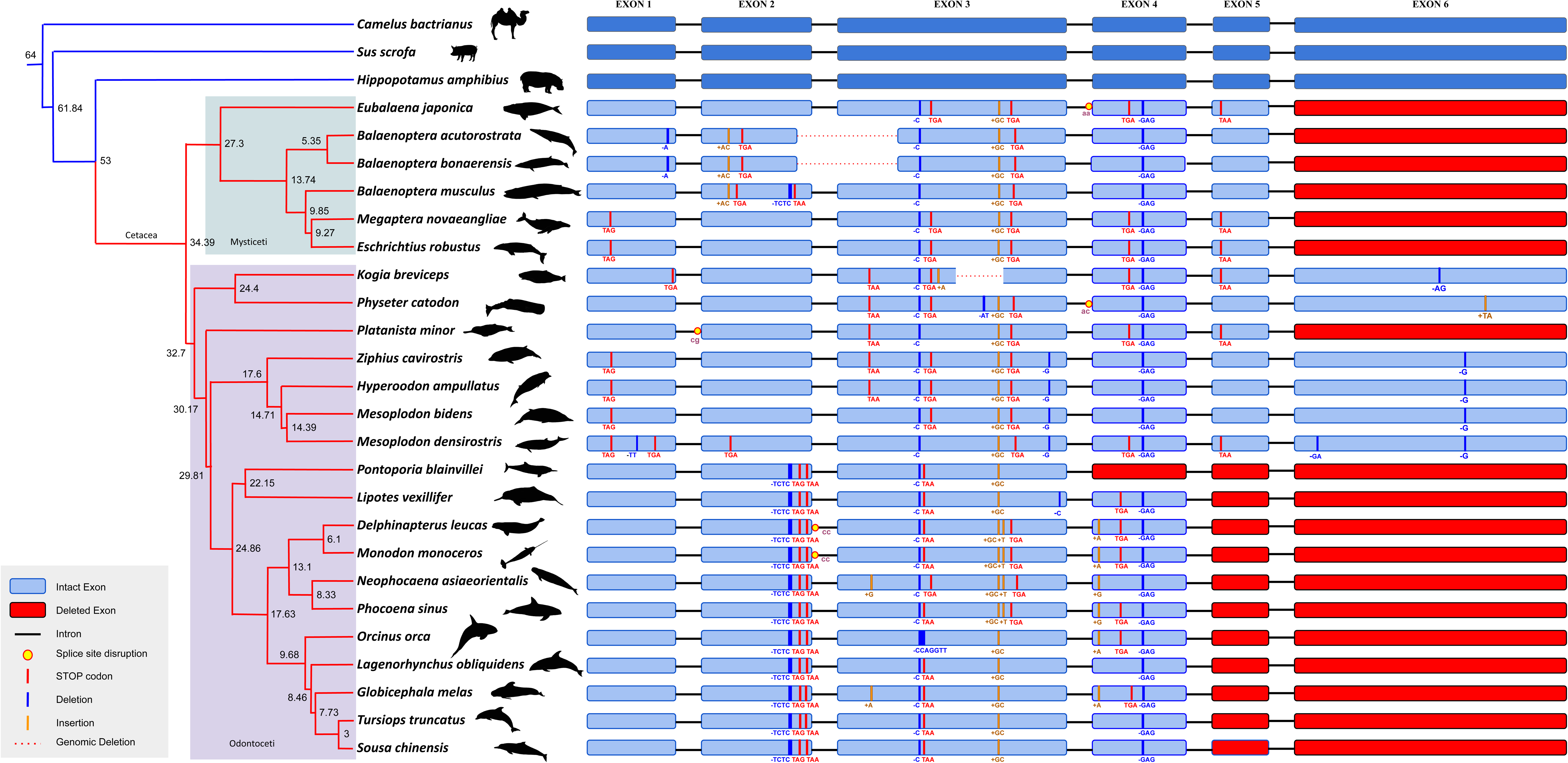
Chronology of GPRC6A gene loss events in Cetacea. The phylogenetic tree, obtained from TimeTree, shows divergence times in millions of years at each node. Blue branches indicate species with intact genes, while red branches represent species with gene loss. Baleen whales (Mysticeti) and toothed whales (Odontoceti) are highlighted in light cyan and light purple, respectively. Exons of intact genes are shown in dark blue, while the species with loss are coloured in a lighter shade of blue. The exons that could not be recovered from the blast search are coloured in red. Splice site disruptions are indicated with yellow and red circles, and genomic deletions are depicted with red dotted lines.

### Loss in Sirenia, another fully aquatic mammalian clade

The same genome-wide study (Turakhia et al. 2020) that identified the loss of the *GPRC6A* gene in cetacean species also screened the genome of the manatee (*Trichechus manatus*) and identified the loss of other genes. However, no evidence for the loss of *GPRC6A* in Sirenia was presented in earlier studies. Sirenia and Cetacea are comparable groups, being fully aquatic mammals. Therefore, we reasoned that a detailed investigation of *GPRC6A* conservation in Sirenia could provide additional insight. The order Sirenia includes two extant genera, namely *Trichechus* and *Dugong*. We identified a short-read validated shared cytosine-to-thymine (C→T) substitution (see **Fig. 3**) in the first exon, resulting in a premature stop codon in both manatee species (*Trichechus manatus* and *Trichechus senegalensis*) and the dugong (*Dugong dugon).* Additionally, we observed a lack of a functional start codon in the coding sequence of *GPRC6A* in these three species. We also identified a 1-base pair deletion in the manatee and a 4-base pair insertion in the dugong, both of which further disrupted the coding frames. These disruptive mutations suggest the absence of a functional *GPRC6A* gene across Sirenia, consistent with the gene’s dispensability in fully aquatic lineages (**Supplementary Text 1**).

**Fig. 3.**
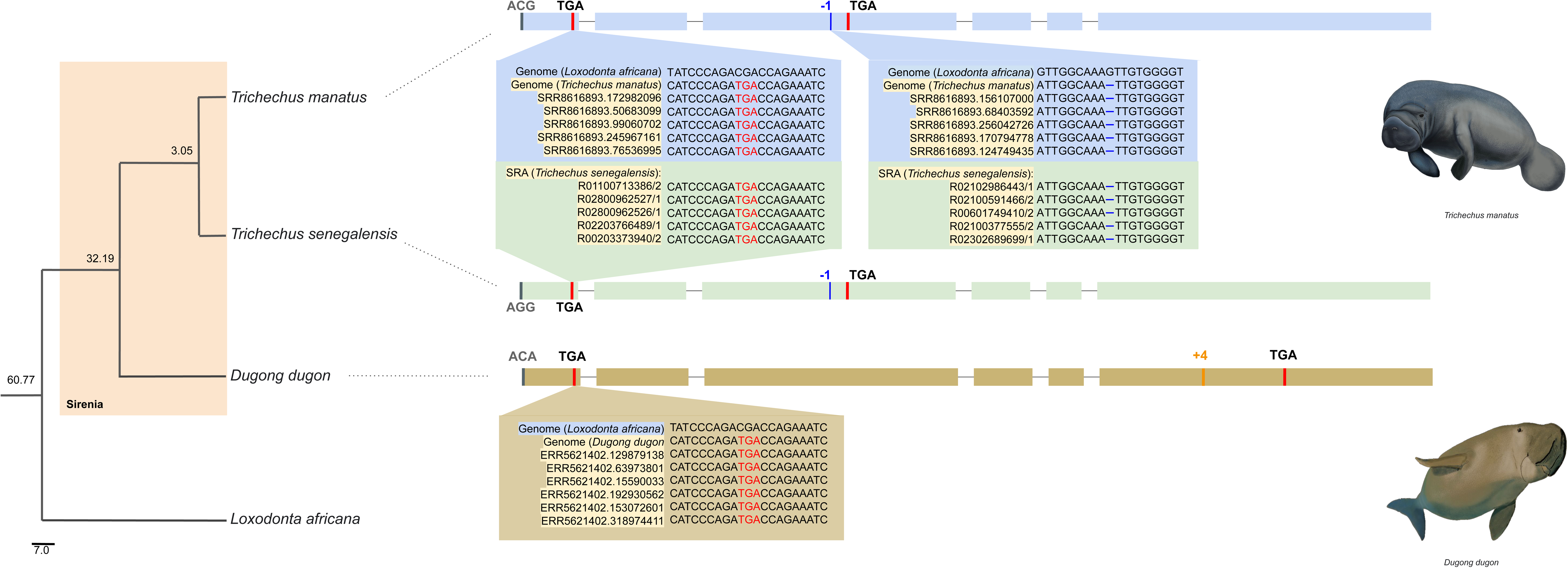
Loss of the GPRC6A gene in Sirenia. Comparative genomic analysis reveals the disruption of the *GPRC6A* gene in the manatee (*Trichechus manatus*) and the dugong (*Dugong dugon*), with the African elephant (*Loxodonta africana*) serving as the reference genome. Exons are represented by blue bars for West Indian manatee (*Trichechus manatus*), as light green bars for the African manatee (*Trichechus senegalensis*) and mustard bars for the dugong (*Dugong dugon*). Red vertical lines indicate premature stop codons, while insertions and deletions are marked by orange and blue lines, respectively, with their lengths (bp) labelled above. Disrupted start codons are highlighted in dark grey. Zoomed-in views display stop codons and a 1 bp deletion in the manatee sequence, validated by short-read alignment, as well as a shared substitution event (C→stop) in both manatee and dugong. On the left, a time-calibrated phylogeny from TimeTree highlights the Sirenia clade and its relationship to *Loxodonta africana*. Divergence times in millions of years are indicated in grey at each node (Heritage and Seiffert 2022).

### Status of *GPRC6A* in ruminants: repeats and other inactivating changes

In contrast to the gene loss reported in toothed whales, earlier studies (Li et al. 2019; Turakhia et al. 2020; Jin et al. 2022) have claimed the widespread conservation of the *GPRC6A* gene, including in ruminants. However, surprisingly, our analysis of ruminant genomes revealed evidence of gene loss through gene-disrupting changes, including the insertion of repeat elements within exonic regions (see **Fig. 4**). A conspicuous shared event among all ruminant genomes investigated is the occurrence of coding frame disruptions, specifically 7-bp and 9-bp deletions within the third exon. Multiple LINE and SINE repeat insertions are also discernible within several exons across various ruminant species. Beyond these shared frame-disrupting mutations, our analyses substantiate a pronounced degradation of the open reading frame (ORF) of the *GPRC6A* gene among ruminants. These findings have been verified by integrating long- and short-read sequencing data (**Supplementary Figure S83-S88**). Notably, the presence of numerous stop codons within the ORF further corroborates our observations. These results are further supported by relaxed selection (see last section of Results) observed in wild boar (*Sus scrofa*), a species closely related to ruminants within Artiodactyla. However, it is important to note that other members of Artiodactyla with intact ORF, for example, Hippopotamus (*Hippopotamus amphibius)* and Bactrian camel (*Camelus bactrianus)*, did not show any sign of relaxed selection, suggesting niche-specific selection pressure on the gene.

**Fig. 4.**
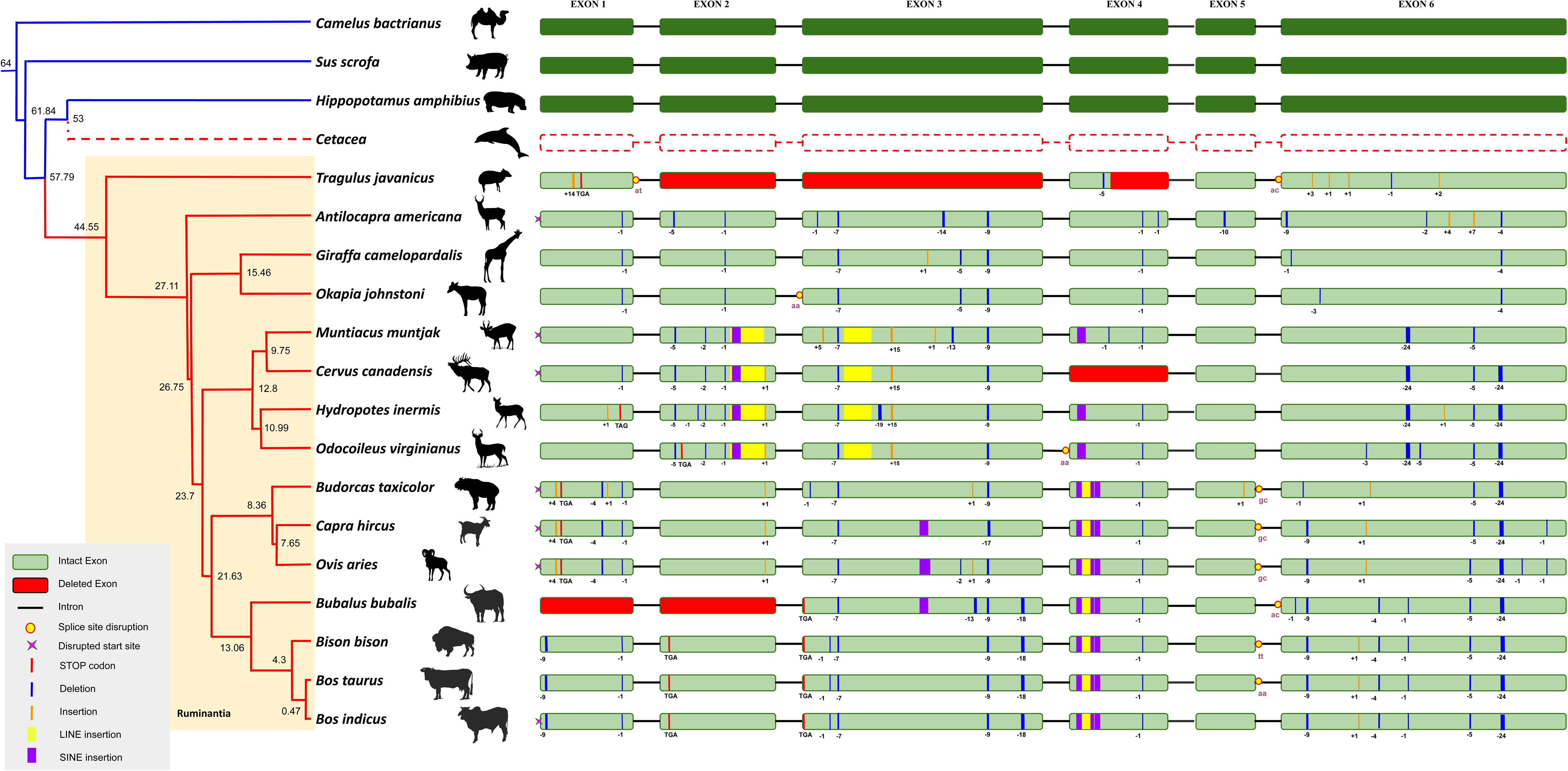
Phylogenetic analysis of GPRC6A loss events in Ruminantia. The phylogenetic tree depicts species relationships within Artiodactyla, with divergence times obtained from TimeTree. The suborder Ruminantia is highlighted in light yellow. Branches in blue indicate species with intact *GPRC6A* genes, while red branches denote species with gene loss. Exons from intact genes are shown in dark green, whereas species with gene loss are indicated in a lighter shade of green. Exons lacking recoverable sequence (based on BLAST searches) are marked in red. Splice site disruptions are highlighted with yellow and red circles, and disrupted start-sites are shown with a pink cross. Insertions are represented by orange bars, deletions by blue, LINE insertions by bright yellow, and SINE insertions by purple. The dotted line denotes gene loss in Cetacea.

### Loss in Caviomorph rodents and Rhinolophoid bats

In our investigation of *GPRC6A*, several rodent and bat species were tagged as “LOW-QUALITY PROTEINS” in GenBank. Upon analysing their genomes and short-read data, we uncovered widespread gene disintegration in three species of Caviomorph rodents: guinea pig (*Cavia porcellus*), common degu (*Octodon degus*), and long-tailed chinchilla (*Chinchilla lanigera*) and found loss in another economically important caviomorph capybara *(Hydrochoerus hydrochaeris*) (**Fig. 5**). We found that the mean GC content in the exonic regions of *GPRC6A* ranged from 40 to 50% in mammals (**Supplementary Table S14-15**). However, the putative exonic regions of caviomorph rodents had a GC content < 50% (**Supplementary Figure S89**), suggesting the sequencing and assembly of these regions are unlikely to be affected by high GC. We extended our analysis across all families with representative genomes to better understand gene loss within Caviomorpha and its potential impact on the broader infraorder Hystricognathi **(Supplementary Figure S90; Supplementary Text 2)**. Our search for *GPRC6A* orthologs revealed extensive gene disintegration across most Caviomorpha families, except for the New World porcupines (Erethizontidae), which retained an intact ORF. Within Phiomorpha, gene loss was found in the Thryonomyidae family, while the ORF remained intact in Bathyergidae and Petromuridae. Additionally, gene disruption was detected in the Old World crested porcupine (*Hystrix cristata*), where the presence of missing exons and stop codons complicated gene reconstruction (**Supplementary Text Section 2**).

**Fig. 5.**
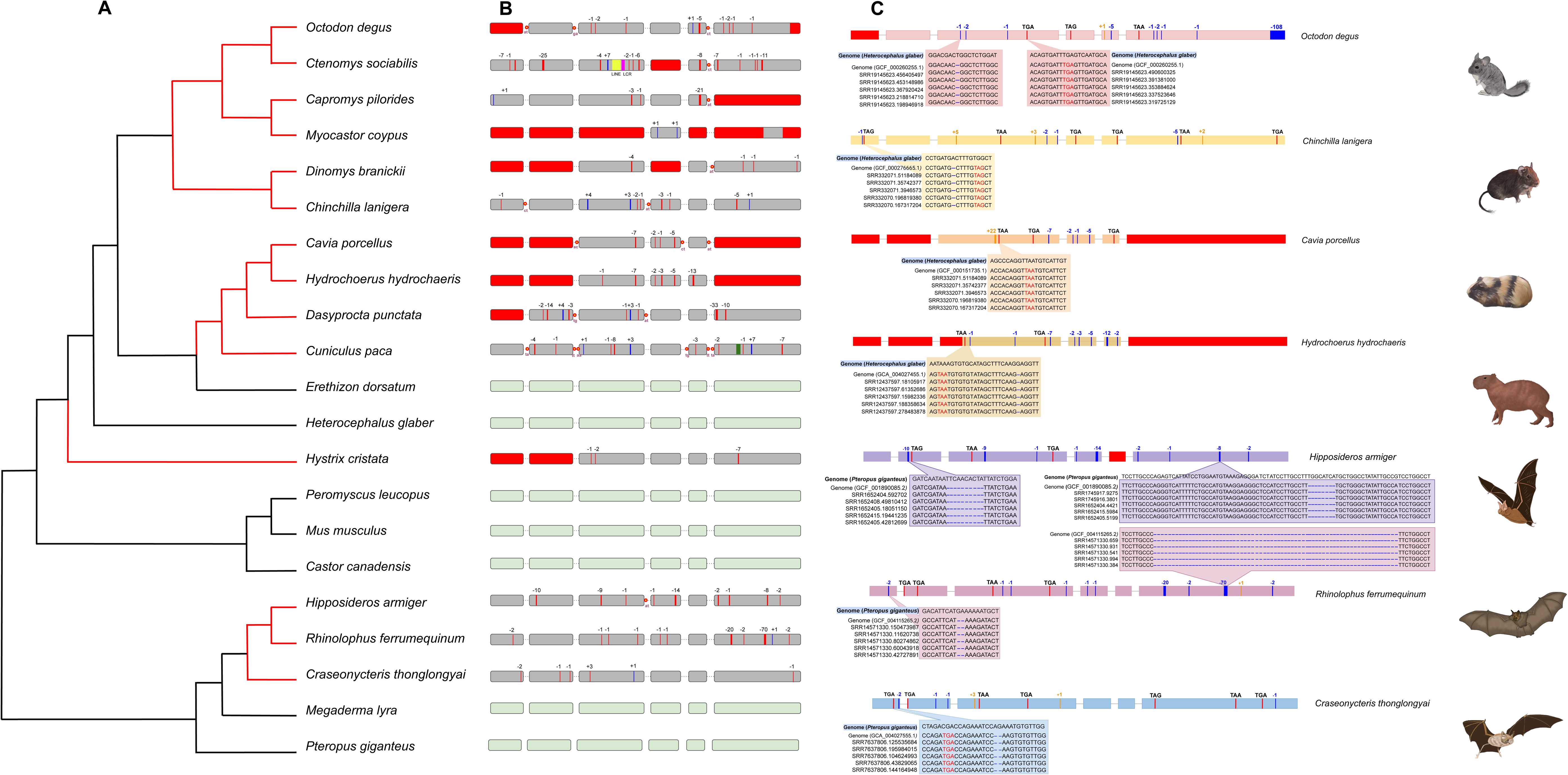
*GPRC6A* gene loss in caviomorph rodents and rhinolophid bats. The left side of the figure displays a phylogenetic tree, with black branches indicating functional *GPRC6A* branches and red branches indicating loss of function in specific species. The right side illustrates loss-of-function events in four representative caviomorph rodents and three rhinolophid bats. The zoomed-in boxes highlight in-frame stop codons and frame-disrupting deletions identified against the reference genome of the closely related naked-mole rat (*Heterocephalus glaber*) in rodents and the Indian flying fox (*Pteropus giganteus*) in Chiroptera, confirmed through short-read alignment.

The two Chiropteran species, the great roundleaf bat (*Hipposideros armiger*) and the greater horseshoe bat (*Rhinolophus ferrumequinum),* have been identified as “LOW-QUALITY” in GenBank. Both species are classified within the Rhinolophoidea superfamily, representing Old World insectivorous bats. Notably, the great roundleaf bat exhibited an 8bp deletion event. In contrast, the greater horseshoe bat displayed a much larger 70bp deletion in the sixth exon, suggesting a shared origin of this deletion between the two species (**Fig. 5**). Furthermore, numerous in-frame stop codons and frame-disrupting mutations further supported our findings. Subsequent evaluation within this clade revealed gene loss in the bumblebee bat also known as kitti’s hog-nosed bat (*Craseonycteris thonglongyai*). However, amidst this variability, it is noteworthy that the *GPRC6A* gene remained preserved in the greater false vampire bat (*Megaderma lyra*) (**Fig. 5**).

### Independent losses in Monotremes, Koala, Pika, and Shrews

During the synteny analysis, it was identified that the *GPRC6A* gene is missing from the genomes of the platypus (*Ornithorhynchus anatinus*) and short-beaked echidna (*Tachyglossus aculeatus*). The pairwise alignment between human and platypus genomes at the region containing the *GPRC6A* gene also supports gene removal by a large genomic deletion (see **Fig. 6A**). The possibility of a genome assembly error/gap was ruled out by verification of the most recent version of the platypus genome using PacBio long-read sequencing (**Supplementary Figure S91**). In the genome of the koala species (*Phascolarctos cinereus*), the absence of the first and third exons indicates the loss of the *GPRC6A* gene (see **Fig. 6B**). We confirmed the exon deletions through analysis of both short-read Illumina and long-read Nanopore sequencing, ruling out assembly errors and long-range inaccuracies (**Supplementary Figures S92-S93)**. Additionally, a chromosome alignment in chain format between the koala and the common wombat (*Vombatus ursinu*s) revealed a single chain encompassing the remaining exons and introns of the *GPRC6A* gene and the syntenic region (**Supplementary Figure S94**). We examined the motif and domain structure of the remaining *GPRC6A* sequence in the koala and compared it with the domain structure of human *GPRC6A* to evaluate the likelihood of functional protein presence despite the absence of two exons. Our analysis indicated disintegration of the Venus Flytrap (VFT) domain in the koala sequence, suggesting that the possibility of a functional *GPRC6A* in this species is minimal. **(Supplementary Text 3**).

**Fig. 6.**
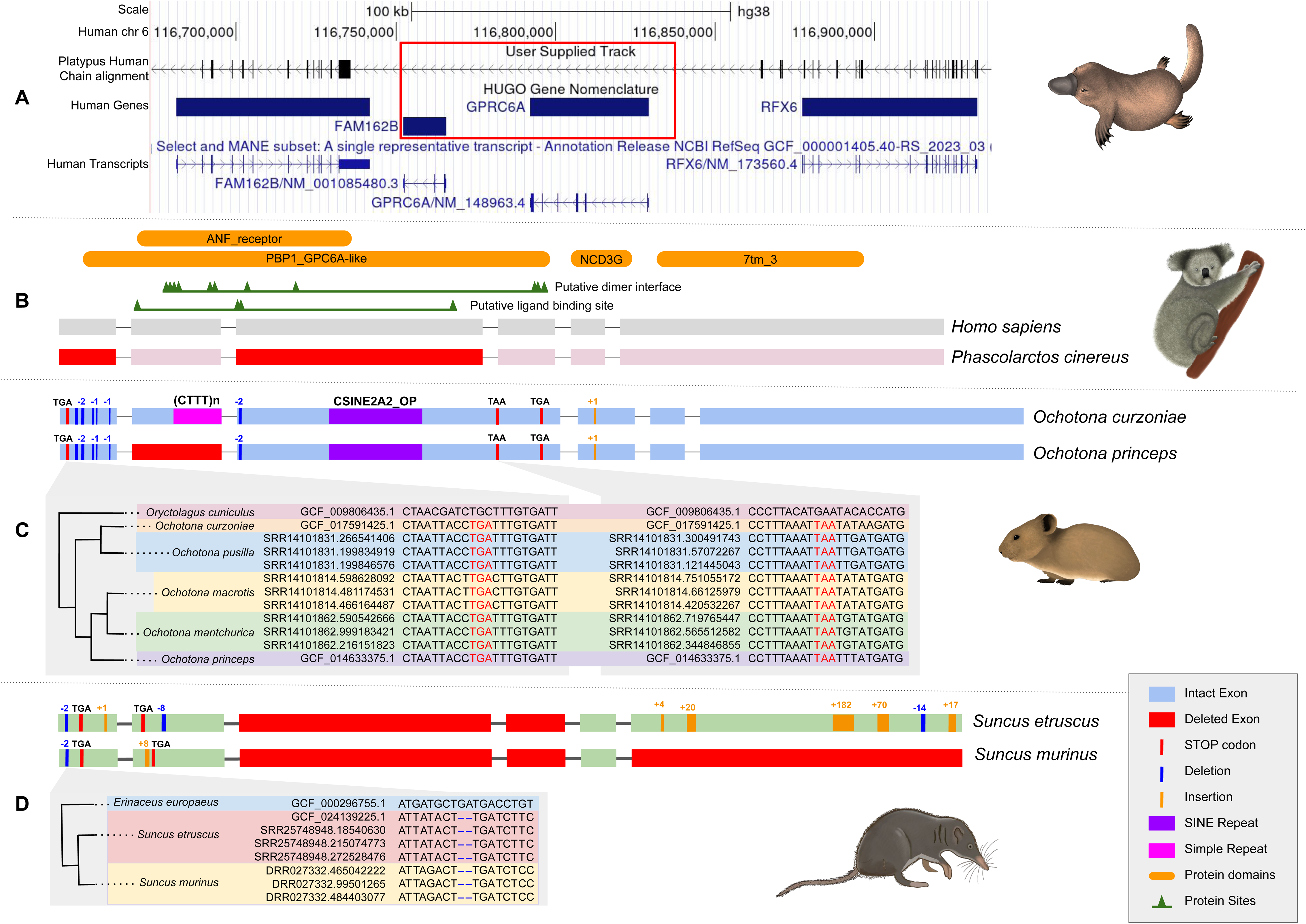
Independent losses of the *GPRC6A* gene. **(A)** Gene loss in *Monotremata* is evident through a genomic alignment of the platypus (*Ornithorhynchus anatinus*) genome (chain file) against the human genome, highlighting the absence of an entire genomic region in a red box. **(B)** The distribution of various domains in the human *GPRC6A* gene, with exons in grey. In contrast, the gene in the koala (*Phascolarctos cinereus*) is depicted in light pink, where red exons indicate those that do not yield BLAST hits and appear to be completely deleted from the genome. **(C)** Loss-of-function events in two *Ochotona* species, with zoomed-in boxes highlighting these events in five *Ochotona* species compared to the rabbit (*Oryctolagus cuniculus*) genome, using short-reads aligned against the rabbit genome. **(D)** illustrates the loss in both *Suncus* species, where a zoomed-in box reveals a two-base pair deletion in the first exon, as identified by aligning short-reads against the European hedgehog (*Erinaceus europaeus*) genome.

While the *GPRC6A* gene in the rabbit (*Oryctolagus cuniculus*) is intact, all four species of pika (Ochotonidae) with sequenced genomes have highly degraded versions of the *GPRC6A* gene at the syntenic locus. In the American pika (*Ochotona princeps)* (Liu et al. 2022), the absence of BLAST hits for the second exon indicates a potential genomic deletion. This deletion was confirmed by aligning PacBio long reads to the American pika genome, ruling out the possibility of assembly artefacts (**Supplementary Figure S95**). The presence of shared gene-disrupting mutations across pika species, including repeat insertions within the coding regions, supports a gene loss event in the common ancestor of pika species (see **Fig. 6C**). Interestingly, the rabbit also showed relaxation in selection pressure (see the last section of Results), suggesting that the gene might be under relaxation in the Lagomorph ancestor, leading to its disintegration in pika.

In the genus *Suncus*, the poor quality of the reference genome limited the reliability of genomic data. However, SRA datasets were available for both the Etruscan shrew (also known as the white-toothed pygmy shrew) (*Suncus etruscus)* and the Asian house shrew (*Suncus murinus*). Our analysis revealed numerous frame-disrupting mutations in both species, particularly in the initial segments of the gene, with pronounced disintegration observed in the Asian house shrew (**Fig. 6D**). Notably, neither exon two nor exon three provided significant hits in either species. In contrast, exon six failed to yield a hit, specifically in the Asian house shew. Validation of the genome using PacBio long-read sequencing supported a gap-free genome at the *GPRC6A* locus (**Supplementary Figure S96**).

Upon further analysis, several species annotated as “low quality” in GenBank were found to contain an intact open reading frame (ORF) and could be reconstructed using short-read datasets. These sequences appeared disintegrated in the NCBI annotation, potentially due to misannotation or assembly errors. After reconstruction, at least one functional isoform was identified in each case (**Supplementary Text 4).**

### Diet and habitat are associated with *GPRC6A* Gene Loss

A strong association between *GPRC6A* gene status and diet/habitat was identified based on a phylogenetically corrected ANOVA (see **Fig. 7A, Supplementary Table S16**). We found that the *GPRC6A* gene loss has happened significantly (p-value: 0.006 of PhyloANOVA) more frequently in fully aquatic species (100% of species with gene loss) compared to terrestrial species (21.15% of species with gene loss [Loss: 33 & Intact: 156]). Subsequent pairwise comparisons among the three diet classes (herbivores, omnivores, carnivores) among terrestrial mammals found a higher frequency of gene loss in herbivores (35.61% of species with gene loss [Loss: 26 & Intact: 47]) than in omnivores (1.85% of species with gene loss [Loss: 1 & Intact: 53]) and carnivores (9.67% of species with gene loss [Loss: 6 & Intact: 56]). The pairwise corrected P-values (method = “holm”) of the post hoc phylANOVA test revealed a significant difference between the Omnivore and Herbivore comparisons (F = 121.77, p-value = 0.006). In contrast, the comparisons between the Carnivore vs Herbivore (p-value=0.28) and the Carnivore vs Omnivore (p-value=0.69) were not statistically significant.

**Fig. 7.**
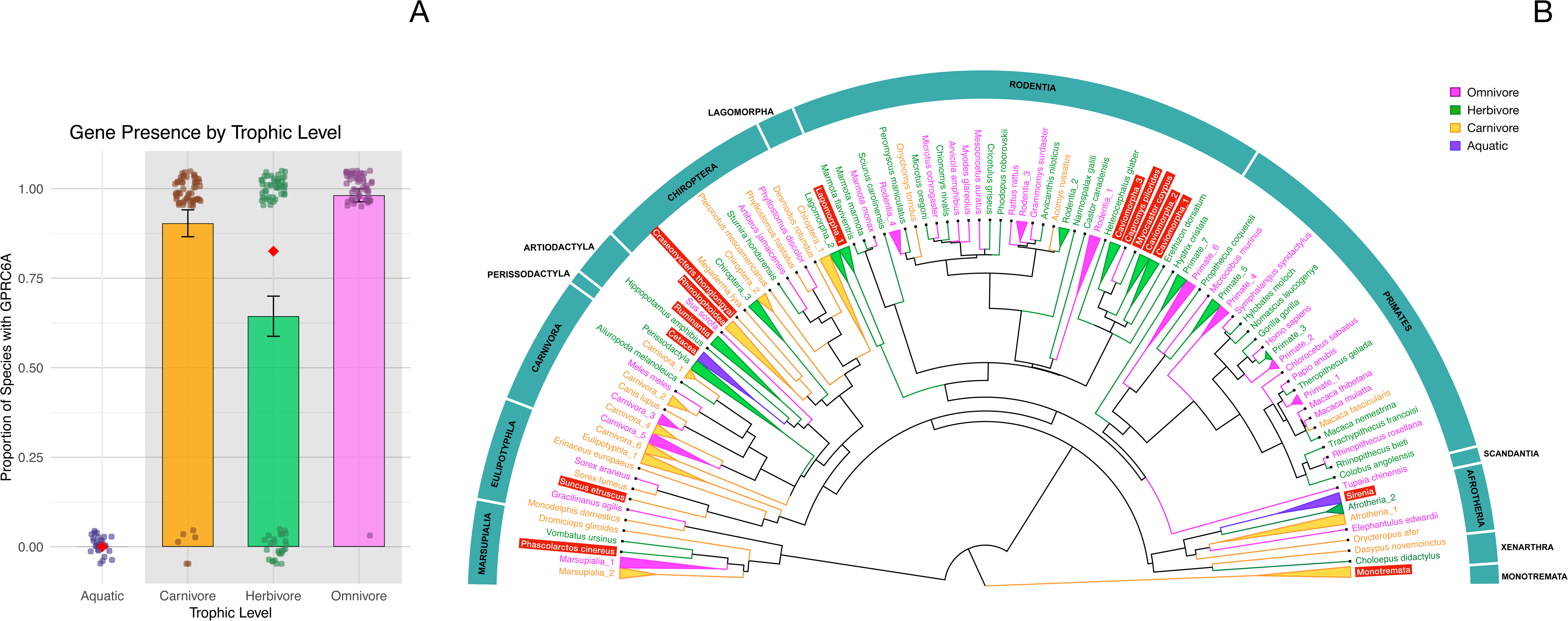
Comparative analysis of the *GPRC6A* gene presence across mammalian groups. **(A)** Bar plots represent the mean proportion of species possessing the *GPRC6A* gene within each category—Aquatic, Carnivore, Herbivore, and Omnivore. Vertical black lines indicate standard error. Individual species data points are represented as overlaid coloured dots: purple for aquatic species, orange for carnivores, green for herbivores, and deep maroon for omnivores. A shaded grey background highlights terrestrial groups (Carnivore, Herbivore, Omnivore). Red diamond markers represent the overall mean gene presence across the Aquatic and Terrestrial categories. (**B)** Phylogenetic tree of representative mammalian species, annotated with habitat-trophic classifications. Tree branches and labels are colored by trophic category: green (Herbivore), magenta (Omnivore), mustard (Carnivore), and purple (Aquatic). Clades are collapsed by shared habitat-trophic group for visual clarity. Red-highlighted labels with white text denote lineages with observed *GPRC6A* gene loss. The tree is downloaded from TimeTree and is visualised in FigTree with manual curation of clade labels.The complete list of species represented within each collapsed clade, including their taxonomic details and gene status, is provided in **Supplementary Table S16**.

The phylogenetic distribution of the gene loss and the associated diet/habitat (see **Fig. 7B**) highlights repeated, independent losses of *GPRC6A* across mammalian clades, with a higher prevalence among herbivorous and aquatic groups. Together, these results demonstrate that *GPRC6A* evolution is tightly linked to ecological niche, with the diet and habitat shaping the likelihood of receptor loss.

### *GPRC6A* is highly similar to the taste receptors *CASR* and *TASRs*

To understand the evolutionary context of *GPRC6A* in relation to other Class C G protein-coupled receptors (GPCRs), we examined its sequence similarity landscape using CLANS analysis of all Class C GPCR protein sequences. The analysis revealed distinct clusters corresponding to major receptor families, each colour-coded and labelled according to representative members (see **Fig. 8**). The clustering pattern reflects pairwise sequence similarities among proteins within the Class C GPCR family. A magnified view of the *GPRC6A-CASR-TAS1R* cluster, highlighted in pink, reveals three subgroups with individual *TAS1R* members shown separately (see **Fig. 8B**). The spatial positioning of the *GPRC6A* cluster underscores its close evolutionary association with calcium-sensing and taste receptor groups. The CLANS results clearly demonstrate the high similarity of *GPRC6A* to *CASR* (the closest annotated paralog in vertebrate species). Notably, the clustering between *GPRC6A* and *CASR* is closer than between *TAS1R1* and *TAS1R3*. This strong similarity suggests that the functional diversification of *GPRC6A* may have co-evolved with chemosensory and nutrient-sensing pathways, providing a molecular link between the ecological patterns of *GPRC6A* loss described above and its role in sensing dietary and environmental cues.

**Fig. 8.**
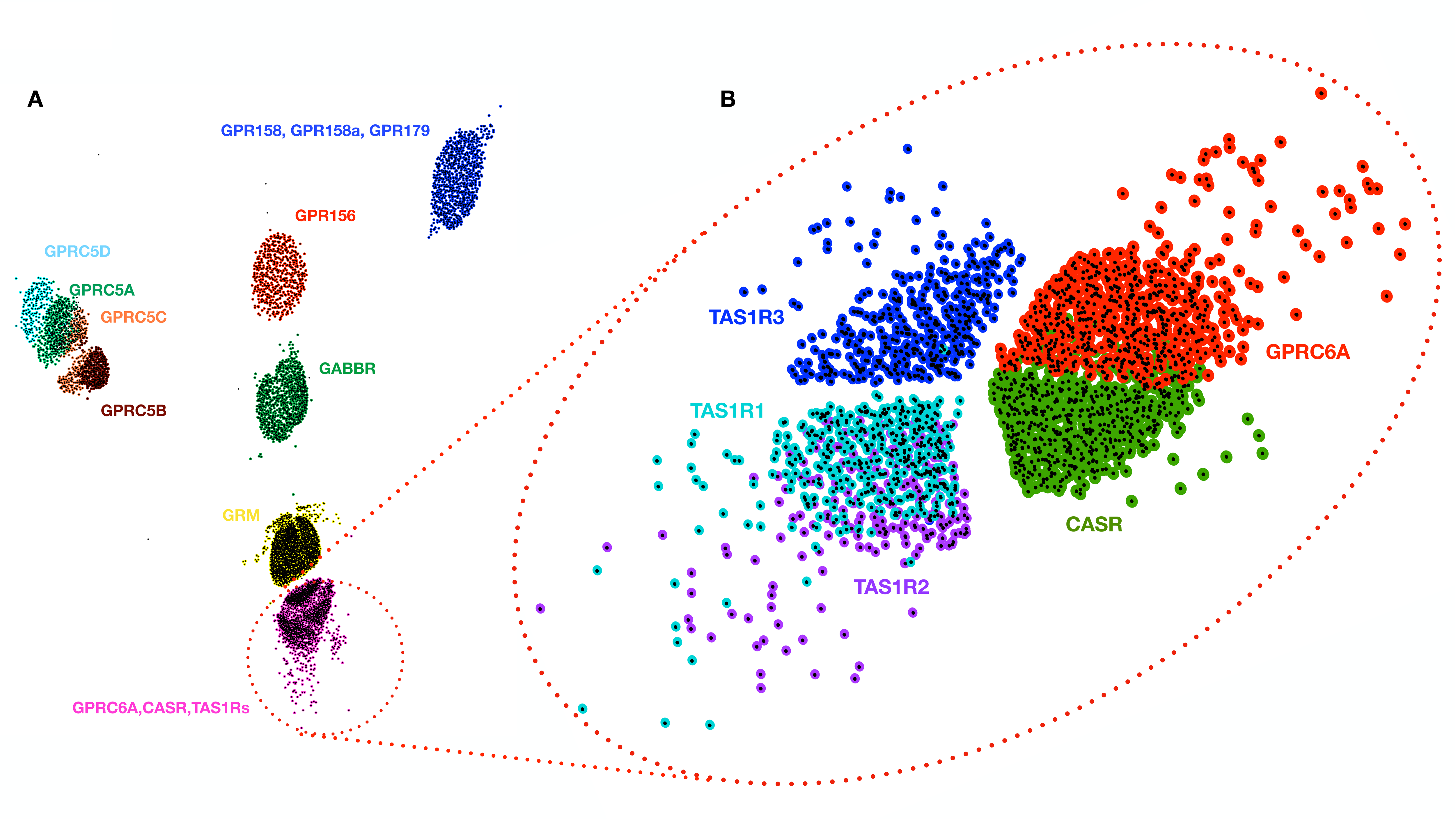
CLANS clustering of Class C GPCR proteins. **(A)** CLANS analysis of all Class C GPCR protein sequences showing distinct clusters corresponding to major receptor families. Each cluster is colour-coded and labelled according to representative members: *GABBR* (green), *GRM* (yellow), *GPR158*/*179* (indigo), *GPR156* (dark red), *GPRC5A* (light green), *GPRC5B* (brown), *GPRC5C* (orange), *GPRC5D* (teal), and the *GPRC6A*–*CASR*–*TAS1R* group (pink). The clustering pattern reflects pairwise sequence similarities among proteins within the Class C GPCR family. **(B)** A magnified view of the *GPRC6A*–*CASR*–*TAS1R* cluster is highlighted in pink in panel A. The zoomed-in view reveals three subgroups—*GPRC6A* (red), *CASR* (green), and *TAS1Rs*—with individual *TAS1R* members shown separately. The spatial positioning of the *GPRC6A* cluster underscores its close evolutionary association with calcium-sensing and taste receptor groups.

### Lineage-Specific Variation in Selective Constraints on *GPRC6A*

Comparative selection analyses revealed that *GPRC6A* is predominantly subject to strong purifying selection across mammals, consistent with functional constraint on its core physiological roles. Results from HyPhy RELAX, codeml branch models, and aBSREL tests showed minimal evidence for widespread relaxation or episodic selection, with the majority of taxa exhibiting ω values indicative of conserved evolutionary pressure. A subset of species demonstrated consistent lineage-specific variation in selection intensity, including intensified purifying selection in *Myodes glareolus*, *Urocitellus parryii*, *Phyllostomus hastatus*, *Neogale vison*, and *Homo sapiens*, and relaxed constraint in *Felis catus*, *Heterocephalus glaber*, and *Oryctolagus cuniculus*. These results were robust to tree topology, with only a few taxa showing topology-dependent outcomes between species- and gene-tree analyses. A detailed, species-specific breakdown of these results is provided in **Supplementary Tables S17-S21 (Supplementary Text 4**). Significant gBGC tracts were detected in several mammalian lineages that also exhibited signatures of selection. However, in most species, these tracts were relatively short (typically 109–180 bp), suggesting that they are unlikely to have substantially influenced the selective patterns inferred for *GPRC6A*. Examples include *Molossus molossus, Gorilla gorilla, Arvicanthis niloticus,* and *Heterocephalus glaber*, where the observed shifts in selection are more plausibly genuine rather than artefacts of biased nucleotide substitution. In contrast, *Otolemur garnettii* displayed a markedly longer tract (∼941 bp), which may have contributed to the signal of intensified purifying selection in this species by altering local GC content and codon usage. Overall, while gBGC may shape *GPRC6A* evolution in specific cases, its broader influence across mammals appears limited and lineage-specific. (**Supplementary Figures S97 – S107**).

## Discussion

Despite considerable research efforts over the past two decades, the functional roles of *GPRC6A* remain a topic of ongoing debate, primarily due to its interactions with multiple ligands, its proposed involvement in metabolic regulation (Pi et al. 2017), and kokumi taste perception (Yamamoto et al. 2025). Understanding the role of *GPRC6A* is vital for elucidating its impact on metabolic health; however, gene-disrupting polymorphisms in the human *GPRC6A* gene and inconsistent results from knockout studies suggest that its role may be context-dependent or limited (Pi et al. 2020, 2021). Our comparative study across approximately 230 mammalian species reveals a broader pattern of *GPRC6A* gene loss than previously recognised, extending beyond toothed whales to include ruminants, indicating significant changes in evolutionary pressures that may influence its functional relevance. We identified independent gene-disrupting changes in various species, including pika, koalas, and shrews, as well as widespread losses in Rhinolophid bats and many Hystricomorphs, particularly caviomorph rodents, suggesting a link between dietary specialisation and gene dispensability.

The evolutionary loss of the *GPRC6A* gene in various mammalian species appears to be a response to specialised diets and ecological niches, influencing metabolic regulation and taste perception. Our CLANS clustering results provide a structural and evolutionary explanation for this pattern. *GPRC6A* clusters closely with CaSR and *TAS1R* receptors, forming a distinct subfamily within Class C GPCRs (Liu et al. 2016). Both *GPRC6A* and *CaSR* share key structural motifs, and both are expressed in taste tissues, particularly in the anterior tongue and taste buds (Bystrova et al. 2010). CaSR detects glutathione and γ-glutamyl peptides associated with kokumi taste—an attribute that increases the perception of richness or savoury depth (Kuroda and Miyamura 2015)—while *GPRC6A* responds to basic amino acids, including ornithine, which is enriched in protein-rich foods (Yamamoto et al. 2025). These two paralogs therefore appear to act within related nutrient-sensing pathways, helping animals evaluate the nutritional quality of their food, especially in terms of amino acid content. Evidence from carnivorous species, such as domestic cats, shows that *CaSR* contributes to kokumi perception in taste tissue (Wolsan and Sato 2022), while expression studies in rodents indicate that *GPRC6A* is active in distinct taste cell subpopulations associated with amino acid sensing (Bystrova et al. 2010). Together, these observations support the idea that *GPRC6A* functions as a taste receptor, despite its historical focus on metabolic contexts. Our interpretation aligns well with known patterns of taste gene evolution.

*TAS1R* receptors also belong to GPCR Class C and are involved in umami taste perception, similar to *GPRC6A* (Pallante et al. 2021). The loss of the *TAS1R* taste receptor in the giant panda, following its shift to a herbivorous diet, underscores how dietary changes can drive the reduction of genes associated with taste perception and nutrient sensing (Zhao et al. 2010). As herbivores transitioned to high-fibre diets, the necessity for *GPRC6A*, which aids in amino acid metabolism and kokumi taste perception in carnivorous diets, might have also diminished. Relaxed selection further supports this trend, as evidenced by our results in species such as wild boar and rabbit, which share evolutionary ties with ruminants and pikas. Moreover, the observed loss of *GPRC6A* in monotremes appears consistent with the significant reduction of *TAS2R*s and other chemosensory receptors (Zhou et al. 2021), underscoring the idea that dietary adaptations significantly influence the evolution of taste receptors and nutrient-sensing genes across various mammalian lineages. Similar to *TAS1Rs*, *GPRC6A* serves as a molecular bridge between nutrient detection and metabolic regulation, suggesting that both receptors have been shaped by convergent evolutionary pressures linking diet, taste, and physiology.

Losses of the vertebrate *TAS1R* sweet and umami taste receptor genes have occurred repeatedly across lineages, primarily driven by dietary specialisation and relaxed selection on gustation. Comparative genomic studies show that when sweetness or umami detection ceases to be advantageous, such as in obligate carnivores, aquatic feeders, or suction-feeding whales, mutations accumulate, leading to pseudogenisation of *TAS1R*s (Jiang et al. 2012; Feng and Zhao 2013; Feng et al. 2014; Li and Zhang 2014; Liu et al. 2016; Antinucci and Risso 2017; Wolsan and Sato 2022). The heterodimeric structure of these receptors (*TAS1R1* + *TAS1R3* for umami, *TAS1R2* + *TAS1R3* for sweet) makes them especially vulnerable, as inactivation of one subunit can abolish multiple taste modalities. However, extra-oral and pleiotropic functions of *TAS1Rs* in the gut and pancreas may preserve some copies under purifying selection, leading to mosaic patterns of retention and loss. Overall, these studies converge on a model where feeding ecology, molecular interdependence, and secondary functions shape the complex, lineage-specific evolutionary history of *TAS1R* genes. This parallel between *TAS1Rs* and *GPRC6A* highlights a shared evolutionary principle, where shifting nutritional demands and sensory relevance jointly dictate the retention or loss of receptors mediating amino acid and nutrient sensing.

Despite instances of gene loss in specific mammalian clades, our selection analyses reveal that *GPRC6A* has remained mainly under strong purifying selection across most lineages, suggesting that its physiological roles continue to impose substantial functional constraints. The absence of widespread relaxed selection or episodic positive selection indicates that *GPRC6A* retains an essential metabolic or signalling role wherever it is preserved. However, lineage-specific deviations—such as intensified constraint in humans, voles, and certain bats, and relaxed selection in species like cats, rabbits, and naked mole rats—suggest that the evolutionary pressures acting on *GPRC6A* are context-dependent, potentially shaped by differences in diet, thermoregulation, or metabolic ecology. These findings support a model in which gene loss is not a consequence of widespread functional decay, but rather a selective fine-tuning process that reflects shifts in metabolic priorities or ecological specialisation. Thus, while *GPRC6A* remains indispensable in many mammals, its loss in others likely represents adaptive streamlining rather than stochastic degeneration. This adaptive streamlining of *GPRC6A* mirrors the evolutionary trajectory of *TAS1R* receptors, where dietary specialisation and shifting nutrient demands similarly dictate gene retention or loss across mammalian lineages.

The investigation into the loss of the *GPRC6A* gene across various species provides valuable insights into the evolutionary mechanisms that enable different organisms to thrive in specific environmental contexts. Our findings indicate that the absence of *GPRC6A* may confer adaptive advantages in certain ecological niches or dietary habits, while in other cases, environmental pressures might render the gene dispensable, thereby illustrating how such pressures shape genetic evolution. Furthermore, these insights could inform conservation efforts by demonstrating how genetic adaptations contribute to the resilience and survival of species in changing environments. Recent insights into the role of *GPRC6A* in regulating ILC3 cells and IL-22 production have highlighted its therapeutic potential for enhancing immune function and tissue repair in inflammatory conditions (Hou et al. 2022). Studying the loss of *GPRC6A* can reveal new treatment approaches and offer new therapeutic avenues for colitis and related gastrointestinal issues. Overall, our analysis highlights the complexity and flexibility of genetic regulation, as well as its role in shaping the physiological and ecological diversity observed across Mammalia.

## Conclusion

Our findings highlight the loss of *GPRC6A* across diverse mammalian lineages, revealing a pattern that aligns closely with specialised ecological and dietary adaptations. This gene loss appears to confer specific advantages within unique metabolic and environmental contexts, such as energy conservation in aquatic mammals, high metabolic turnover in insectivores, and reliance on microbial fermentation in herbivores. The role of *GPRC6A* as a nutrient-sensing receptor within complex metabolic and immune pathways underscores its potential as a therapeutic target for metabolic and inflammatory disorders. Our comparative genomic analysis contributes to understanding how evolutionary pressures can lead to gene dispensability, shaping genetic diversity and metabolic regulation across mammalian species.

## Materials and methods

### Synteny and orthology assessment

Identifying 1-to-1 orthologs of genes from multigene families relies upon conserved synteny in the flanking regions of the focal gene. Hence, we used the NCBI genome data viewer and Ensembl synteny view to compare the synteny using three flanking genes upstream and downstream of the *GPRC6A* gene. Approximately 450 species representing all extant classes of the Infraphylum jawed vertebrates (Gnathostomata) except Monotremata have annotations for *GPRC6A* orthologs in the NCBI database (**Supplementary Table S1**). We compared the synteny in representative species with well-annotated, high-quality genomes spanning the mammalian phylogeny to evaluate changes in gene order. Lack of annotation in the genomic regions flanking the *GPRC6A* gene was also investigated to either perform annotation of intact genes or identify gene remnants using a blastn search of the genome assemblies using the intact human gene as the query **(Supplementary Table S2)**. The *GPRC6A* orthologs annotated at the syntenic locations were further evaluated for completeness by performing multiple sequence alignments of the ORFs using Guidance (v2.02) with MAFFT as the aligner (Sela et al. 2015).

### Evaluation of sequence homology relationships among class C GPCRs

The sequence homology-based reconstruction of the protein evolutionary history is a reasonably reliable predictor of function (Loewenstein et al. 2009). Hence, the evaluation of sequence homology among the class C GPCRs was performed. All the protein sequences of annotated class C GPCRs orthologs (*CASR*, *GABBR1*, *GABBR2*, *GPR156*, *GPR158*, *GPR179*, *GPRC5A*, *GPRC5B*, *GPRC5C*, *GPRC5D*, *GPRC6A*, *GRM1*, *GRM3*, *GRM4*, *GRM5*, *GRM6*, *GRM7*, *GRM8*, *TAS1R1*, *TAS1R2*, *TAS1R3*) were obtained from the NCBI database. The longest protein isoform was selected from each species. Using the CLANS program, we clustered the proteins based on their all-against-all pairwise sequence similarities (Frickey and Lupas 2004) with default settings. The CLANS output was visualised in the CLANS GUI and manually annotated based on the most abundant protein in that cluster. The clusters contained well-annotated proteins from the same orthology group.

### Identification and corroboration of putative gene loss events

We considered all mammalian species labelled “LOW QUALITY” in GenBank as potential candidates for gene loss (**Supplementary Table S3**). Species with incomplete sequences and those that failed to align with the human *GPRC6A* gene were aligned with the phylogenetically closest high-quality reference genomes to scrutinise and re-annotate to get complete open reading frames. Roughly 13% of the annotated sequences labelled “LOW QUALITY” had discrepancies that could be attributed to genome assembly or annotation errors and were reconstructed (**Supplementary Table S4**). We employed a previously published five-pass strategy to validate genuine gene loss events in each candidate species (Sharma et al. 2020, 2022; Shinde et al. 2021). Moreover, we utilised the TOGA pipeline (Kirilenko et al., 2023) as an additional step to identify gene loss events and assess splice-site mutations. The Chain alignments needed for TOGA, as well as the detection of deletions, were generated using the “make_lastz_chains” pipeline available at https://github.com/hillerlab/make_lastz_chains. We employed either a human chromosome or the chromosome containing the focal gene in the phylogenetically closest species with a genome assembly as a reference for this pipeline.

Gene loss could be ascertained in several closely related species of ruminants and cetaceans using genomic datasets (**Supplementary Table S5**). We subsequently sought shared gene-disrupting changes to parsimoniously explain the events as part of the five-pass gene loss validation strategy (Sharma et al. 2020; Shinde et al. 2021). The presence of the same gene loss causing changes in closely related species suggests a single shared incident in the common ancestor of the species that share the change. Hence, we could map the chronology of gene-disrupting events onto the phylogenetic tree downloaded from the TimeTree5 website (Kumar et al. 2022) to reconstruct the history of gene loss parsimoniously. We have defined gene loss/pseudogenisation based only on the presence of gene-disrupting changes within the coding regions. The *GPRC6A* gene expression is highly variable in the RNA-seq data and cannot be conclusive evidence of loss. Moreover, gene expression at the RNA or protein level is not a reliable indicator for pseudogenes and therefore does not serve as a criterion for defining gene loss (Xu and Zhang 2015).

Identifying protein domains and features, including binding sites, was conducted through the CD-search interface, utilising the National Library of Medicine’s Conserved Domain Database (CDD) (Lu et al. 2020). The amino acid sequence of *GPRC6A* served as the input query for the CD-search, employing the default settings and hits meeting the “specific” classification criteria were carefully selected. We examined functional domains that coincide with the deleted or disrupted exons to assess the consequences of the exon deletions and repeat insertions. RepeatMasker (Smit et al. 1996) was used to identify and classify repeats in these regions.

### Sequencing-based validation of gene-disruption

Errors in genome assembly can manifest as gene-disruptive events due to incorrect nucleotide sequences or inadequate long-range assembly contiguity. For instance, incorporating the erroneous nucleotide bases in the genome assembly can masquerade as premature stop codons. Such base-pair level errors in the genomic assembly sequence were rectified by searching short-read sequence databases (for example, using Illumina sequencing technology) to identify the nucleotide base supported by the sequencing reads. The blastn tool was used to search the short-read archive databases, and the gene-disrupting changes were verified by manual inspection of the blast hits. The intact *GPRC6A* gene of the phylogenetically closely related species was used as the query sequence in addition to that of the focal species as an additional validation step (**Supplementary Table S6-13**). Long-range errors in the assembly contiguity can result in the erroneous ordering of the exons, missing or incomplete exons, and artifactual translocation of part or the entirety of the gene to a non-syntenic location. Long-read length technologies such as PacBio and Oxford Nanopore can identify and correct genome assembly errors spanning several thousand nucleotides. We mapped the long reads to the genome assembly using Minimap2 (Li 2018) and visualised the resulting alignments in the UCSC genome browser. Reads longer than 50Kb spanning the genomic region encompassing the putative location of *GPRC6A* and flanking genes on both sides were used to validate the genome assembly.

Genes with extreme GC content tend to be missing from genome assemblies and short-read datasets. To mitigate potential complications associated with high GC content, we extracted the coding sequence of the longest isoforms of all annotated mammals available from NCBI. We evaluated the prevalence of extreme GC regions by comparing the mean GC content and the fraction of high GC stretches in each sequence (**Supplementary Table S14)**. However, in the case of Caviomorpha rodents, for which long-read sequencing data were unavailable, we evaluated GC content by analysing 100 bp windows using bedtools windows and nuc commands (**Supplementary Table S15**).

### Association of diet/habitat with the *GPRC6A* gene loss

We compiled species-level diet information and gene status data to assess the relationship between diet and the presence of the *GPRC6A* gene. The diet preferences of terrestrial mammals were obtained from an earlier study (Kissling et al. 2014). Since this dataset is for terrestrial mammals, we grouped them separately based on their habitat (aquatic or terrestrial) to include fully aquatic mammals. Based on the curated dietary dataset, terrestrial species were grouped by diet (Carnivore, Herbivore, Omnivore). Gene status was coded as binary (1 = gene present, 0 = gene absent) from genomic screening results (**Supplementary Table S16**). Phylogenetically informed ANOVA (phylANOVA; (Ives and Garland 2010)) was performed in phytools to test for differences in *GPRC6A* gene presence between groups.

### Inference of selection signatures

The longest isoform from each gene was utilised to generate clade-specific multiple sequence alignments using Guidance2 (Sela et al. 2015) with the codon option, and MAFFT(v7.407) (Katoh et al. 2002) as the alignment tool. To ensure that substitution saturation did not obscure true dN/dS estimates, sequence saturation was assessed in DAMBE v7 (Xia 2013) by calculating the index of substitution saturation (Xia et al. 2003) for each codon position and comparing it with the critical Iss value (Iss.c). These multiple sequence alignments were used in conjunction with time-calibrated species trees downloaded from the TimeTree5 (Kumar et al. 2022) website for selection tests. We used the RELAX (Wertheim et al. 2015) program from the HyPhy package (Kosakovsky Pond et al. 2005) to detect signatures of relaxation or intensification of selection in each species (**Supplementary Table S17-18**). To obtain direct ω estimates and confirm patterns detected by RELAX, branch model analyses (model = 2) were conducted in codeml (PAML v4.9 (Yang 2007)) for each clade to test for lineage-specific variation in selective pressure (**Supplementary Table S20-21**). Using likelihood ratio tests, the two-ratio model (separate ω values for foreground and background) was compared to the one-ratio null model (model = 0, a single ω across all branches) (Goldman and Yang 1994; Yang 1998). Positive selection at individual sites (**Supplementary Table S19**) along each lineage was identified using ABSREL (Smith et al. 2015). To complement selection analyses, GC-biased gene conversion (gBGC) was evaluated. Neutral substitution models were estimated using phyloFit, and phastBias from the PHAST package (Siepel et al. 2005; Hubisz et al. 2011) was applied to identify putative gBGC tracts across the alignments.

## Data availability

The associated data is available in an easy-to-view format on the GitHub repository: https://github.com/saumyagupta09/GPRC6A.

## Acknowledgements

We thank the Ministry of Human Resource Development for the fellowship to S.G. The Department of Biotechnology, Ministry of Science and Technology, India (Grant no. BT/11/IYBA/2018/03) and Science and Engineering Research Board (Grant no. ECR/2017/001430) provided funds for procuring computational resources (i.e., Har Gobind Khorana Computational Biology cluster) used.

## Author contributions

S.G., A.B.P., and N.V. devised the project, including the main conceptual ideas and proof outline. S.G. wrote the manuscript with inputs from N.V. Initial collection and analysis were done by A.B.P. and verified by S.G. Preliminary results for key findings were made by A.B.P. Main results and figures were made by S.G. with inputs from A.B.P. A.S. performed additional analysis and verified the results. All authors reviewed the manuscript.

## Competing interests

The authors declare that they have no competing financial interests.

## Notes

https://github.com/saumyagupta09/GPRC6A.git

### Competing Interest Statement

The authors have declared no competing interest.

