## Supplementary Figures for "Master of none: *GPRC6A* gene loss is more widespread than previously known"

S1

*Homo sapiens*

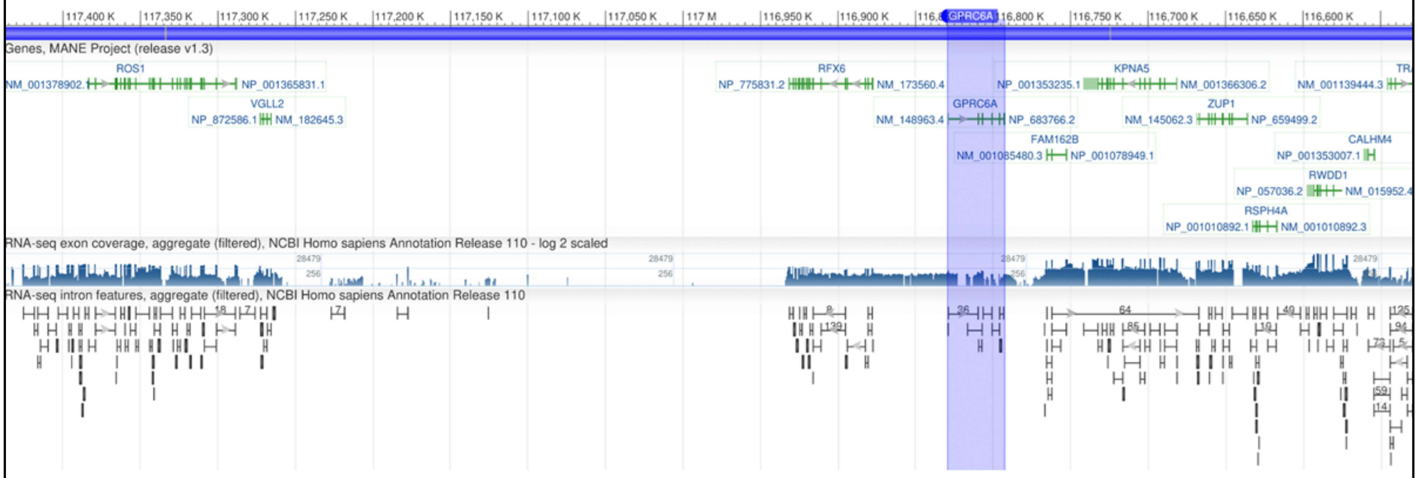


**Supplementary Figure S1. Synteny of the *GPRC6A* region in *Homo sapiens*:** This figure illustrates the synteny of the genomic region containing the *GPRC6A* gene in *Homo sapiens*, as visualized using the NCBI Genome Data Viewer. The highlighted region specifically marks the *GPRC6A* gene and its flanking genes, providing a detailed view of the genomic context in which *GPRC6A* resides. This figure highlights that the genomic region around *GPRC6A* is conserved.

S2

*Rattus norvegicus*

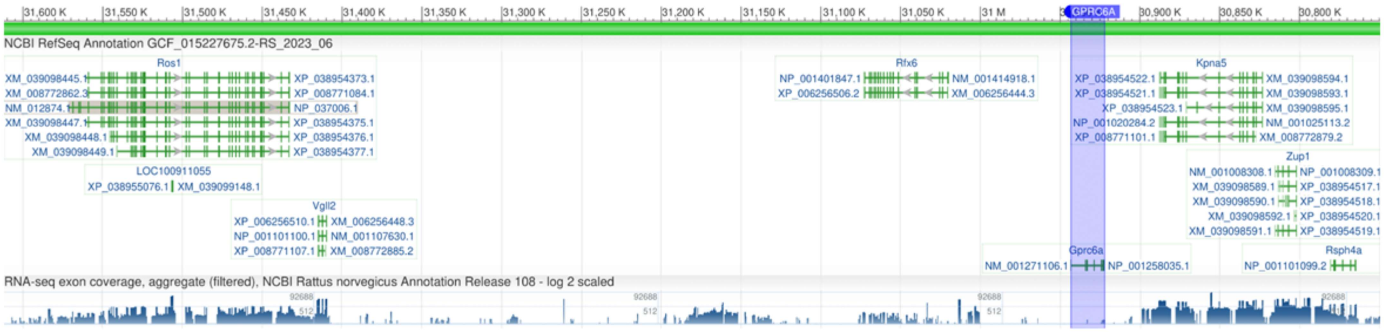


**Supplementary Figure S2. Synteny of the *GPRC6A* Region in *Rattus norvegicus***: This figure illustrates the synteny of the genomic region containing the *GPRC6A* gene in *Rattus norvegicus*, as visualized using the NCBI Genome Data Viewer. The highlighted region specifically marks the *GPRC6A* gene and its flanking genes, providing a detailed view of the genomic context in which *GPRC6A* resides. This figure highlights that the genomic region around *GPRC6A* is conserved.

S3

*Erinaceus europaeus*

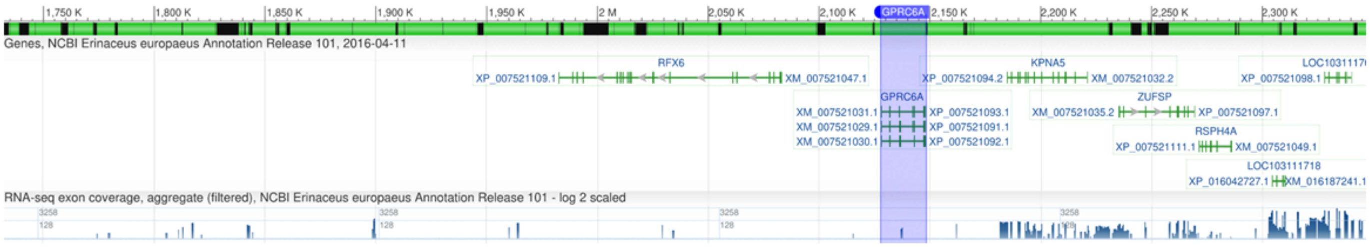


**Supplementary Figure S3. Synteny of the *GPRC6A* Region in *Erinaceus europaeus*:** This figure illustrates the synteny of the genomic region containing the *GPRC6A* gene in *Erinaceus europaeus*, as visualized using the NCBI Genome Data Viewer. The highlighted region specifically marks the *GPRC6A* gene and its flanking genes, providing a detailed view of the genomic context in which *GPRC6A* resides. This figure highlights that the genomic region around *GPRC6A* is conserved.

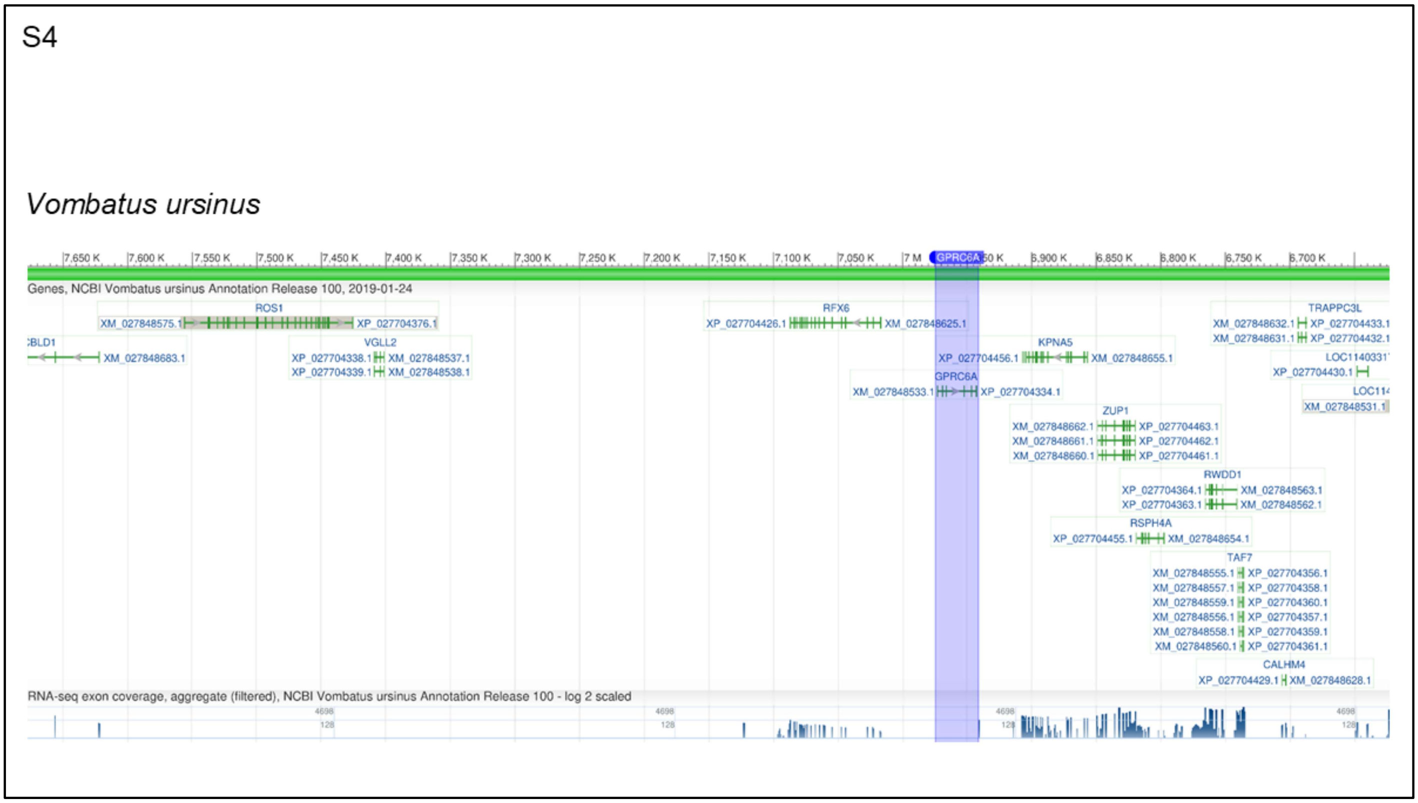


**Supplementary Figure S4. Synteny of the *GPRC6A* region in *Vombatus ursinus*:** This figure illustrates the synteny of the genomic region containing the *GPRC6A* gene in *Vombatus ursinus*, as visualized using the NCBI Genome Data Viewer. The highlighted region specifically marks the *GPRC6A* gene and its flanking genes, providing a detailed view of the genomic context in which *GPRC6A* resides. This figure highlights that the genomic region around *GPRC6A* is conserved.

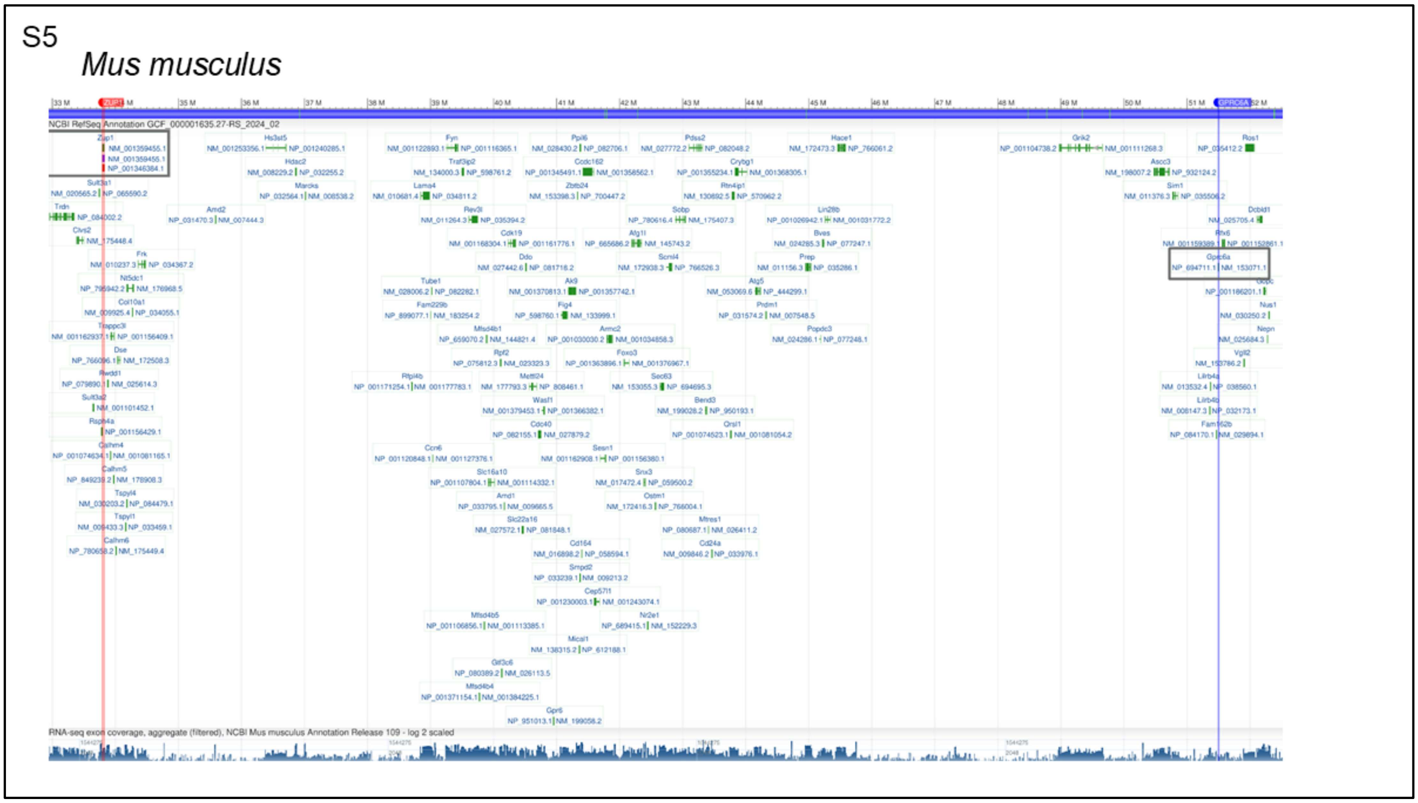


**Supplementary Figure S5.Genomic organization between *GPRC6A* and ZUP1 locus in *Mus musculus*:** The figure shows the genomic region between *ZUP1* and *GPRC6A* genes as visualized in the NCBI Genome Data Viewer. *ZUP1* is located approximately 19 Mb upstream of *GPRC6A*, with the ZUP1 highlighted in red while GPRC6A highlighted in blue. Notably, *KPNA5* is absent in *M. musculus*, whereas *ZUP1* remains present. This arrangement suggests that *GPRC6A* resides within an evolutionary breakpoint region (EBR) in rodents, where frequent local genomic shuffling has occurred.

S6

*Mus musculus*

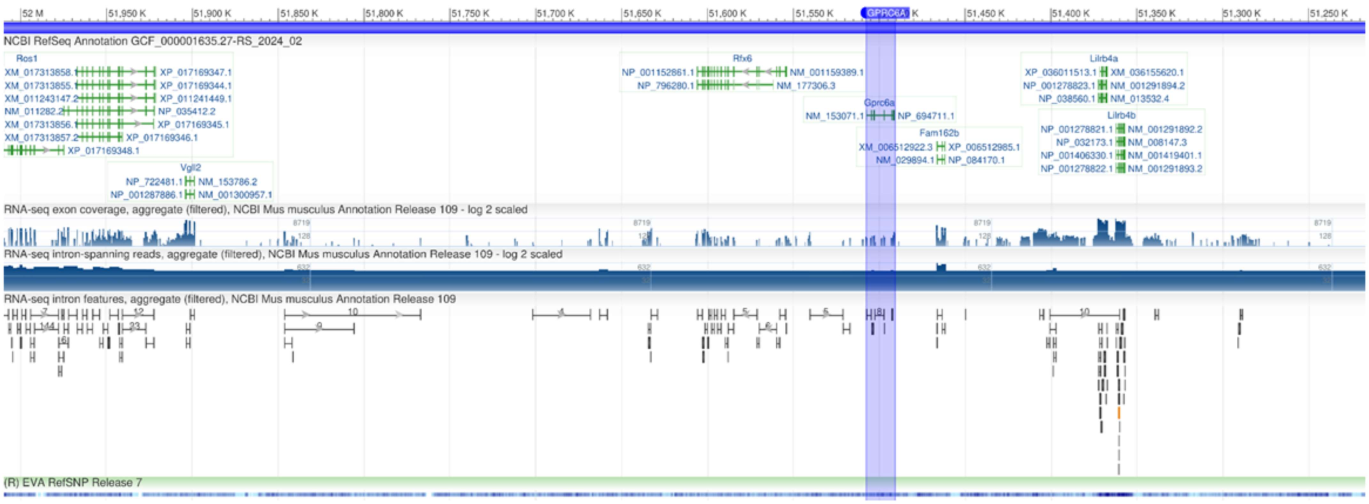


**Supplementary Figure S6. Synteny of the *GPRC6A* Region in *Mus musculus*:** This figure illustrates the synteny of the genomic region containing the *GPRC6A* gene in *Mus musculus*, as visualized using the NCBI Genome Data Viewer. The highlighted region specifically marks the *GPRC6A* gene and its flanking genes, providing a detailed view of the genomic context in which *GPRC6A* resides. This figure highlights that the genomic region around *GPRC6A* might fall in an EBR region.

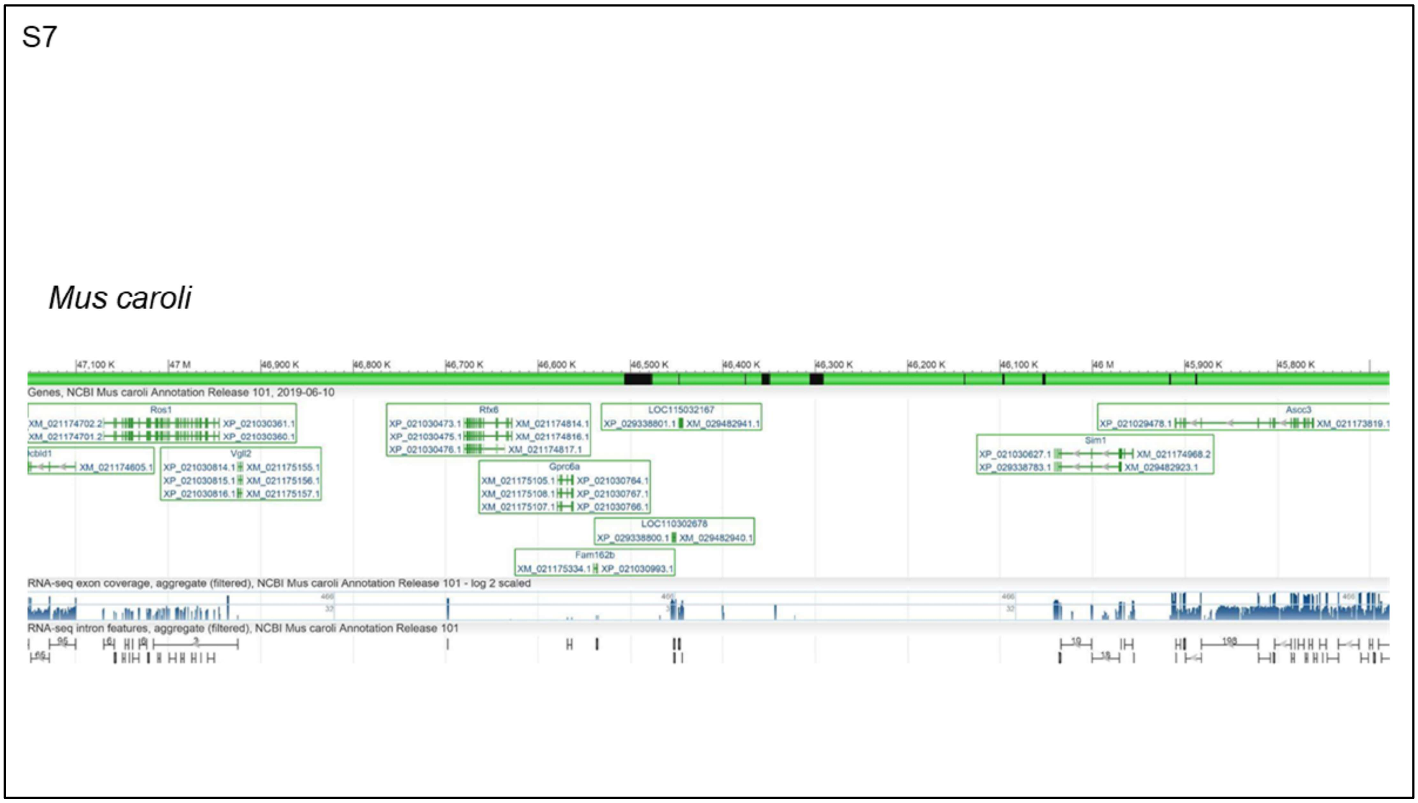


**Supplementary Figure S7:Synteny of the *GPRC6A* Region in *Mus caroli*:** This figure illustrates the synteny of the genomic region containing the *GPRC6A* gene in *Mus caroli*, as visualized using the NCBI Genome Data Viewer. The highlighted region specifically marks the *GPRC6A* gene and its flanking genes, providing a detailed view of the genomic context in which *GPRC6A* resides. This figure highlights that the genomic region around *GPRC6A* might fall in an EBR region.

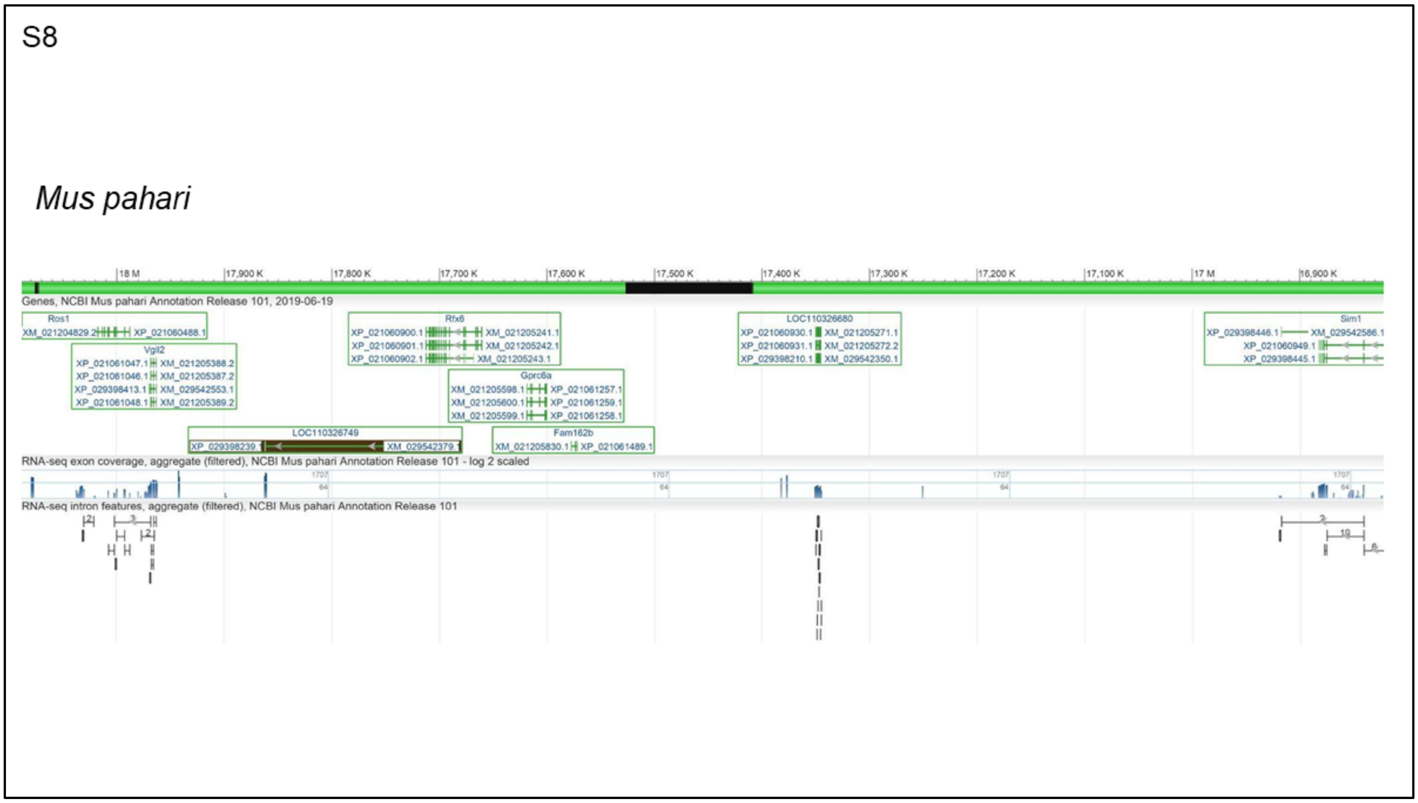


**Supplementary Figure S8:Synteny of the *GPRC6A* Region in *Mus pahari*:** This figure illustrates the synteny of the genomic region containing the *GPRC6A* gene in *Mus pahari*, as visualized using the NCBI Genome Data Viewer. The highlighted region specifically marks the *GPRC6A* gene and its flanking genes, providing a detailed view of the genomic context in which *GPRC6A* resides. This figure highlights that the genomic region around *GPRC6A* might fall in an EBR region.

S9

*Tachyglossus aculeatus*

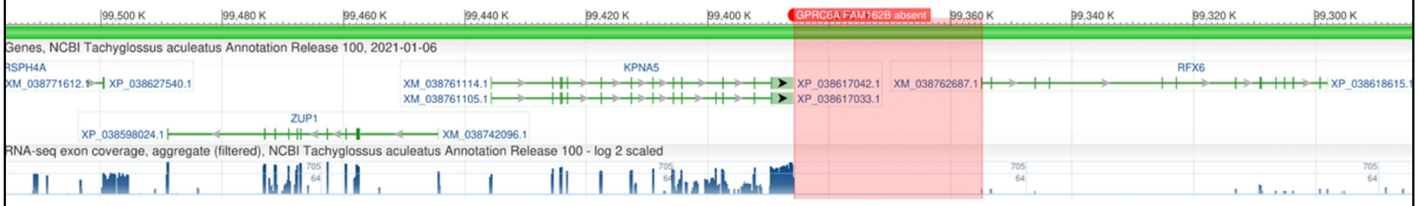


**Supplementary Figure S9. Synteny of the *GPRC6A* region in *Tachyglossus aculeatus***: This figure illustrates the synteny of the genomic region containing the *GPRC6A* gene in *Tachyglossus aculeatus*, as visualized using the NCBI Genome Data Viewer. The highlighted region specifically marks the genomic region of the putative location of the *GPRC6A* gene and its flanking genes, providing a detailed view of the genomic context in which *GPRC6A* resides. This figure highlights that the genomic region containing *GPRC6A* and its flanking genes are deleted from the genome.

S10

*Ornithorhynchus anatinus*

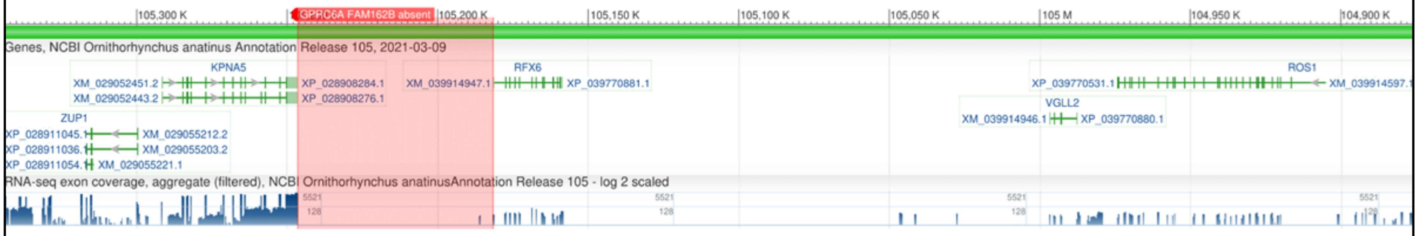


**Supplementary Figure S10. Synteny of the *GPRC6A* region in *Ornithorhynchus anatinus*:** This figure illustrates the synteny of the genomic region containing the *GPRC6A* gene in *Ornithorhynchus anatinus*, as visualized using the NCBI Genome Data Viewer. The highlighted region specifically marks the genomic region of putative location of *GPRC6A* gene and its flanking genes, providing a detailed view of the genomic context in which *GPRC6A* resides. This figure highlights that the genomic region containing *GPRC6A* and flanking genes are deleted from the genome.

S11

*Gallus gallus*

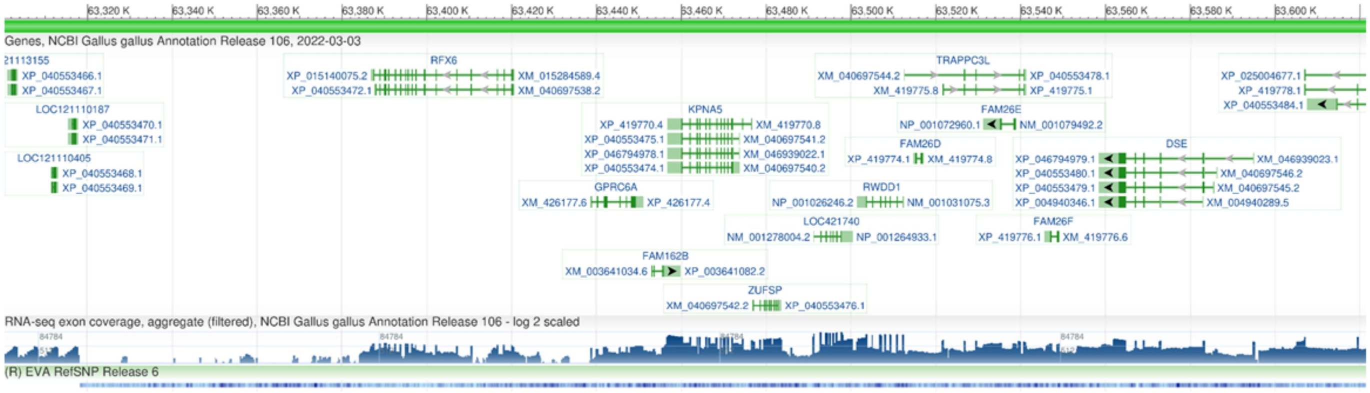


**Supplementary Figure S11. Synteny of the *GPRC6A* region in *Gallus gallus*:** This figure illustrates that the synteny of the genomic region containing the *GPRC6A* gene is conserved in outgroups such as birds, with *Gallus gallus* serving as the representative species. The highlighted region specifically marks the *GPRC6A* gene and its flanking genes, providing a detailed view of the genomic context in which *GPRC6A* resides. This figure underscores that the genomic region around *GPRC6A* is conserved across clades beyond Mammalia.

S12

*Rana temporaria*

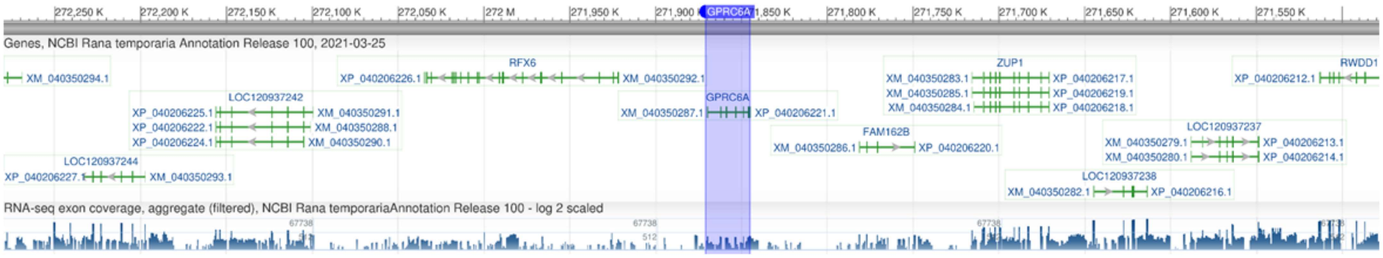


**Supplementary Figure S12. Synteny of *GPRC6A* region in *Rana temporaria*:** This figure demonstrates that the synteny of the genomic region containing the *GPRC6A* gene is conserved across diverse vertebrate lineages, including amphibians, with *Rana temporaria* as the representative species. The highlighted region specifically marks the *GPRC6A* gene and its flanking genes, providing a detailed view of the genomic context in which *GPRC6A* resides.

S13

*Hippopotamus amphibius*

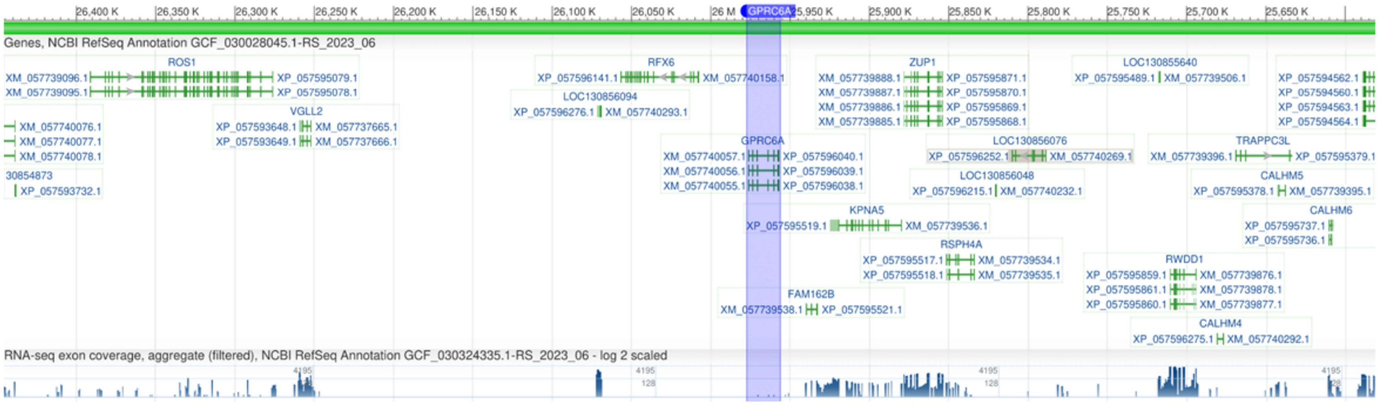


**Supplementary Figure S13. Synteny of the GPRC6A Region in *Hippopotamus amphibius*:** This figure illustrates the synteny of the genomic region containing the *GPRC6A* gene in *Hippopotamus amphibius*, as visualized using the NCBI Genome Data Viewer. The highlighted region specifically marks the *GPRC6A* gene and its flanking genes, providing a detailed view of the genomic context in which *GPRC6A* resides. This figure highlights that the genomic region around *GPRC6A* is conserved.

S14

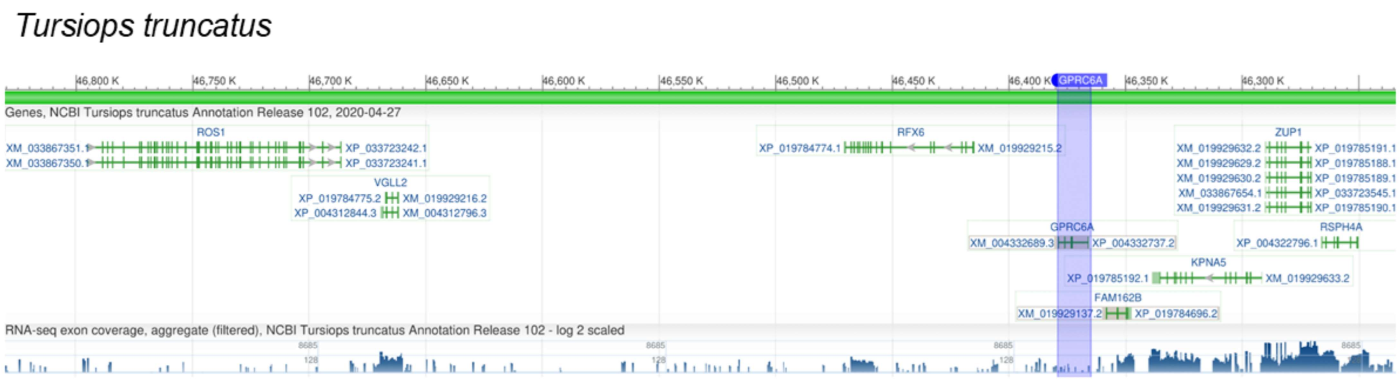


**Supplementary Figure S14. Synteny of the GPRC6A Region in *Tursiops truncatus*:** This figure illustrates the synteny of the genomic region containing the *GPRC6A* gene in *Tursiops truncatus*, as visualized using the NCBI Genome Data Viewer. The highlighted region specifically marks the *GPRC6A* gene and its flanking genes, providing a detailed view of the genomic context in which *GPRC6A* resides. This figure highlights that the genomic region around *GPRC6A* is conserved.

S15

*Bos taurus*

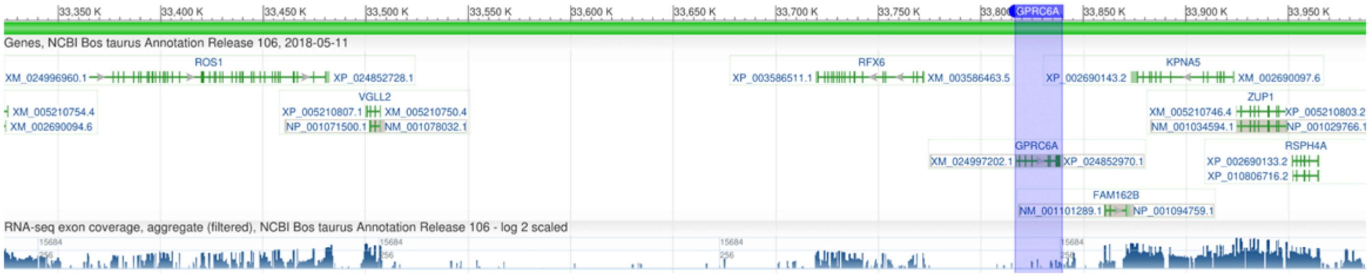


**Supplementary Figure S15. Synteny of the *GPRC6A* Region in *Bos taurus*:** This figure illustrates the synteny of the genomic region containing the *GPRC6A* gene in *Bos taurus*, as visualized using the NCBI Genome Data Viewer. The highlighted region specifically marks the *GPRC6A* gene and its flanking genes, providing a detailed view of the genomic context in which *GPRC6A* resides. This figure highlights that the genomic region around *GPRC6A* is conserved.

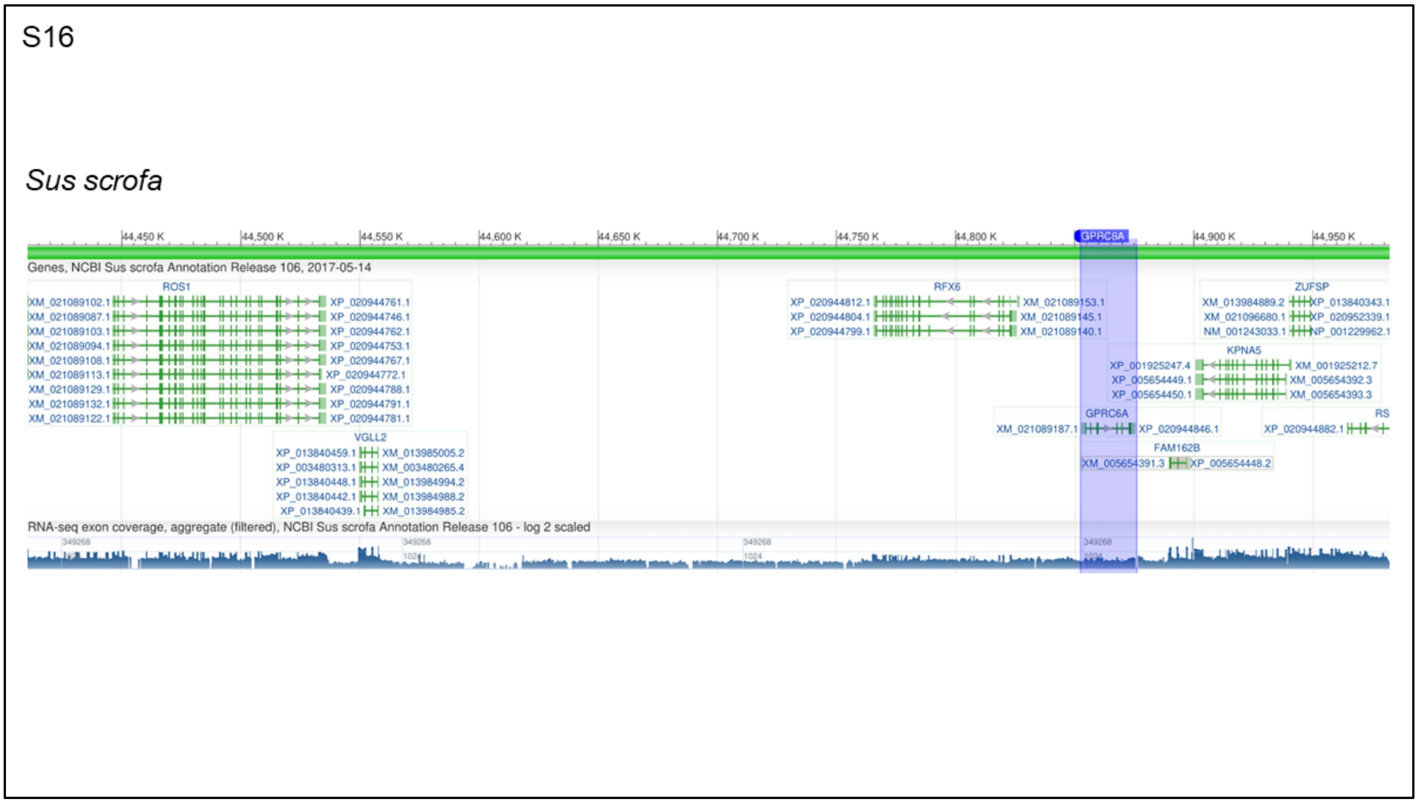


**Supplementary Figure S16. Synteny of the GPRC6A Region in *Sus scrofa*:** This figure illustrates the synteny of the genomic region containing the *GPRC6A* gene in *Sus scrofa*, as visualized using the NCBI Genome Data Viewer. The highlighted region specifically marks the *GPRC6A* gene and its flanking genes, providing a detailed view of the genomic context in which *GPRC6A* resides. This figure highlights that the genomic region around *GPRC6A* is conserved.

S17

*Camelus bactrianus*

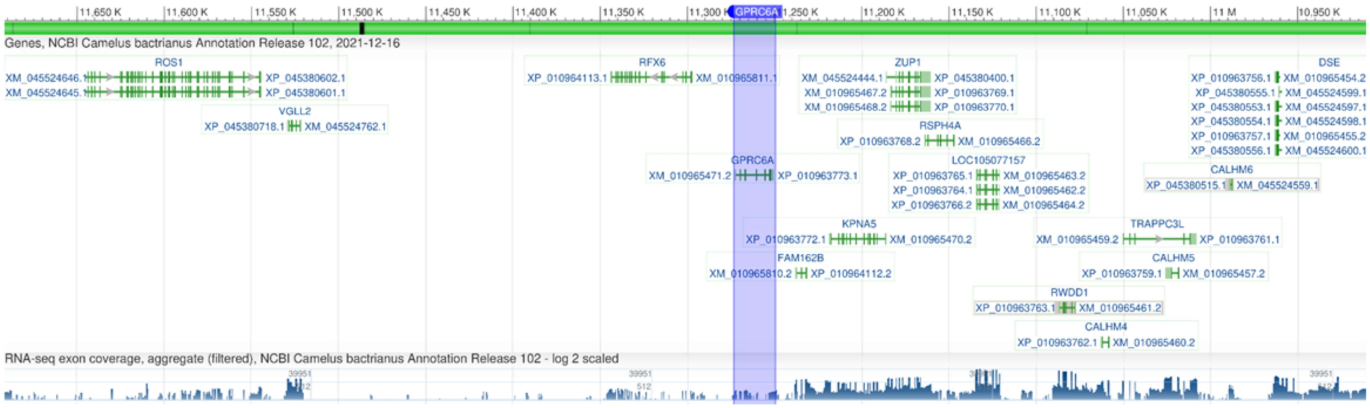


**Supplementary Figure S17. Synteny of the *GPRC6A* Region in *Camelus bactrianus***: This figure illustrates the synteny of the genomic region containing the *GPRC6A* gene in *Camelus bactrianus*, as visualized using the NCBI Genome Data Viewer. The highlighted region specifically marks the *GPRC6A* gene and its flanking genes, providing a detailed view of the genomic context in which *GPRC6A* resides. This figure highlights that the genomic region around *GPRC6A* is conserved.

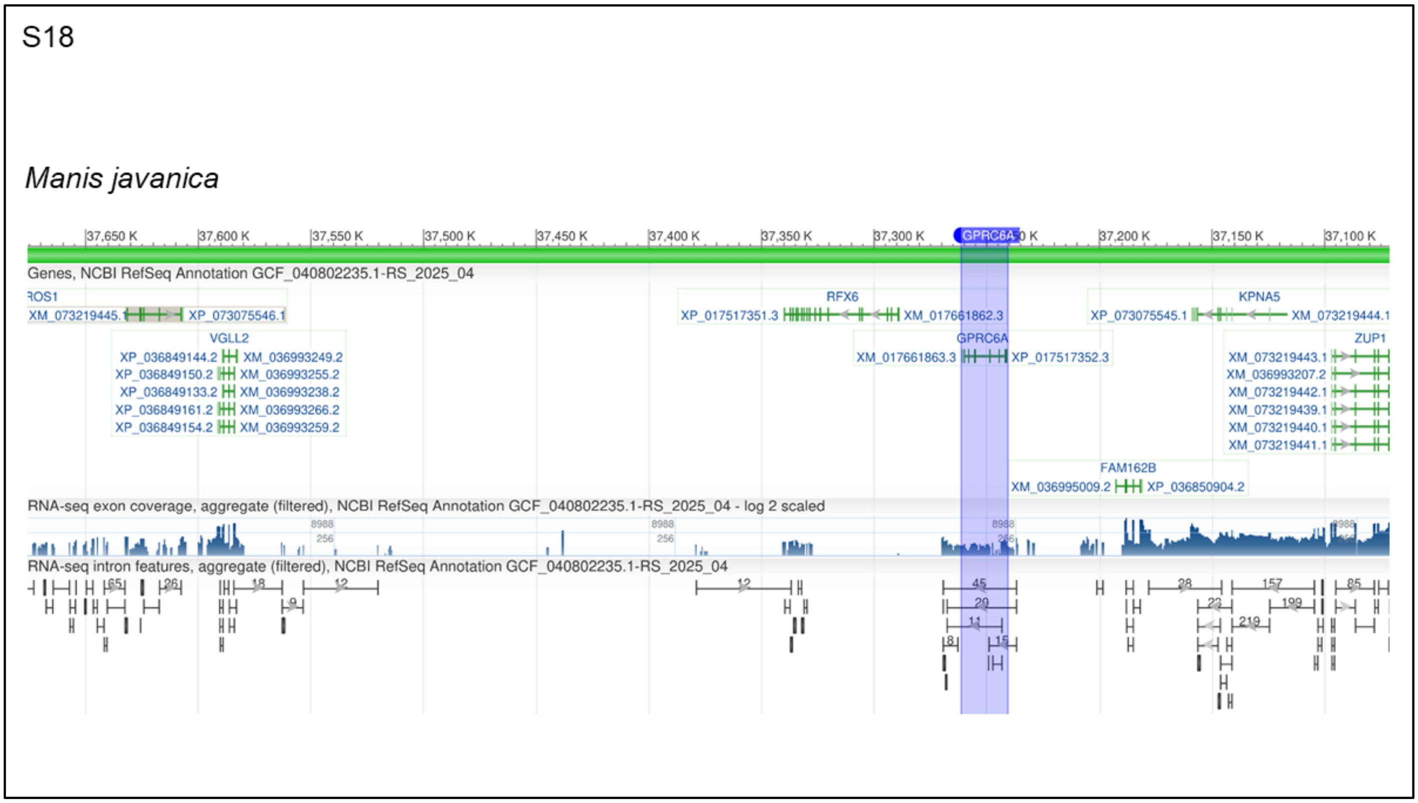


**Supplementary Figure S18. Synteny of the *GPRC6A* Region in *Manis javanica*:** This figure illustrates the synteny of the genomic region containing the *GPRC6A* gene in *Manis javanica*, as visualized using the NCBI Genome Data Viewer. The highlighted region specifically marks the *GPRC6A* gene and its flanking genes, providing a detailed view of the genomic context in which *GPRC6A* resides. This figure highlights that the genomic region around *GPRC6A* is conserved.

S19

*Canis lupus*

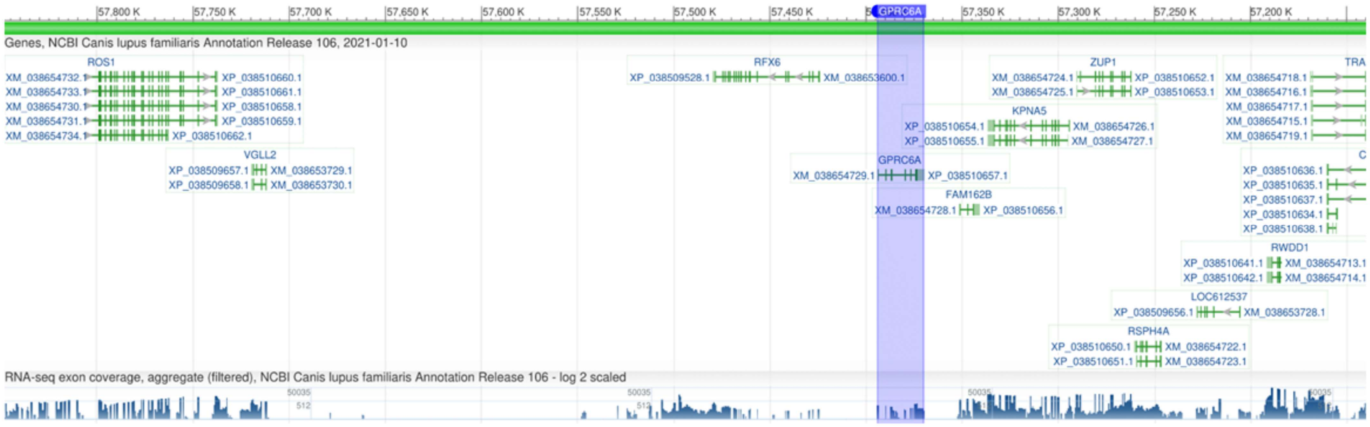


**Supplementary Figure S19. Synteny of the *GPRC6A* Region in *Canis lupus*:** This figure illustrates the synteny of the genomic region containing the *GPRC6A* gene in *Canis lupus*, as visualized using the NCBI Genome Data Viewer. The highlighted region specifically marks the *GPRC6A* gene and its flanking genes, providing a detailed view of the genomic context in which *GPRC6A* resides. This figure highlights that the genomic region around *GPRC6A* is conserved.

S20

*Equus caballus*

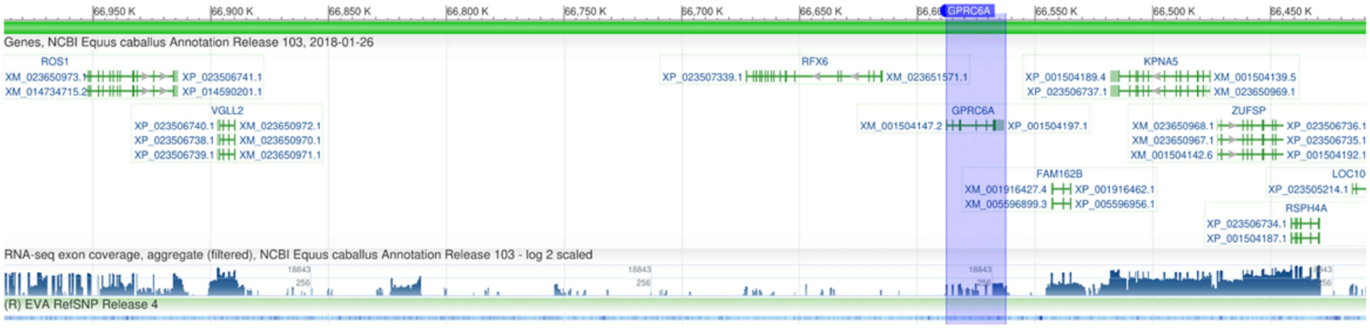


**Supplementary Figure S20. Synteny of the *GPRC6A* Region in *Equus caballus*:** This figure illustrates the synteny of the genomic region containing the *GPRC6A* gene in *Equus caballus*, as visualized using the NCBI Genome Data Viewer. The highlighted region specifically marks the *GPRC6A* gene and its flanking genes, providing a detailed view of the genomic context in which *GPRC6A* resides. This figure highlights that the genomic region around *GPRC6A* is conserved.

S21

*Rhinolophus ferrumequinum*

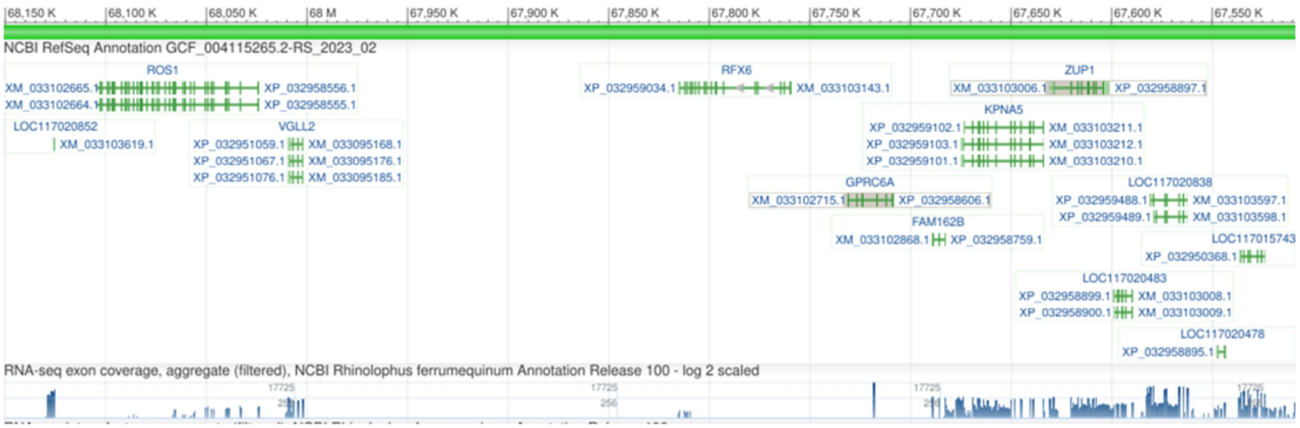


**Supplementary Figure S21. Synteny of *GPRC6A* region in *Rhinolophus ferrumequinum***: This figure illustrates the synteny of the genomic region containing the *GPRC6A* gene in *Rhinolophus ferrumequinum*, as visualized using the NCBI Genome Data Viewer. The highlighted region specifically marks the *GPRC6A* gene and its flanking genes, providing a detailed view of the genomic context in which *GPRC6A* resides. This figure highlights that the genomic region around *GPRC6A* is conserved.

S22

*Myotis myotis*

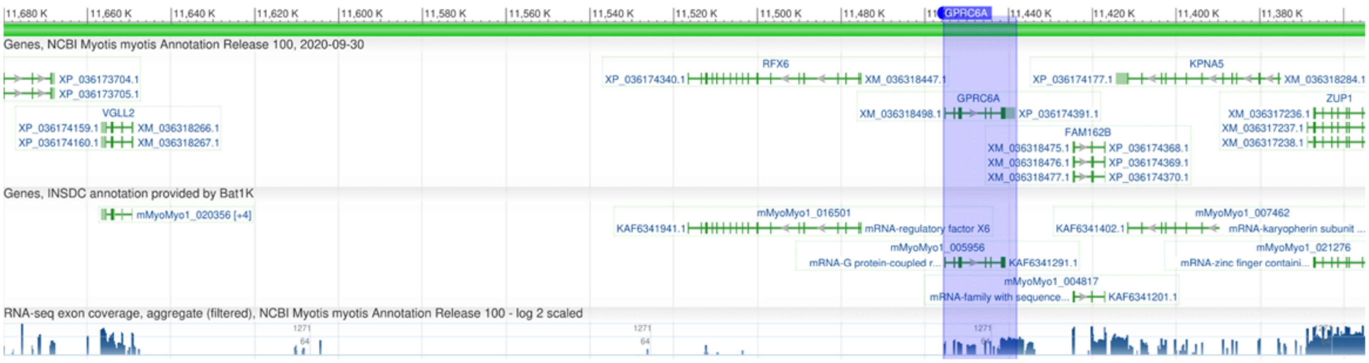


**Supplementary Figure S22. Synteny of the *GPRC6A* Region in *Myotis myotis*:** This figure illustrates the synteny of the genomic region containing the *GPRC6A* gene in *Myotis myotis*, as visualized using the NCBI Genome Data Viewer. The highlighted region specifically marks the *GPRC6A* gene and its flanking genes, providing a detailed view of the genomic context in which *GPRC6A* resides. This figure highlights that the genomic region around *GPRC6A* is conserved.

S23

*Pteropus giganteus*

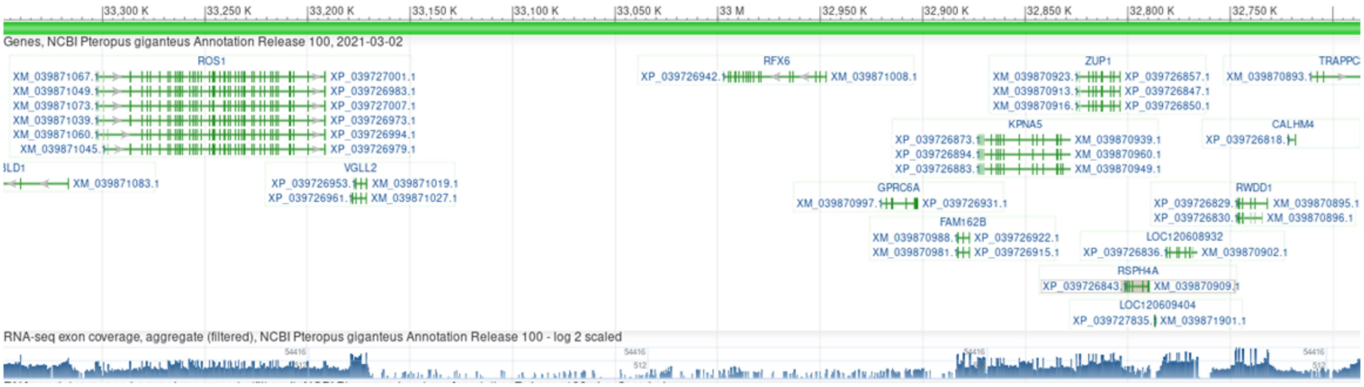


**Supplementary Figure S23. Synteny of *GPRC6A* Region in *Pteropus giganteus*:** This figure illustrates the synteny of the genomic region containing the *GPRC6A* gene in *Pteropus giganteus*, as visualized using the NCBI Genome Data Viewer. The highlighted region specifically marks the *GPRC6A* gene and its flanking genes, providing a detailed view of the genomic context in which *GPRC6A* resides. This figure highlights that the genomic region around *GPRC6A* is conserved.

S24

*Suncus etruscus*

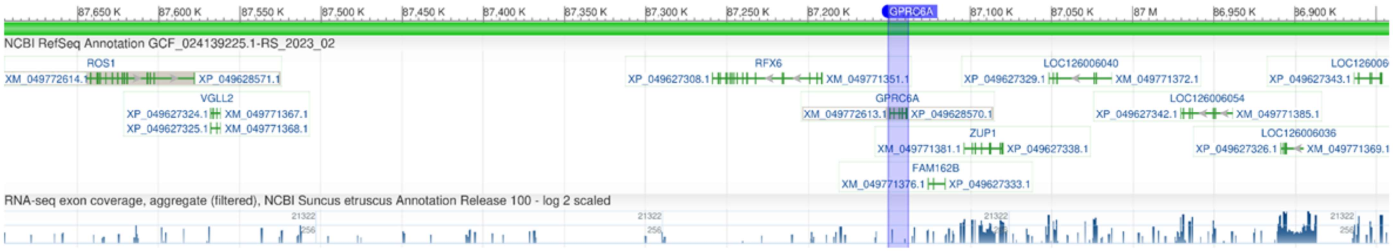


**Supplementary Figure S24. Synteny of the *GPRC6A* Region in *Suncus etruscus*:** This figure illustrates the synteny of the genomic region containing the *GPRC6A* gene in *Suncus etruscus*, as visualized using the NCBI Genome Data Viewer. The highlighted region specifically marks the *GPRC6A* gene and its flanking genes, providing a detailed view of the genomic context in which *GPRC6A* resides. This figure highlights that the genomic region around *GPRC6A* is conserved.

S25

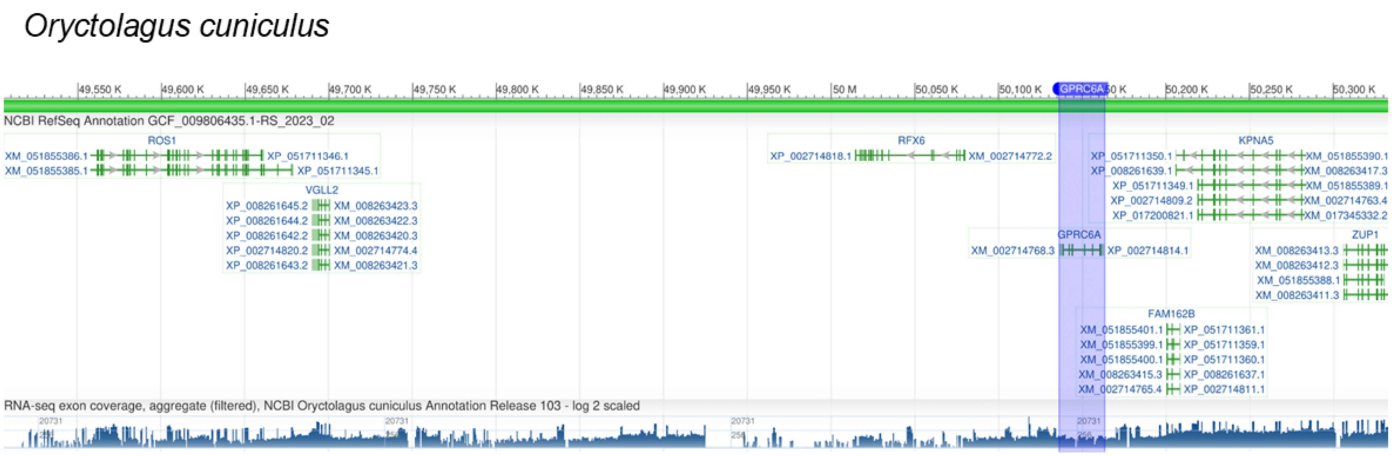


**Supplementary Figure S25. Synteny of the *GPRC6A* Region in *Oryctolagus cuniculus***: This figure illustrates the synteny of the genomic region containing the *GPRC6A* gene in *Oryctolagus cuniculus*, as visualized using the NCBI Genome Data Viewer. The highlighted region specifically marks the *GPRC6A* gene and its flanking genes, providing a detailed view of the genomic context in which *GPRC6A* resides. This figure highlights that the genomic region around *GPRC6A* is conserved.

S26

Ochotona princeps

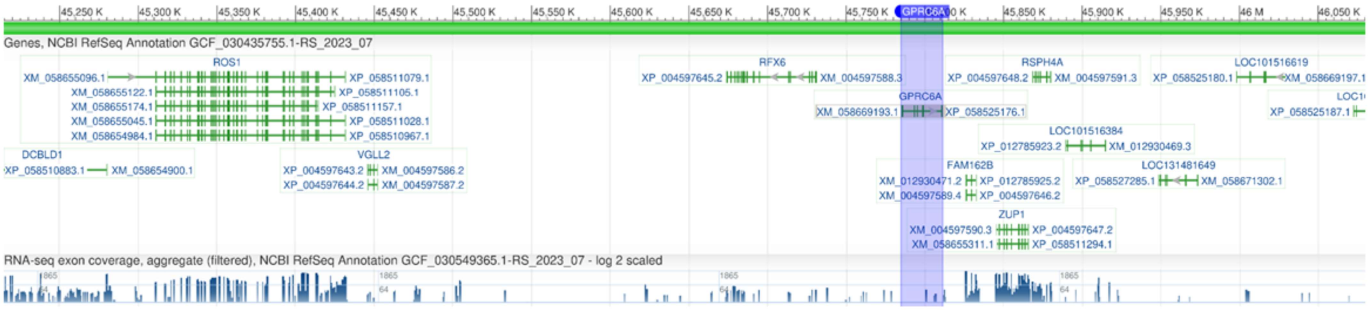

**Supplementary Figure S26. Synteny of the *GPRC6A* Region in *Ochotona princeps***: This figure illustrates the synteny of the genomic region containing the *GPRC6A* gene in *Ochotona princeps*, as visualized using the NCBI Genome Data Viewer. The highlighted region specifically marks the *GPRC6A* gene and its flanking genes, providing a detailed view of the genomic context in which *GPRC6A* resides. This figure highlights that the genomic region around *GPRC6A* is conserved.

S27

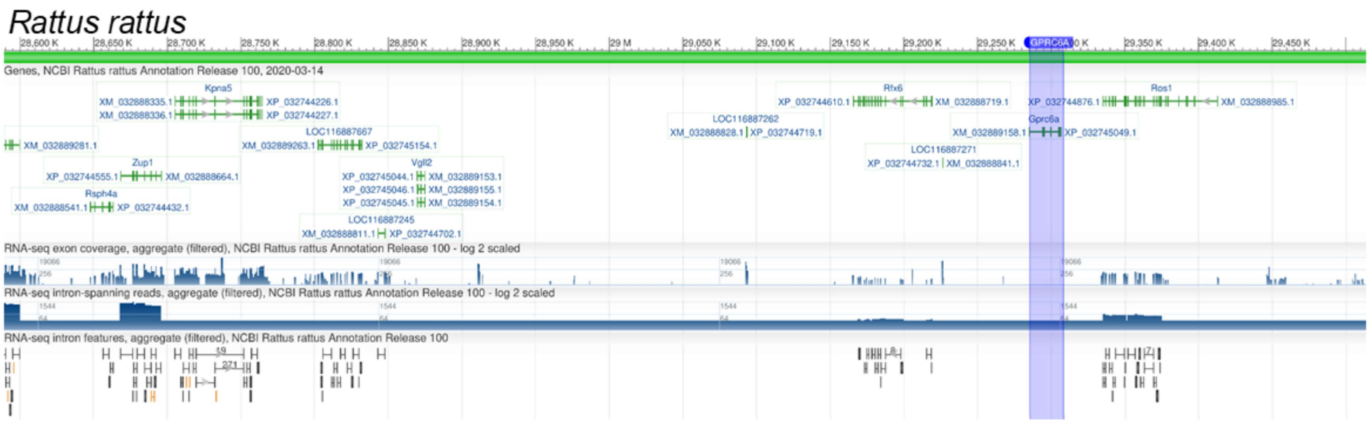


**Supplementary Figure S27. Synteny of the *GPRC6A* Region in *Rattus rattus*:** This figure illustrates the synteny of the genomic region containing the *GPRC6A* gene in *Rattus rattus*, as visualized using the NCBI Genome Data Viewer. The highlighted region specifically marks the *GPRC6A* gene and its flanking genes, providing a detailed view of the genomic context in which *GPRC6A* resides. This figure highlights that the genomic region around *GPRC6A* in *Rattus rattus* is shuffled and might be lying on an evolutionary breakpoint region.

S28

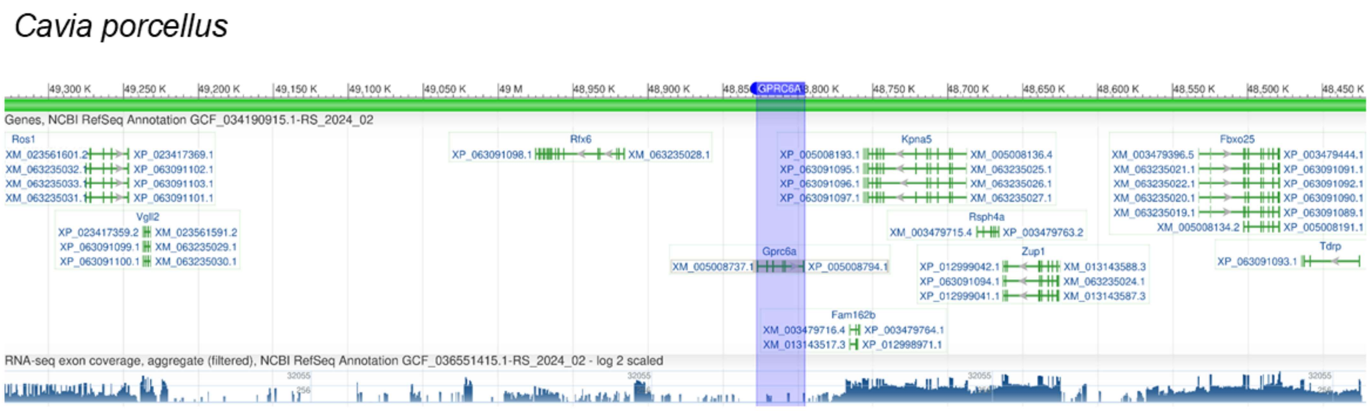


**Supplementary Figure S28. Synteny of the *GPRC6A* Region in *Cavia porcellus*:** This figure illustrates the synteny of the genomic region containing the *GPRC6A* gene in *Cavia porcellus*, as visualized using the NCBI Genome Data Viewer. The highlighted region specifically marks the *GPRC6A* gene and its flanking genes, providing a detailed view of the genomic context in which *GPRC6A* resides. This figure highlights that the genomic region around *GPRC6A* is conserved.

S29

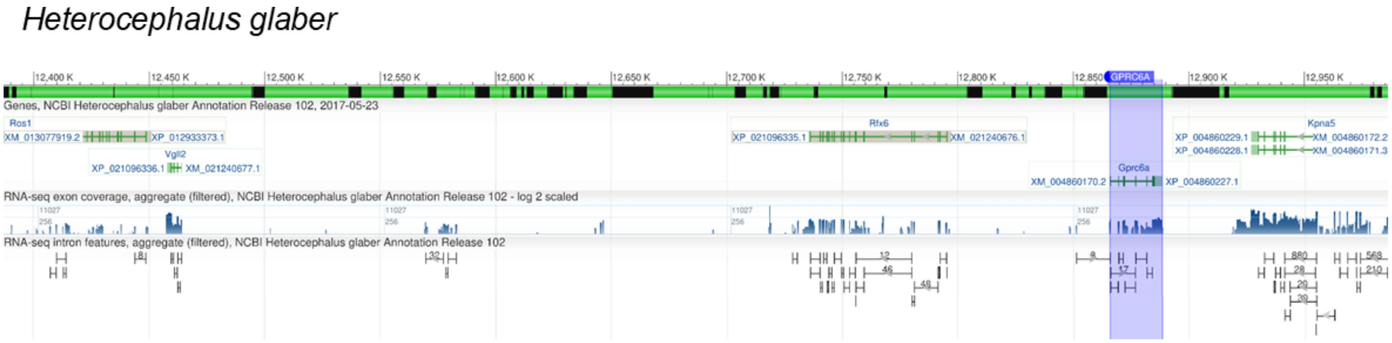


**Supplementary Figure S29. Synteny of the *GPRC6A* region in *Heterocephalus glaber***: This figure illustrates the synteny of the genomic region containing the *GPRC6A* gene in *Heterocephalus glaber*, as visualized using the NCBI Genome Data Viewer. The highlighted region specifically marks the *GPRC6A* gene and its flanking genes, providing a detailed view of the genomic context in which *GPRC6A* resides. This figure highlights that the genomic region around *GPRC6A* is conserved.

S30

*Trichechus manatus*

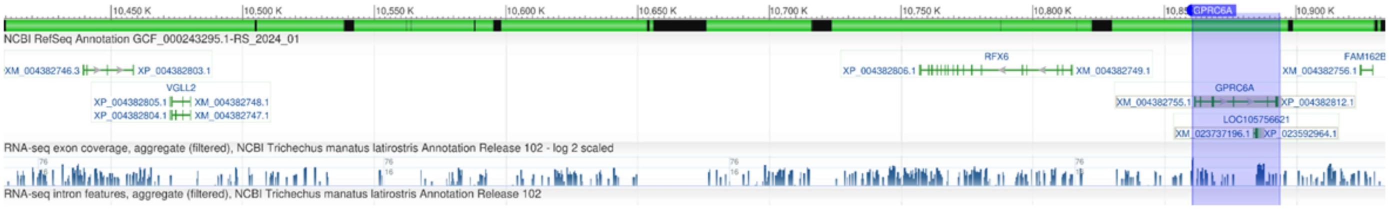


**Supplementary Figure S30. Synteny of the *GPRC6A* region in *Trichechus manatus*:** This figure illustrates the synteny of the genomic region containing the *GPRC6A* gene in *Trichechus manatus*, as visualized using the NCBI Genome Data Viewer. The highlighted region specifically marks the *GPRC6A* gene and its flanking genes, providing a detailed view of the genomic context in which *GPRC6A* resides. This figure highlights that the genomic region around *GPRC6A* is conserved.

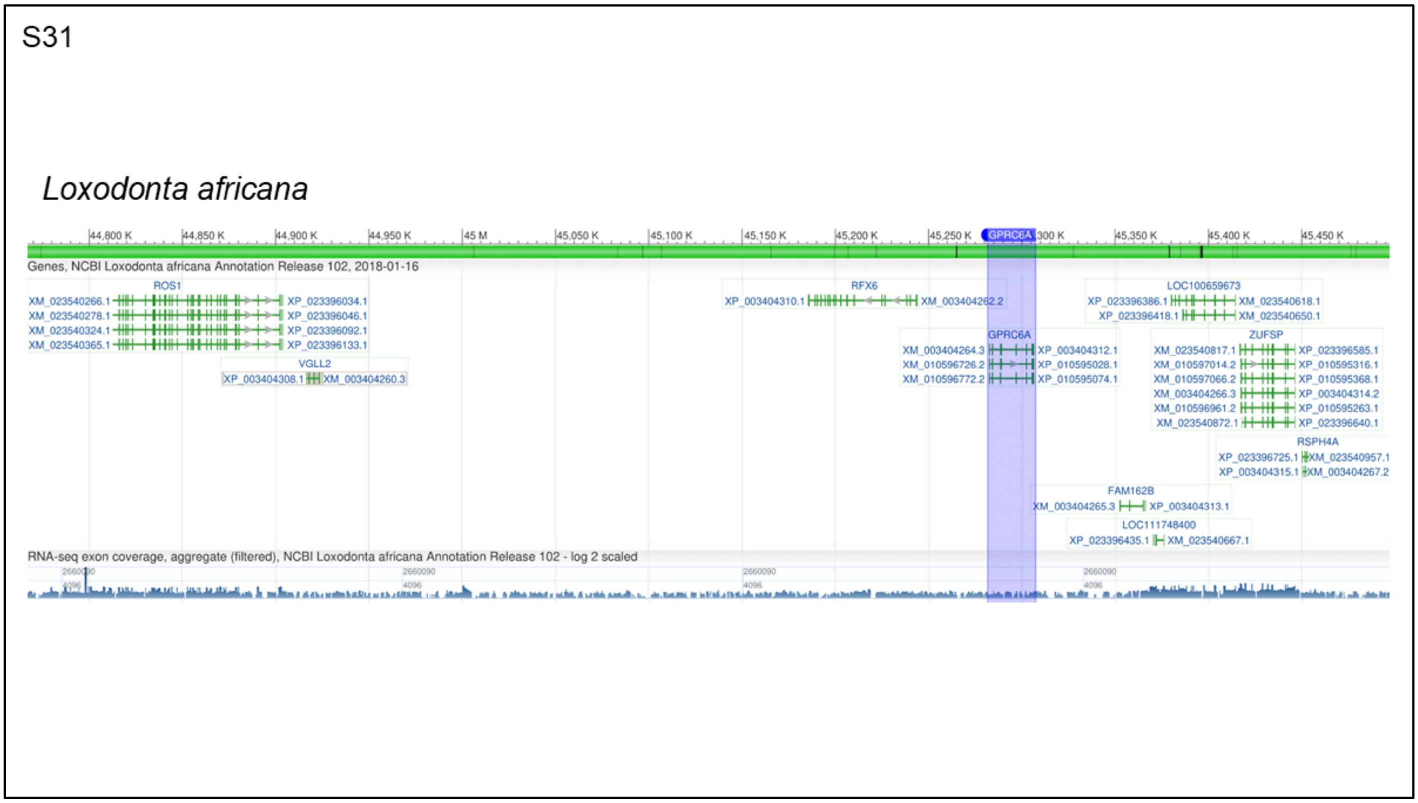


**Supplementary Figure S31. Synteny of the *GPRC6A* region in *Loxodonta africana***: This figure illustrates the synteny of the genomic region containing the *GPRC6A* gene in *Loxodonta africana*, as visualized using the NCBI Genome Data Viewer. The highlighted region specifically marks the *GPRC6A* gene and its flanking genes, providing a detailed view of the genomic context in which *GPRC6A* resides. This figure highlights that the genomic region around *GPRC6A* is conserved.

S32

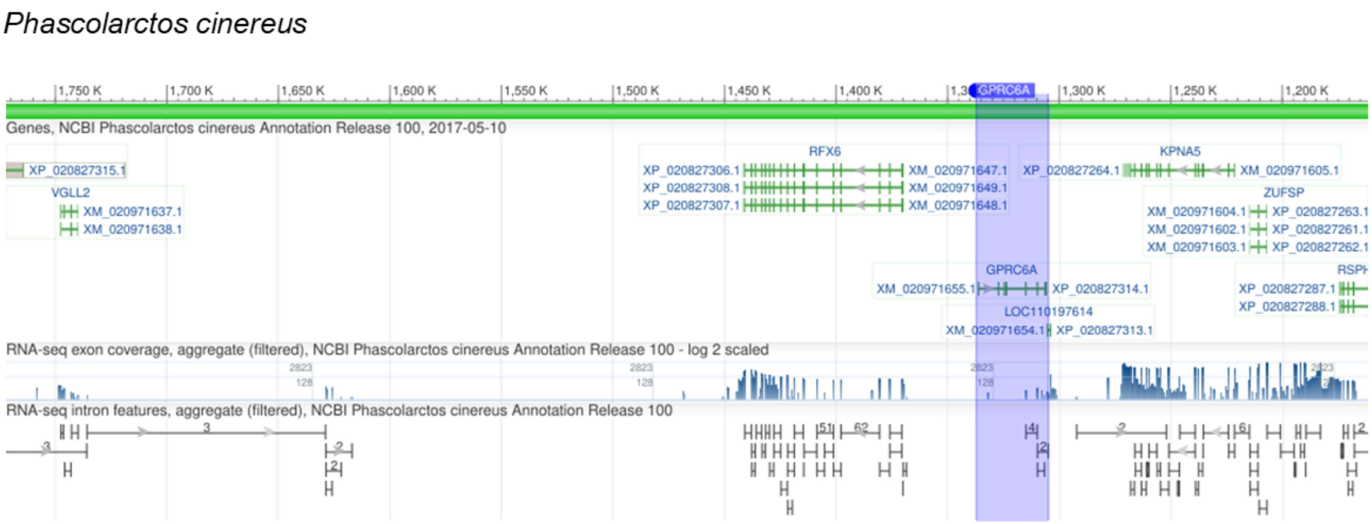

**Supplementary Figure S32. Synteny of the *GPRC6A* region in *Phascolarctos cinereus*:** This figure illustrates the synteny of the genomic region containing the *GPRC6A* gene in *Phascolarctos cinereus*, as visualized using the NCBI Genome Data Viewer. The highlighted region specifically marks the *GPRC6A* gene and its flanking genes, providing a detailed view of the genomic context in which *GPRC6A* resides. This figure highlights that the genomic region around *GPRC6A* is conserved.

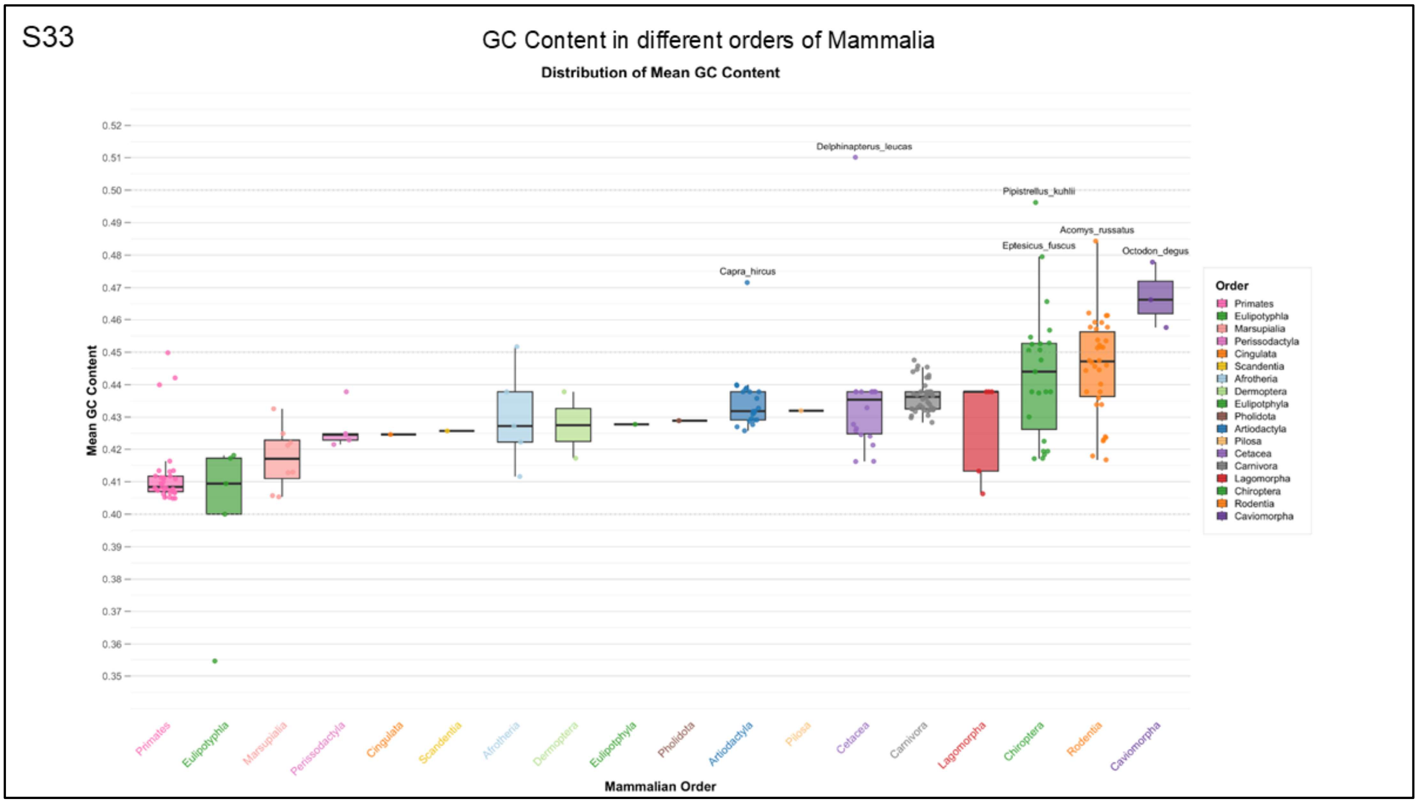


**Supplementary Figure S33. Distribution of mean GC content across mammalian orders**. Boxplots show the range and variation of mean GC content within each order, with individual species with high GC labeled for reference (e.g., *Acomys russatus*, *Capra hircus*, *Delphinapterus leucas*, *Eptesicus fuscus*, *Octodon degus*, and *Pipistrellus kuhlii*). The figure highlights inter-order differences in GC composition, with rodents and bats (Chiroptera) exhibiting distinct patterns compared to other mammalian groups.

S34

RNA-seq status of *GPRC6A* in *Homo sapiens*

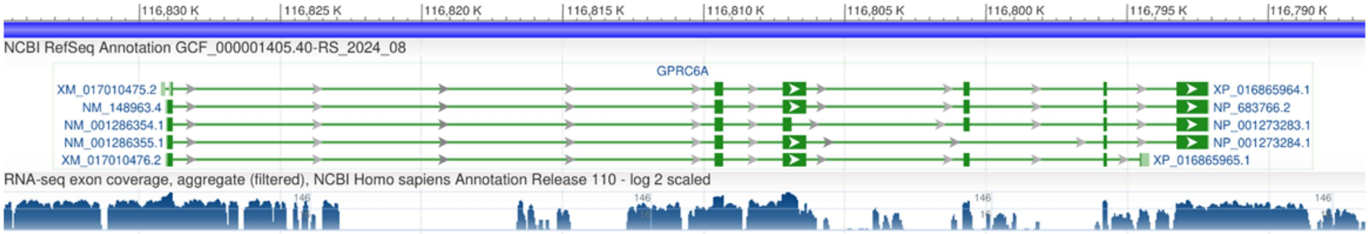


**Supplementary Figure S34. RNA-seq expression in *Homo sapiens:*** This figure displays the RNA-seq expression profile of *GPRC6A* in *Homo sapiens*, visualized using the NCBI Genome Data Viewer. Expression levels are shown across various tissues, with the X-axis representing distinct tissue types and the Y-axis indicating relative expression.

S35

RNA-seq status of *GPRC6A* in *Camelus bactrianus*

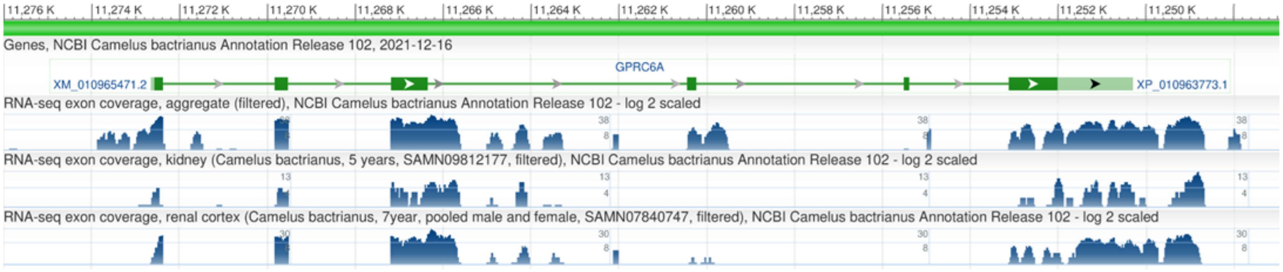


**Supplementary Figure S35.RNA-seq expression in *Camelus bactrianus:*** This figure displays the RNA-seq expression profile of *GPRC6A* in *Camelus bactrianus*, visualized using the NCBI Genome Data Viewer. Expression levels are shown across various tissues, with the X-axis representing distinct tissue types and the Y-axis indicating relative expression.

S36

RNA-seq status of *GPRC6A* in *Myotis myotis*

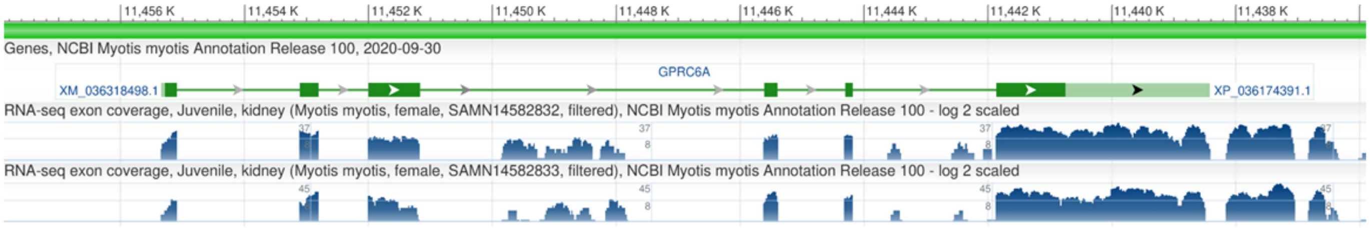


**Supplementary Figure S36. RNA-seq expression in *Myotis myotis:*** This figure displays the RNA-seq expression profile of *GPRC6A* in *Myotis myotis*, visualized using the NCBI Genome Data Viewer. Expression levels are shown across various tissues, with the X-axis representing distinct tissue types and the Y-axis indicating relative expression.

S37

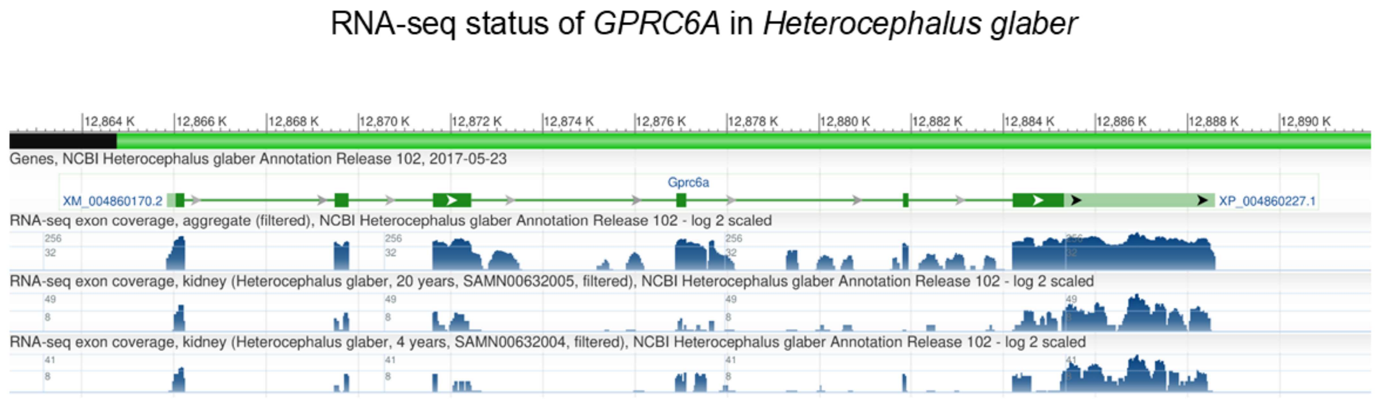


**Supplementary Figure S37. RNA-seq expression in *Heterocephalus glaber:*** This figure displays the RNA-seq expression profile of *GPRC6A* in *Heterocephalus glaber*, visualized using the NCBI Genome Data Viewer. Expression levels are shown across various tissues, with the X-axis representing distinct tissue types and the Y-axis indicating relative expression.

S38

RNA-seq status of *GPRC6A* in *Mus musculus*

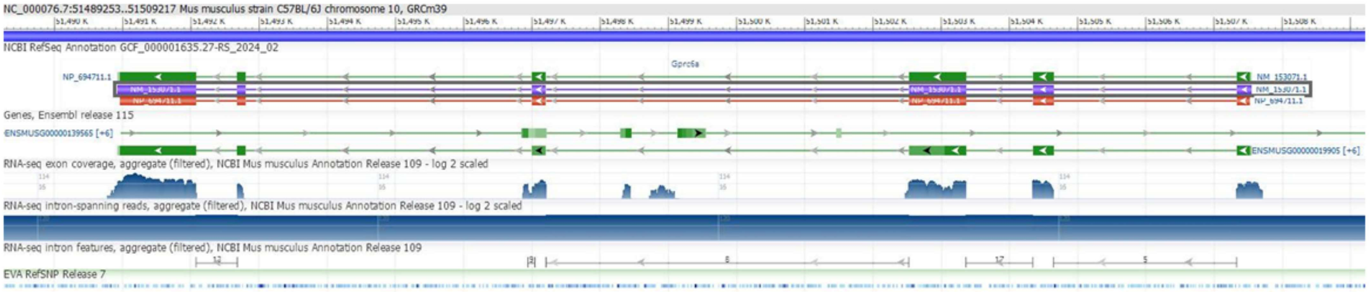


**Supplementary Figure S38. RNA-seq expression in *Mus musculus:*** This figure displays the RNA-seq expression profile of *GPRC6A* in *Mus musculus*, visualized using the NCBI Genome Data Viewer. Filtered aggregates of RNA-seq exon coverage, intron-spanning reads and intron features are shown as separate tracks.

S39

### RNA-seq status of *GPRC6A* in *Rattus norvegicus*

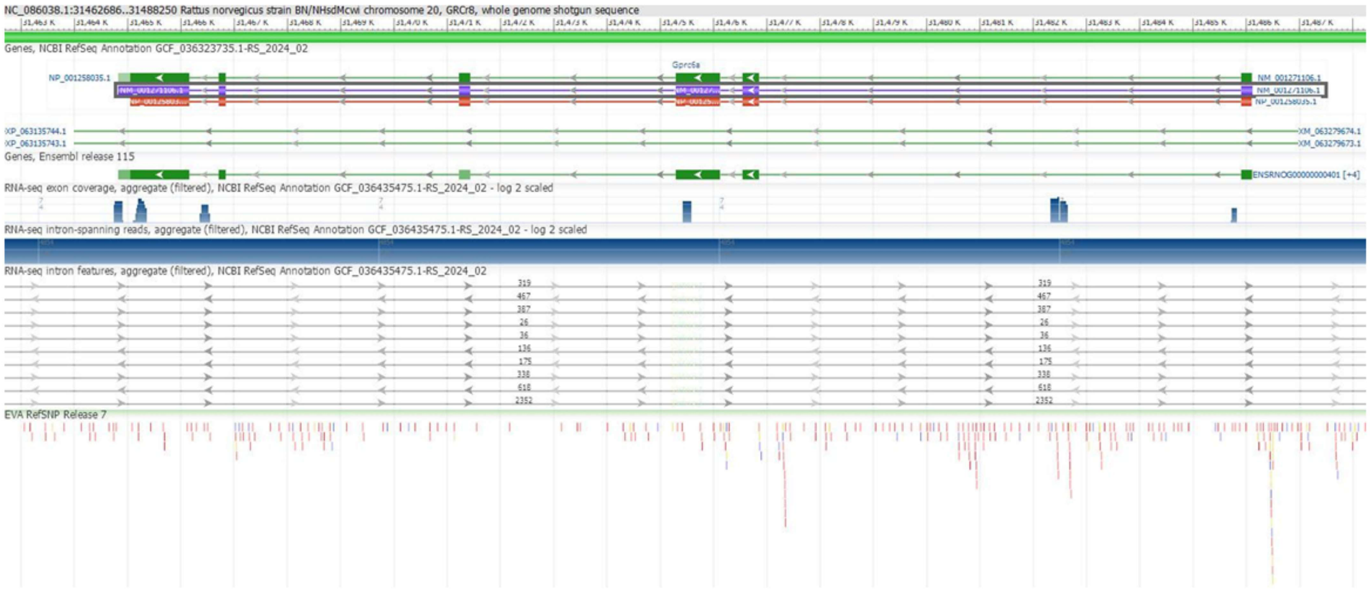


**Supplementary Figure S39. RNA-seq expression in *Rattus norvegicus:*** This figure displays the RNA-seq expression profile of *GPRC6A* in *Rattus norvegicus*, visualized using the NCBI Genome Data Viewer. Filtered aggregates of RNA-seq exon coverage, intron-spanning reads and intron features are shown as separate tracks.

S40

RNA-seq status of *GPRC6A* in *Ailuropoda melanoleuca*

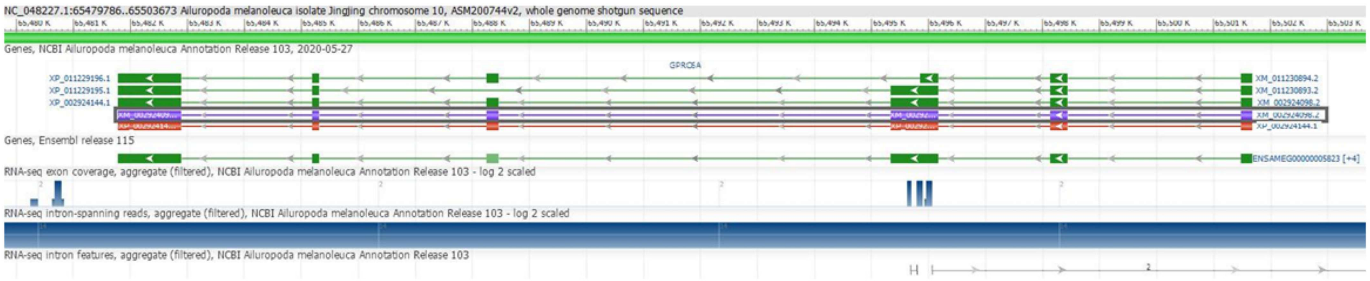


**Supplementary Figure S40. RNA-seq expression in *Ailuropoda melanoleuca:*** This figure displays the RNA-seq expression profile of *GPRC6A* in *Ailuropoda melanoleuca*, visualized using the NCBI Genome Data Viewer. Filtered aggregates of RNA-seq exon coverage, intron-spanning reads and intron features are shown as separate tracks.

S41

RNA-seq status of *GPRC6A* in *Bos taurus*

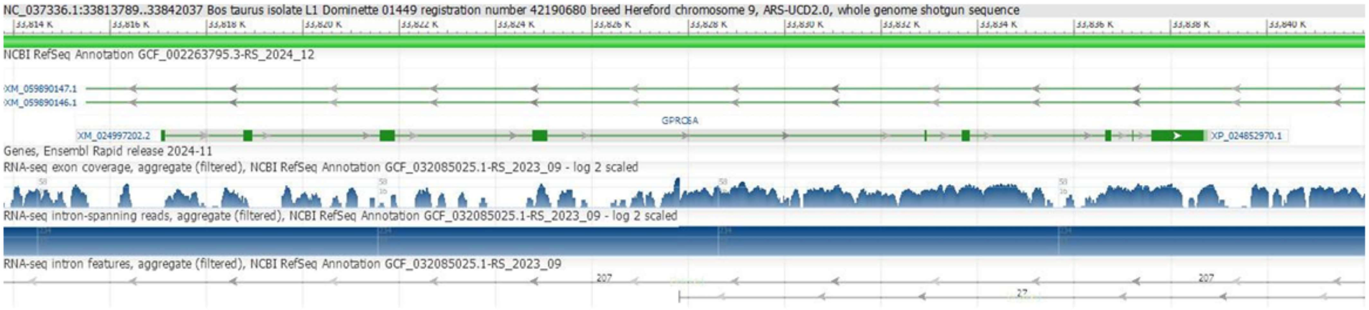


**Supplementary Figure S41. RNA-seq expression in *Bos taurus:*** This figure displays the RNA-seq expression profile of *GPRC6A* in *Bos taurus*, visualized using the NCBI Genome Data Viewer. Filtered aggregates of RNA-seq exon coverage, intron-spanning reads and intron features are shown as separate tracks.

S42

RNA-seq status of *GPRC6A* in *Bubalus bubalis*

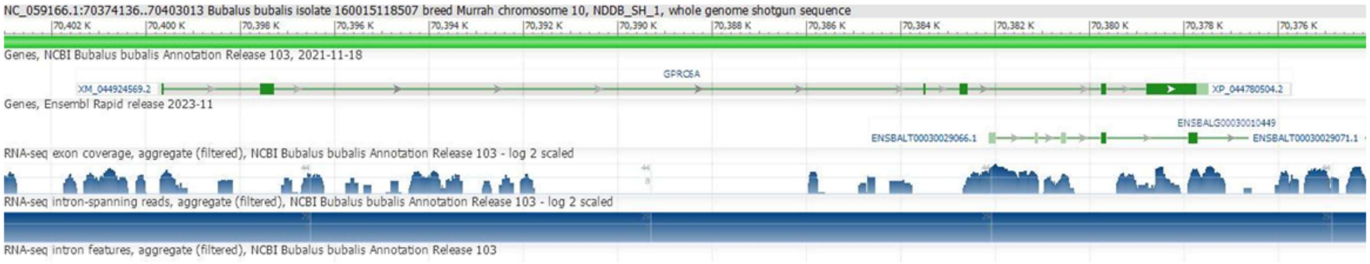


**Supplementary Figure S42. RNA-seq expression in *Bubalus bubalis:*** This figure displays the RNA-seq expression profile of *GPRC6A* in *Bubalus bubalis*, visualized using the NCBI Genome Data Viewer. Filtered aggregates of RNA-seq exon coverage, intron-spanning reads and intron features are shown as separate tracks.

S43

RNA-seq status of *GPRC6A* in *Capra hircus*

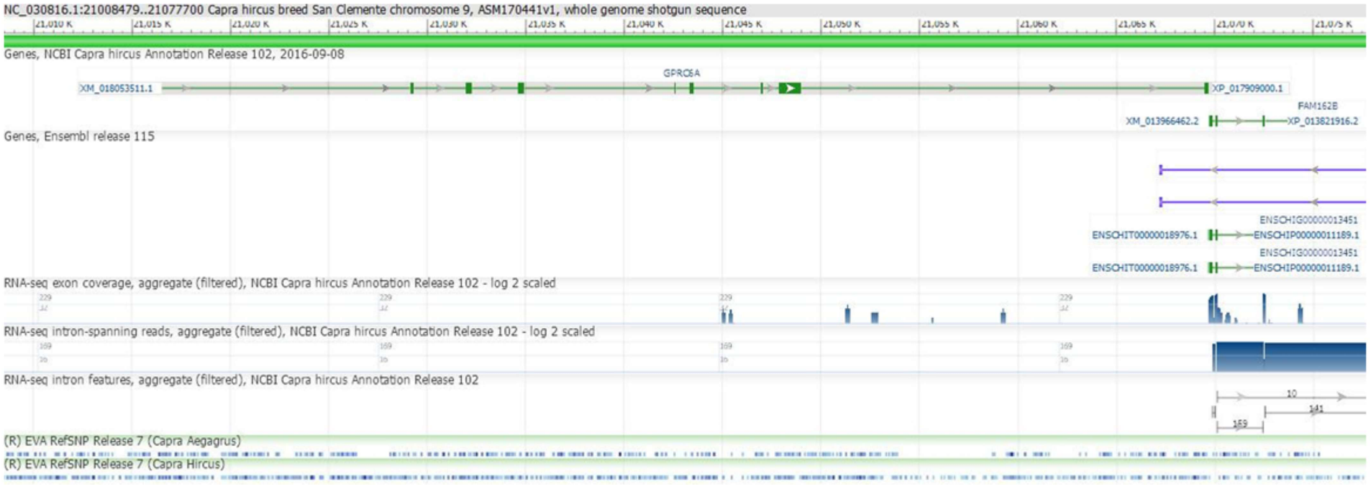


**Supplementary Figure S43. RNA-seq expression in *Capra hircus:*** This figure displays the RNA-seq expression profile of *GPRC6A* in *Capra hircus*, visualized using the NCBI Genome Data Viewer. Filtered aggregates of RNA-seq exon coverage, intron-spanning reads and intron features are shown as separate tracks.

S44

RNA-seq status of *GPRC6A* in *Castor canadensis*

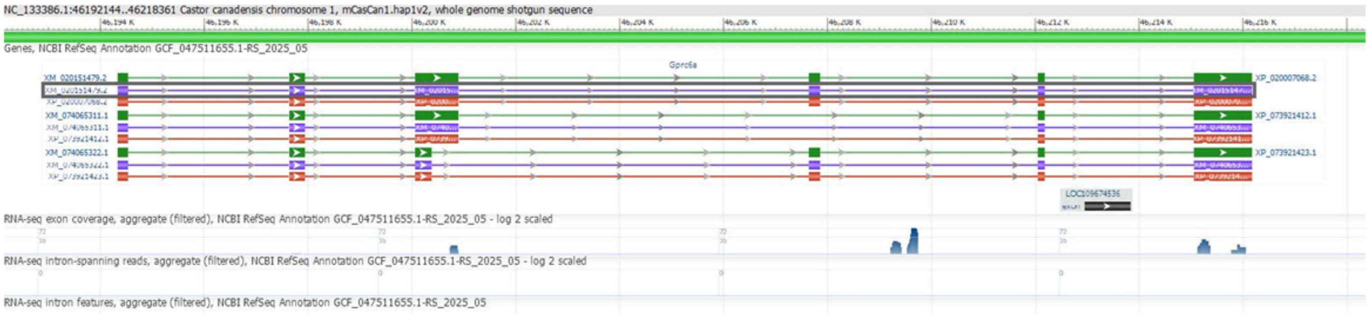


**Supplementary Figure S44. RNA-seq expression in *Castor canadensis:*** This figure displays the RNA-seq expression profile of *GPRC6A* in *Castor canadensis*, visualized using the NCBI Genome Data Viewer. Filtered aggregates of RNA-seq exon coverage, intron-spanning reads and intron features are shown as separate tracks.

S45

RNA-seq status of *GPRC6A* in *Monodelphis domestica*

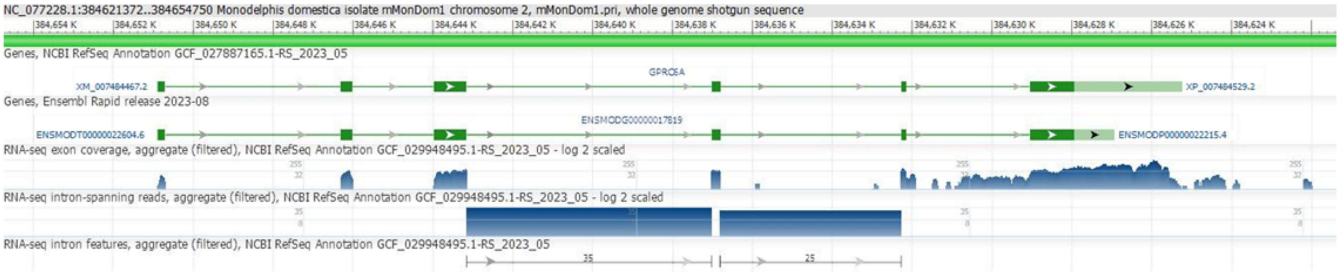


**Supplementary Figure S45. RNA-seq expression in *Monodelphis domestica:*** This figure displays the RNA-seq expression profile of *GPRC6A* in *Monodelphis domestica* visualized using the NCBI Genome Data Viewer. Filtered aggregates of RNA-seq exon coverage, intron-spanning reads and intron features are shown as separate tracks.

S46

RNA-seq status of *GPRC6A* in *Dasypus novemcinctus*

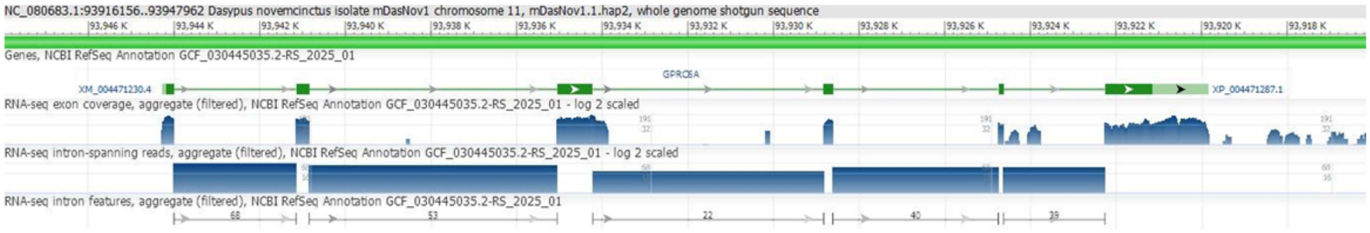


**Supplementary Figure S46. RNA-seq expression in *Dasypus novemcinctus:***
This figure displays the RNA-seq expression profile of *GPRC6A* in *Dasypus novemcinctus* visualized using the NCBI Genome Data Viewer. Filtered aggregates of RNA-seq exon coverage, intron-spanning reads and intron features are shown as separate tracks.

S47

### RNA-seq status of *GPRC6A* in *Echinops telfairi*

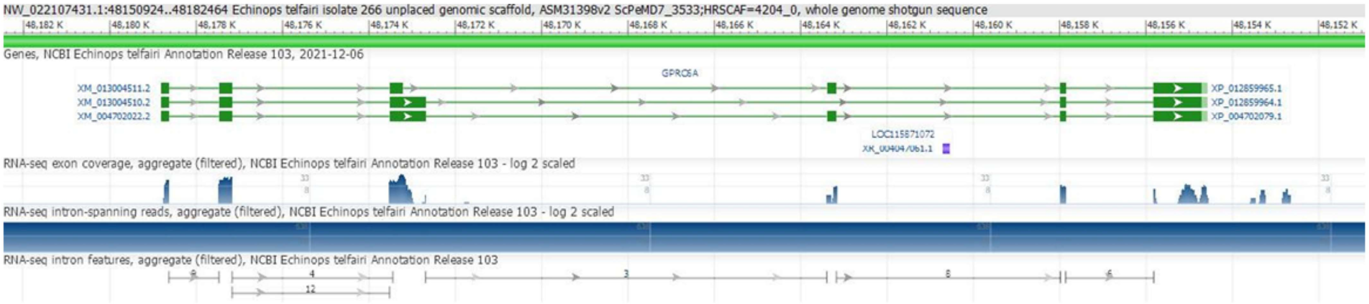


**Supplementary Figure S47. RNA-seq expression in *Echinops telfairi:*** This figure displays the RNA-seq expression profile of *GPRC6A* in *Echinops telfairi*, visualized using the NCBI Genome Data Viewer. Filtered aggregates of RNA-seq exon coverage, intron-spanning reads and intron features are shown as separate tracks.

S48

### RNA-seq status of *GPRC6A* in *Felis catus*

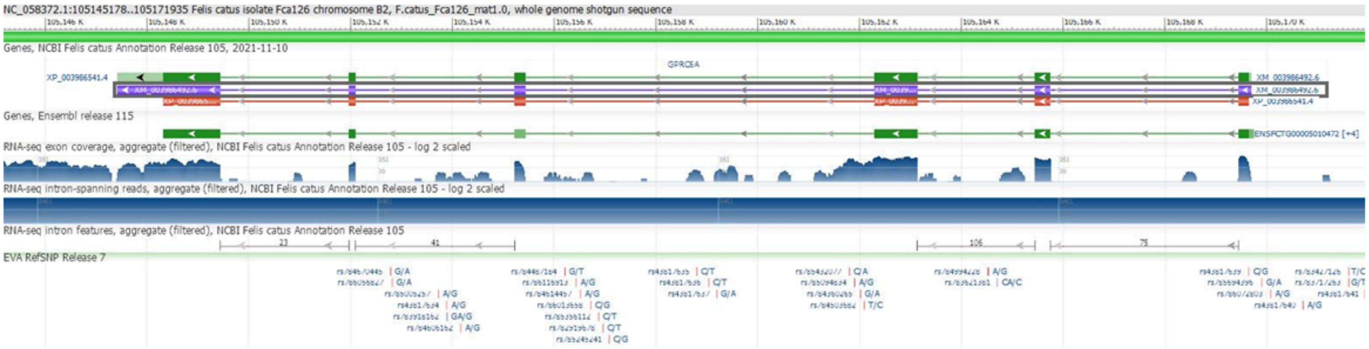


**Supplementary Figure S48. RNA-seq expression in *Felis catus:*** This figure displays the RNA-seq expression profile of *GPRC6A* in *Felis catus*, visualized using the NCBI Genome Data Viewer. Filtered aggregates of RNA-seq exon coverage, intron-spanning reads and intron features are shown as separate tracks.

S49

RNA-seq status of *GPRC6A* in *Ictidomys tridecemlineatus*

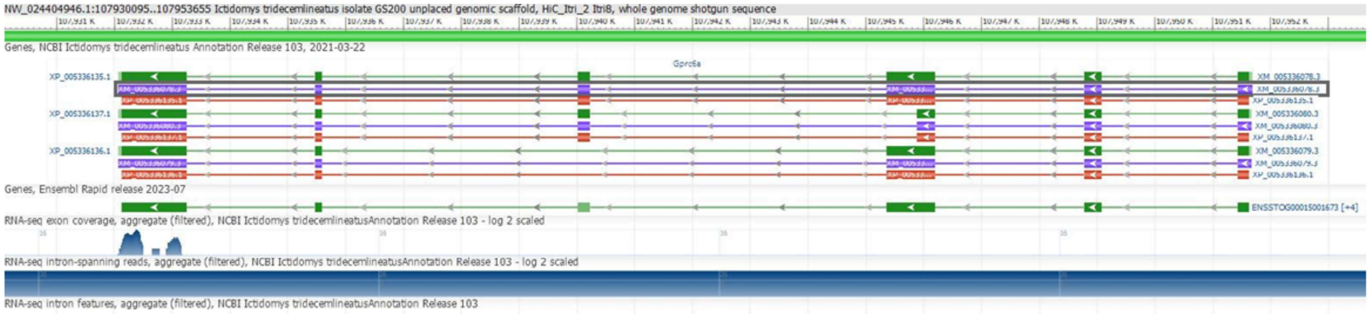


**Supplementary Figure S49. RNA-seq expression in *Ictidomys tridecemlineatus:*** This figure displays the RNA-seq expression profile of *GPRC6A* in *Ictidomys tridecemlineatus*, visualized using the NCBI Genome Data Viewer. Filtered aggregates of RNA-seq exon coverage, intron-spanning reads and intron features are shown as separate tracks.

S50

RNA-seq status of *GPRC6A* in *Loxodonta africana*

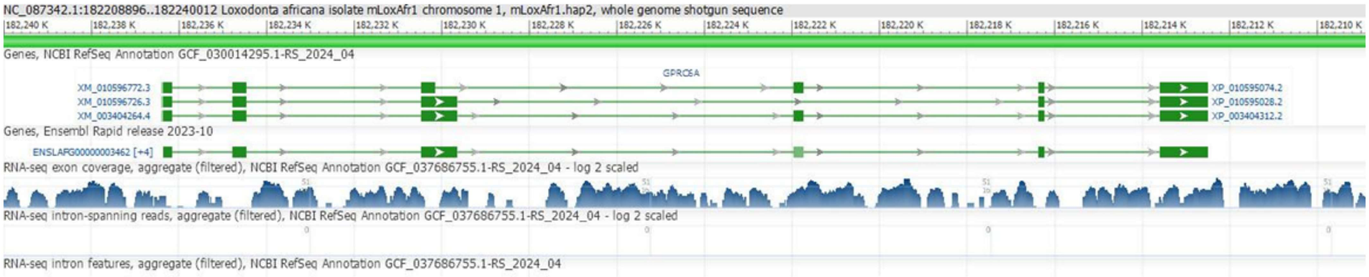


**Supplementary Figure S50. RNA-seq expression in *Loxodonta africana:*** This figure displays the RNA-seq expression profile of *GPRC6A* in *Loxodonta africana*, visualized using the NCBI Genome Data Viewer. Filtered aggregates of RNA-seq exon coverage, intron-spanning reads and intron features are shown as separate tracks.

S51

RNA-seq status of *GPRC6A* in *Nannospalax galili*

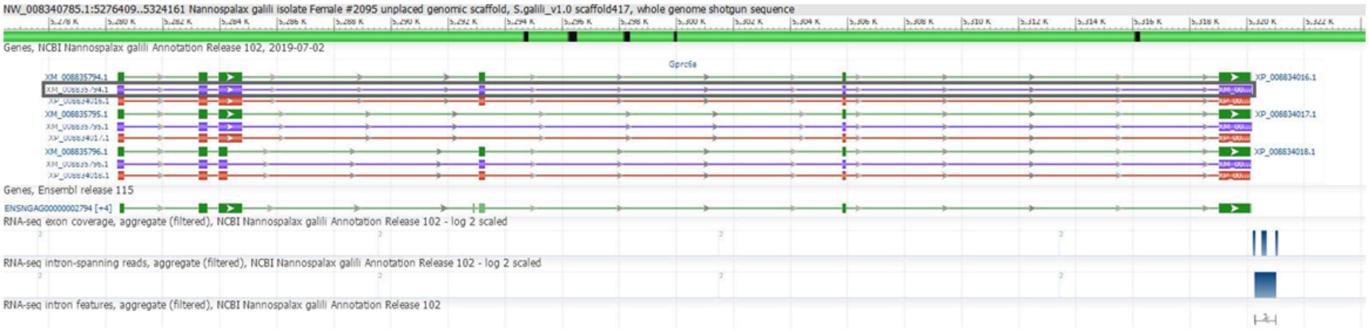


**Supplementary Figure S51. RNA-seq expression in *Nannospalax galili:*** This figure displays the RNA-seq expression profile of *GPRC6A* in *Nannospalax galili*, visualized using the NCBI Genome Data Viewer. Filtered aggregates of RNA-seq exon coverage, intron-spanning reads and intron features are shown as separate tracks.

S52

RNA-seq status of *GPRC6A* in *Ochotona princeps*

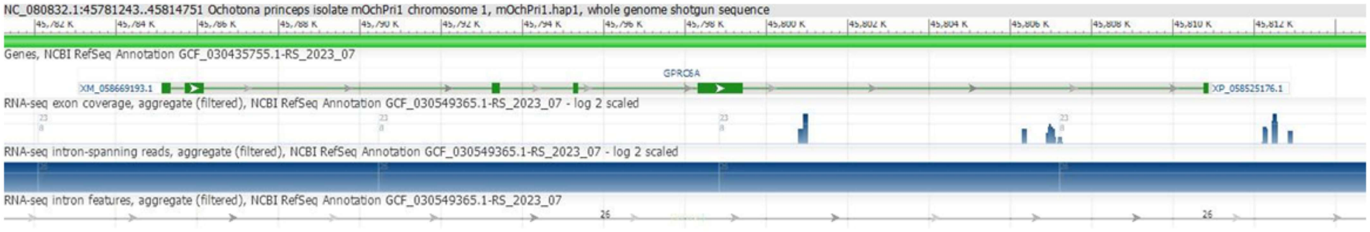


**Supplementary Figure S52. RNA-seq expression in *Ochotona princeps:*** This figure displays the RNA-seq expression profile of *GPRC6A* in *Ochotona princeps*, visualized using the NCBI Genome Data Viewer. Filtered aggregates of RNA-seq exon coverage, intron-spanning reads and intron features are shown as separate tracks.

S53

RNA-seq status of *GPRC6A* in *Orcinus orca*

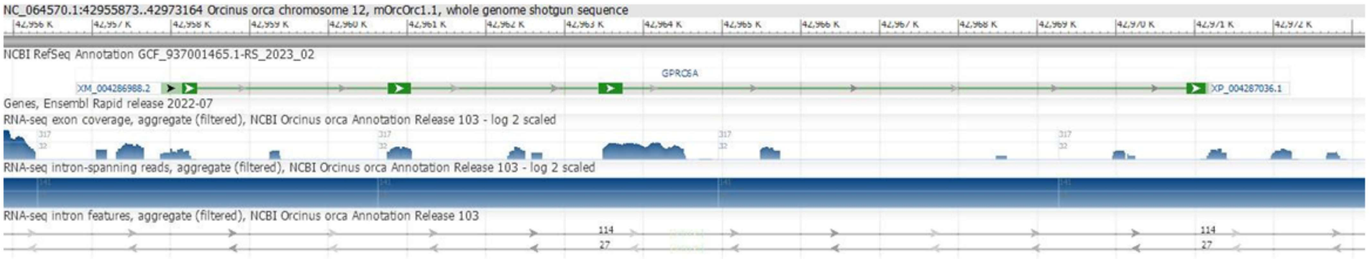


**Supplementary Figure S53. RNA-seq expression in *Orcinus orca:*** This figure displays the RNA-seq expression profile of *GPRC6A* in *Orcinus orca*, visualized using the NCBI Genome Data Viewer. Filtered aggregates of RNA-seq exon coverage, intron-spanning reads and intron features are shown as separate tracks.

S54

RNA-seq status of *GPRC6A* in *Oryctolagus cuniculus*

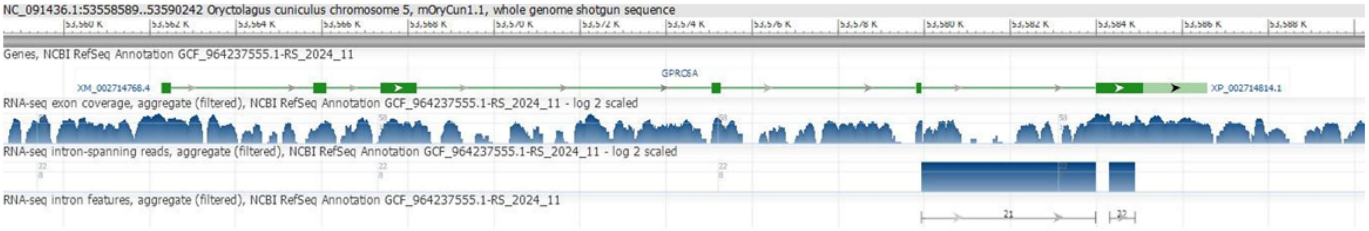


**Supplementary Figure S54. RNA-seq expression in *Oryctolagus cuniculus:*** This figure displays the RNA-seq expression profile of *GPRC6A* in *Oryctolagus cuniculus*, visualized using the NCBI Genome Data Viewer. Filtered aggregates of RNA-seq exon coverage, intron-spanning reads and intron features are shown as separate tracks.

S55

RNA-seq status of *GPRC6A* in *Ovis aries*

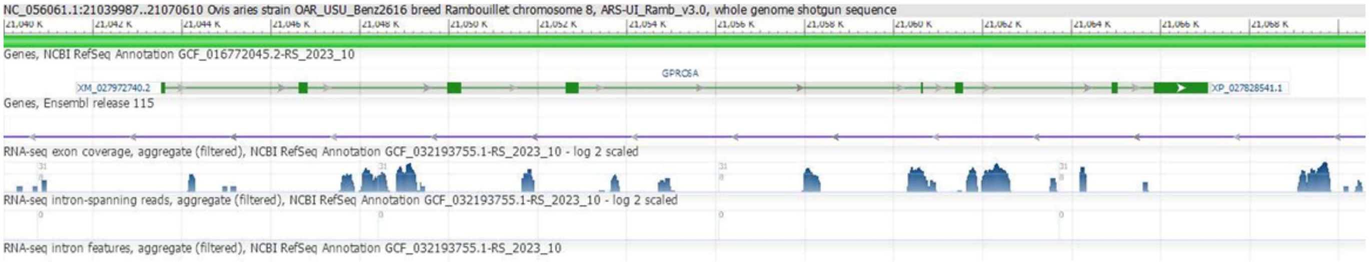


**Supplementary Figure S55. RNA-seq expression in *Ovis aries:*** This figure displays the RNA-seq expression profile of *GPRC6A* in *Ovis aries*, visualized using the NCBI Genome Data Viewer. Filtered aggregates of RNA-seq exon coverage, intron-spanning reads and intron features are shown as separate tracks.

S56

RNA-seq status of *GPRC6A* in *Panthera tigris*

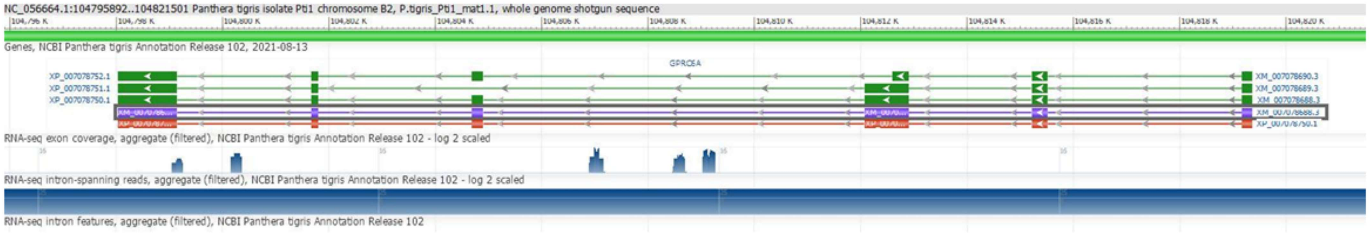


**Supplementary Figure S56. RNA-seq expression in *Panthera tigris:*** This figure displays the RNA-seq expression profile of *GPRC6A* in *Panthera tigris*, visualized using the NCBI Genome Data Viewer. Filtered aggregates of RNA-seq exon coverage, intron-spanning reads and intron features are shown as separate tracks.

S57

RNA-seq status of *GPRC6A* in *Physeter macrocephalus*

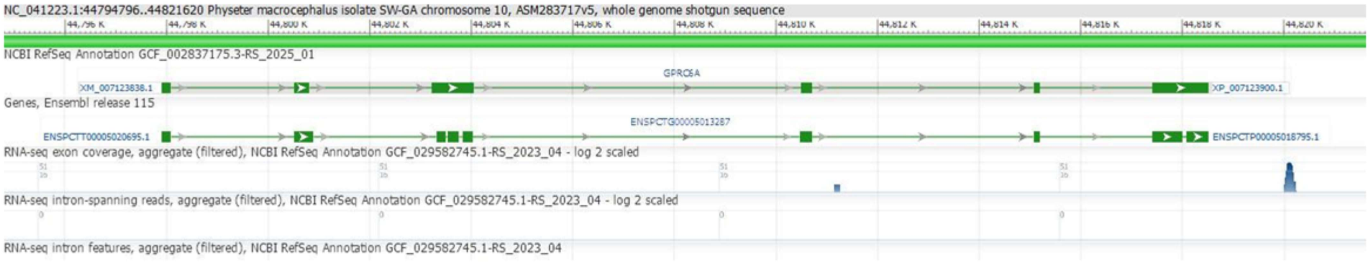


**Supplementary Figure S57. RNA-seq expression in *Physeter macrocephalus:*** This figure displays the RNA-seq expression profile of *GPRC6A* in *Physeter macrocephalus*, visualized using the NCBI Genome Data Viewer. Filtered aggregates of RNA-seq exon coverage, intron-spanning reads and intron features are shown as separate tracks.

S58

RNA-seq status of *GPRC6A* in *Talpa occidentalis*

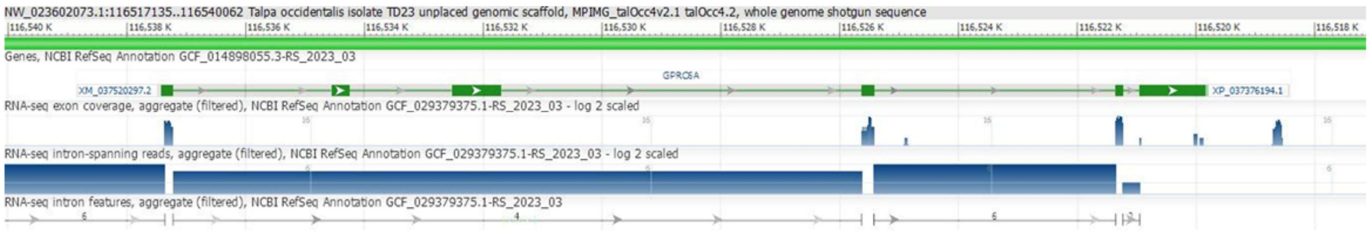


**Supplementary Figure S58. RNA-seq expression in *Talpa occidentalis:*** This figure displays the RNA-seq expression profile of *GPRC6A* in *Talpa occidentalis*, visualized using the NCBI Genome Data Viewer. Filtered aggregates of RNA-seq exon coverage, intron-spanning reads and intron features are shown as separate tracks.

S59

RNA-seq status of *GPRC6A* in *Trichechus manatus latirostris*

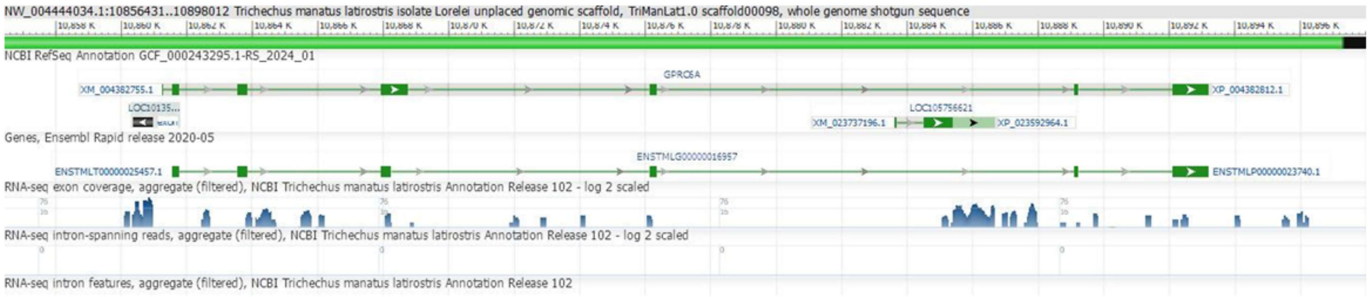


**Supplementary Figure S59. RNA-seq expression in *Trichechus manatus latirostris:*** This figure displays the RNA-seq expression profile of *GPRC6A* in *Trichechus manatus latirostris*, visualized using the NCBI Genome Data Viewer. Filtered aggregates of RNA-seq exon coverage, intron-spanning reads and intron features are shown as separate tracks.

S60

RNA-seq status of *GPRC6A* in *Ursus americanus*

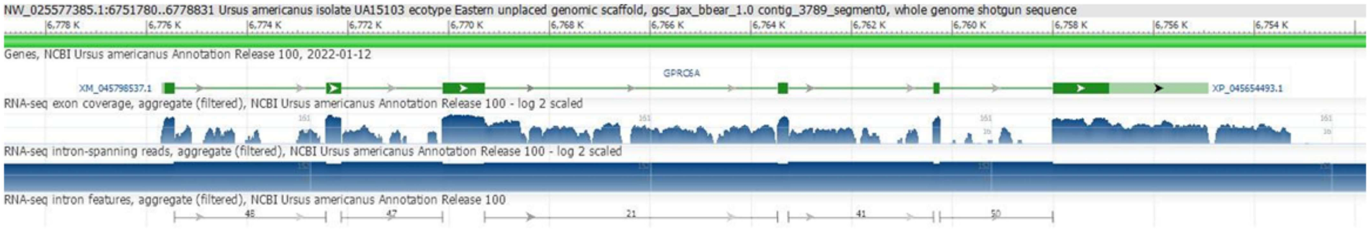


**Supplementary Figure S60. RNA-seq expression in *Ursus americanus:*** This figure displays the RNA-seq expression profile of *GPRC6A* in *Ursus americanus*, visualized using the NCBI Genome Data Viewer. Filtered aggregates of RNA-seq exon coverage, intron-spanning reads and intron features are shown as separate tracks.

S61

RNA-seq status of *GPRC6A* in *Ursus arctos*

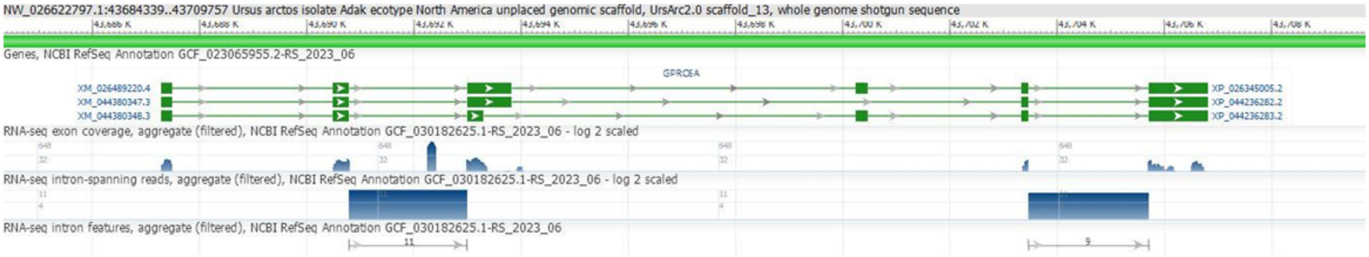


**Supplementary Figure S61. RNA-seq expression in *Ursus arctos:*** This figure displays the RNA-seq expression profile of *GPRC6A* in *Ursus arctos*, visualized using the NCBI Genome Data Viewer. Filtered aggregates of RNA-seq exon coverage, intron-spanning reads and intron features are shown as separate tracks.

S62

RNA-seq status of *GPRC6A* in *Ursus maritimus*

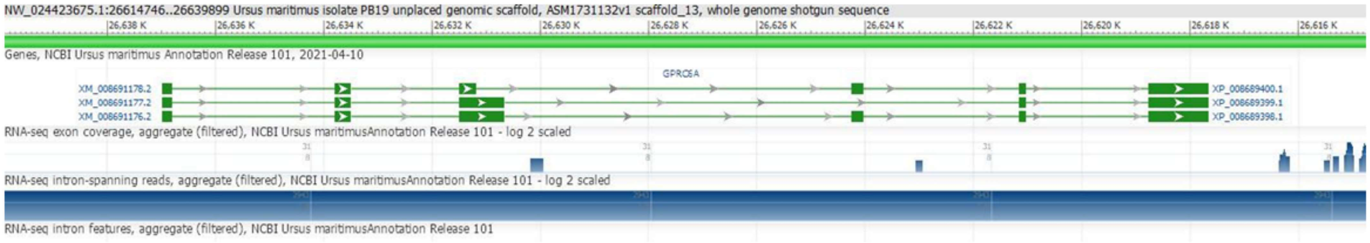


**Supplementary Figure S62. RNA-seq expression in *Ursus maritimus:*** This figure displays the RNA-seq expression profile of *GPRC6A* in *Ursus maritimus*, visualized using the NCBI Genome Data Viewer. Filtered aggregates of RNA-seq exon coverage, intron-spanning reads and intron features are shown as separate tracks.

S63

RNA-seq status of *GPRC6A* in *Chlorocebus sabaeus*

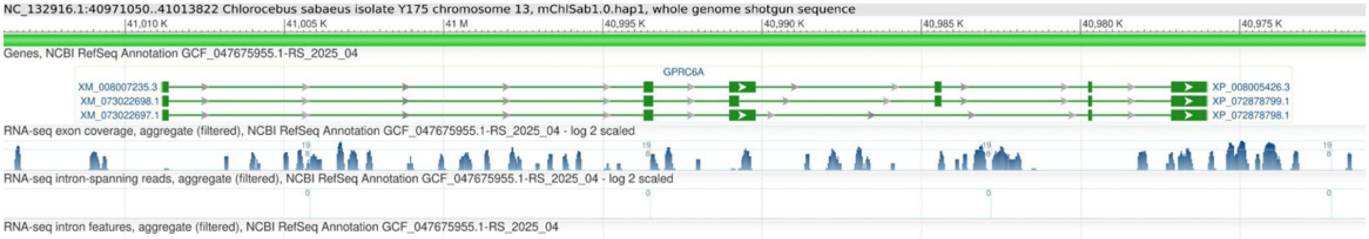


**Supplementary Figure S63. RNA-seq expression in *Chlorocebus sabaeus:*** This figure displays the RNA-seq expression profile of *GPRC6A* in *Chlorocebus sabaeus*, visualized using the NCBI Genome Data Viewer. Filtered aggregates of RNA-seq exon coverage, intron-spanning reads and intron features are shown as separate tracks.

S64

RNA-seq status of *GPRC6A* in *Diceros bicornis minor*

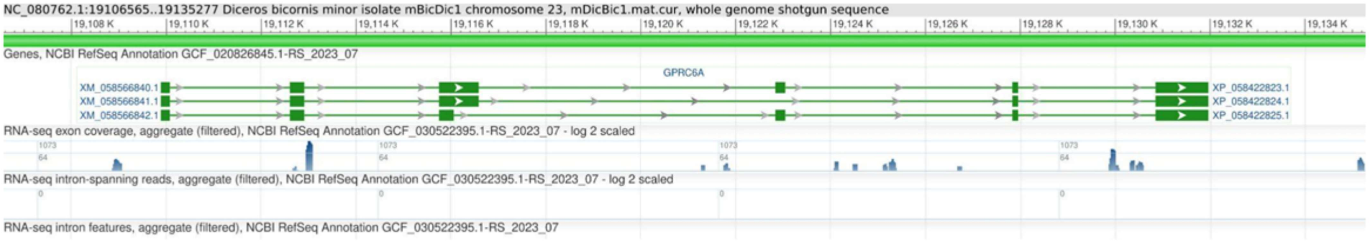


**Supplementary Figure S64. RNA-seq expression in *Diceros bicornis minor:*** This figure displays the RNA-seq expression profile of *GPRC6A* in *Diceros bicornis minor*, visualized using the NCBI Genome Data Viewer. Filtered aggregates of RNA-seq exon coverage, intron-spanning reads and intron features are shown as separate tracks.

S65

RNA-seq status of *GPRC6A* in *Equus asinus*

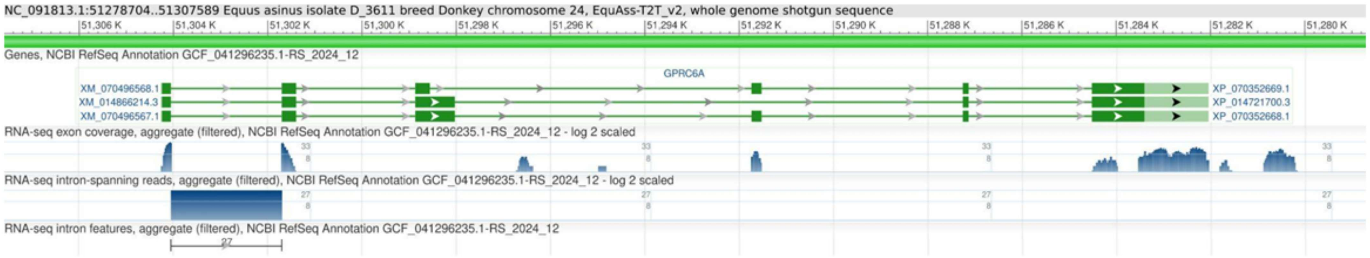


**Supplementary Figure S65. RNA-seq expression in *Equus asinus:*** This figure displays the RNA-seq expression profile of *GPRC6A* in *Equus asinus*, visualized using the NCBI Genome Data Viewer. Filtered aggregates of RNA-seq exon coverage, intron-spanning reads and intron features are shown as separate tracks.

S66

RNA-seq status of *GPRC6A* in *Equus caballus*

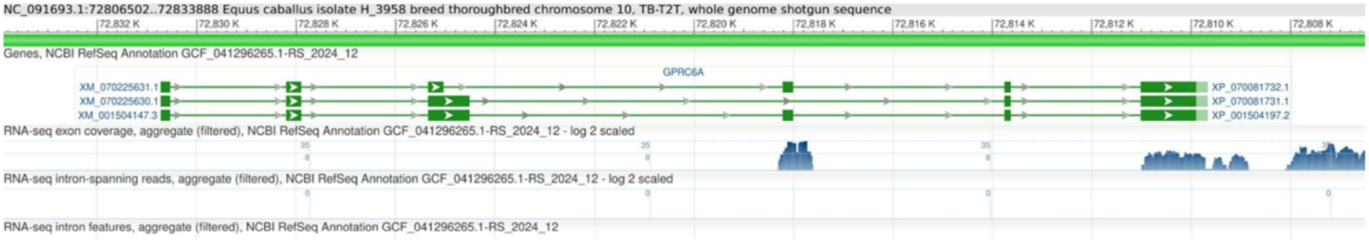


**Supplementary Figure S66. RNA-seq expression in *Equus caballus:*** This figure displays the RNA-seq expression profile of *GPRC6A* in *Equus caballus canadensis*, visualized using the NCBI Genome Data Viewer. Filtered aggregates of RNA-seq exon coverage, intron-spanning reads and intron features are shown as separate tracks.

S67

RNA-seq status of *GPRC6A* in *Galeopterus variegatus*

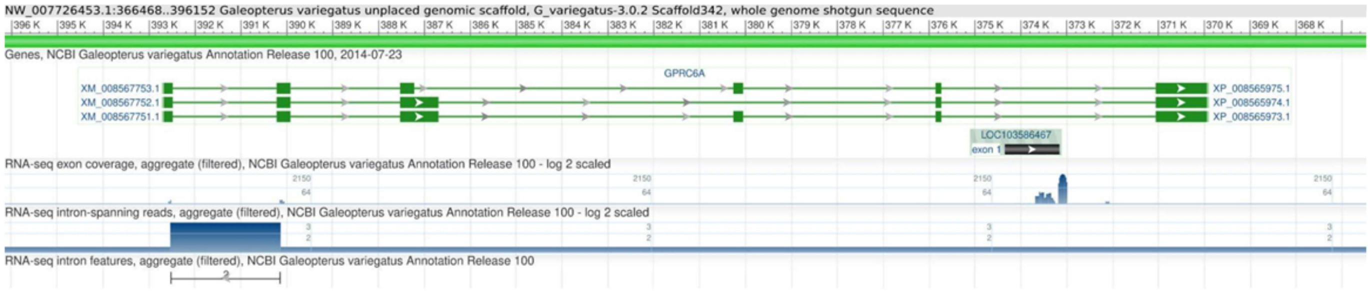


**Supplementary Figure S67. RNA-seq expression in *Galeopterus variegatus:*** This figure displays the RNA-seq expression profile of *GPRC6A* in *Galeopterus variegatus*, visualized using the NCBI Genome Data Viewer. Filtered aggregates of RNA-seq exon coverage, intron-spanning reads and intron features are shown as separate tracks.

S68

RNA-seq status of *GPRC6A* in *Hylobates moloch*

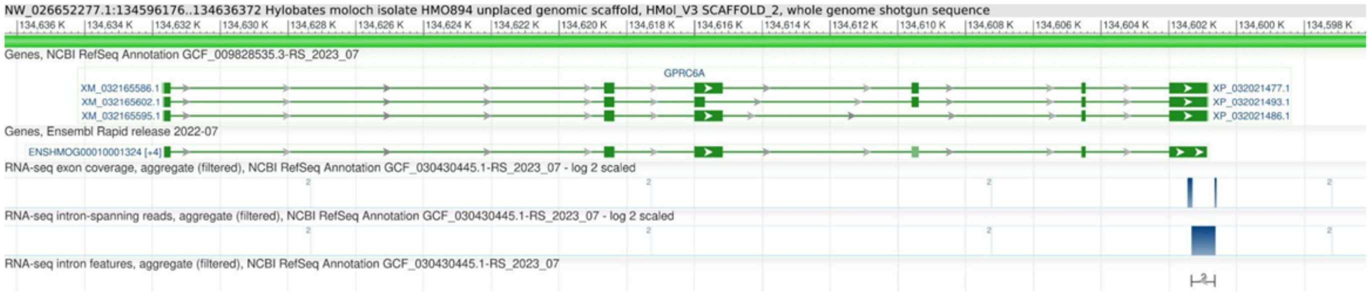


**Supplementary Figure S68. RNA-seq expression in *Hylobates moloch:*** This figure displays the RNA-seq expression profile of *GPRC6A* in *Hylobates moloch*, visualized using the NCBI Genome Data Viewer. Filtered aggregates of RNA-seq exon coverage, intron-spanning reads and intron features are shown as separate tracks.

S69

RNA-seq status of *GPRC6A* in *Microcebus murinus*

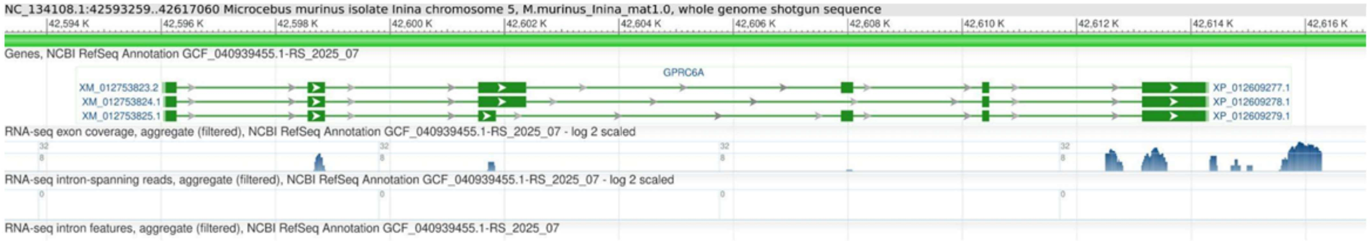


**Supplementary Figure S69. RNA-seq expression in *Microcebus canadensis:*** This figure displays the RNA-seq expression profile of *GPRC6A* in *Microcebus murinus*, visualized using the NCBI Genome Data Viewer. Filtered aggregates of RNA-seq exon coverage, intron-spanning reads and intron features are shown as separate tracks.

S70

RNA-seq status of *GPRC6A* in *Pan troglodytes*

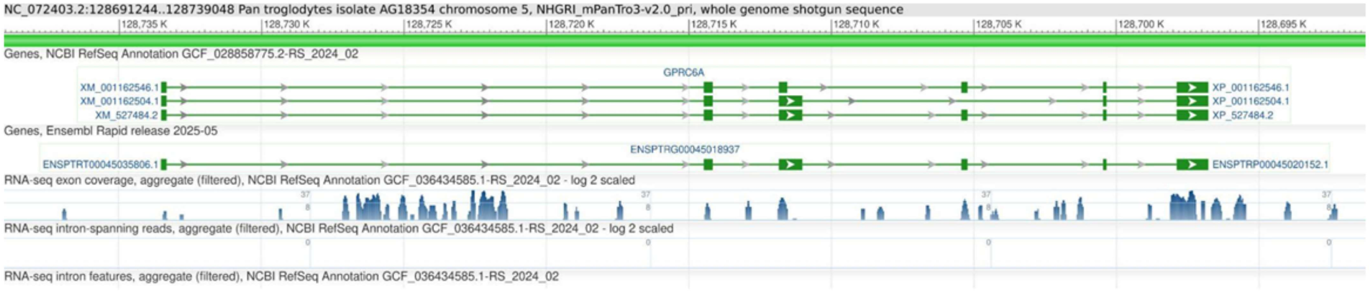


**Supplementary Figure S70. RNA-seq expression in *Pan troglodytes:*** This figure displays the RNA-seq expression profile of *GPRC6A* in *Pan troglodytes*, visualized using the NCBI Genome Data Viewer. Filtered aggregates of RNA-seq exon coverage, intron-spanning reads and intron features are shown as separate tracks.

S71

RNA-seq status of *GPRC6A* in *Papio anubis*

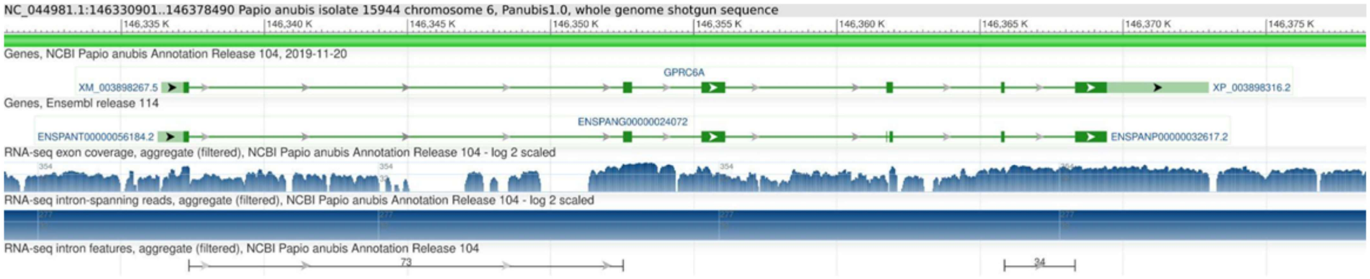


**Supplementary Figure S71. RNA-seq expression in *Papio anubis:*** This figure displays the RNA-seq expression profile of *GPRC6A* in *Papio anubis*, visualized using the NCBI Genome Data Viewer. Filtered aggregates of RNA-seq exon coverage, intron-spanning reads and intron features are shown as separate tracks.

S72

RNA-seq status of *GPRC6A* in *Tupaia chinensis*

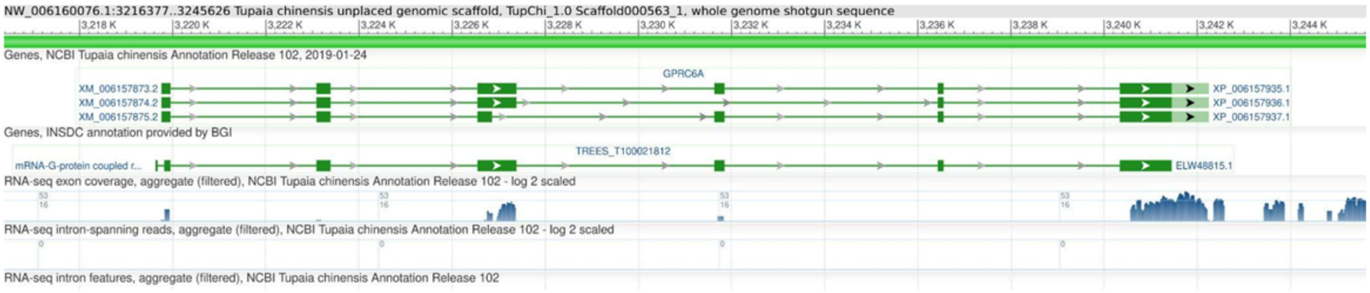


**Supplementary Figure S72. RNA-seq expression in *Tupaia chinensis:*** This figure displays the RNA-seq expression profile of *GPRC6A* in *Tupaia chinensis*, visualized using the NCBI Genome Data Viewer. Filtered aggregates of RNA-seq exon coverage, intron-spanning reads and intron features are shown as separate tracks.

S73

RNA-seq status of *GPRC6A* in *Manis javanica*

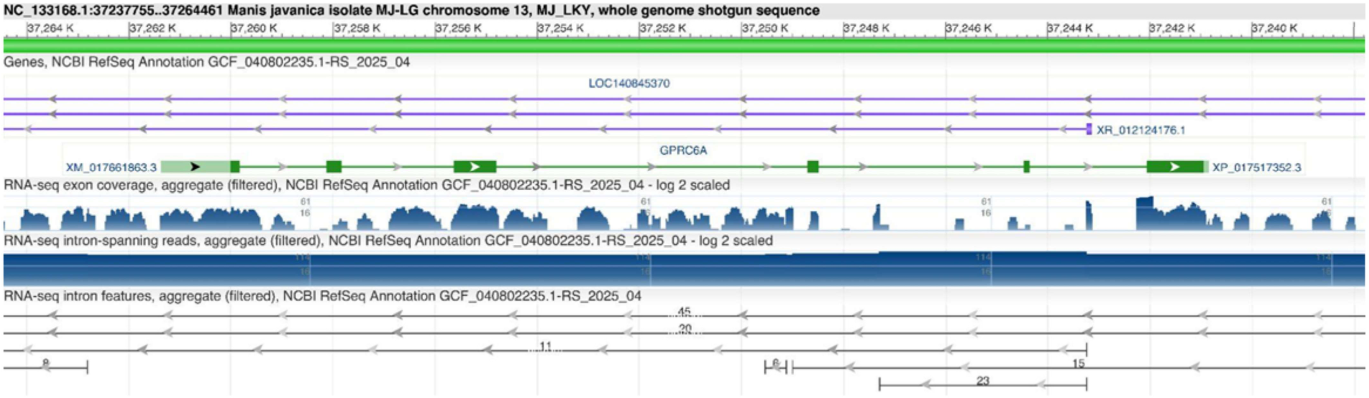


**Supplementary Figure S73. RNA-seq expression in *Manis javanica:*** This figure displays the RNA-seq expression profile of *GPRC6A* in *Manis javanica*, visualized using the NCBI Genome Data Viewer. Filtered aggregates of RNA-seq exon coverage, intron-spanning reads and intron features are shown as separate tracks.

S74

RNA-seq status of *GPRC6A* in *Manis pentadactyla*

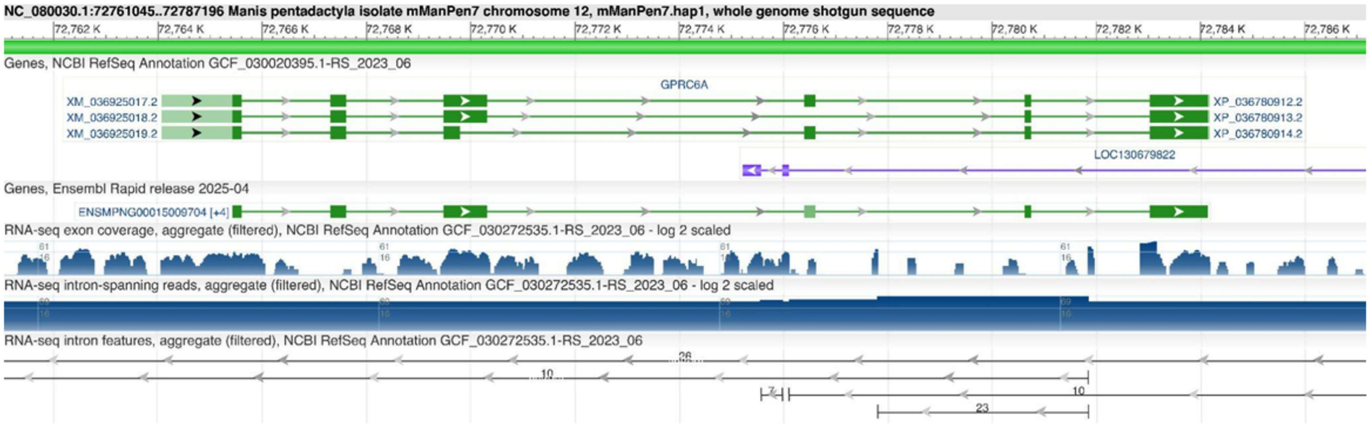


**Supplementary Figure S74. RNA-seq expression in *Manis pentadactyla:*** This figure displays the RNA-seq expression profile of *GPRC6A* in *Manis pentadactyla*, visualized using the NCBI Genome Data Viewer. Filtered aggregates of RNA-seq exon coverage, intron-spanning reads and intron features are shown as separate tracks.

S75

RNA-seq status of *GPRC6A* in *Pteropus giganteus*

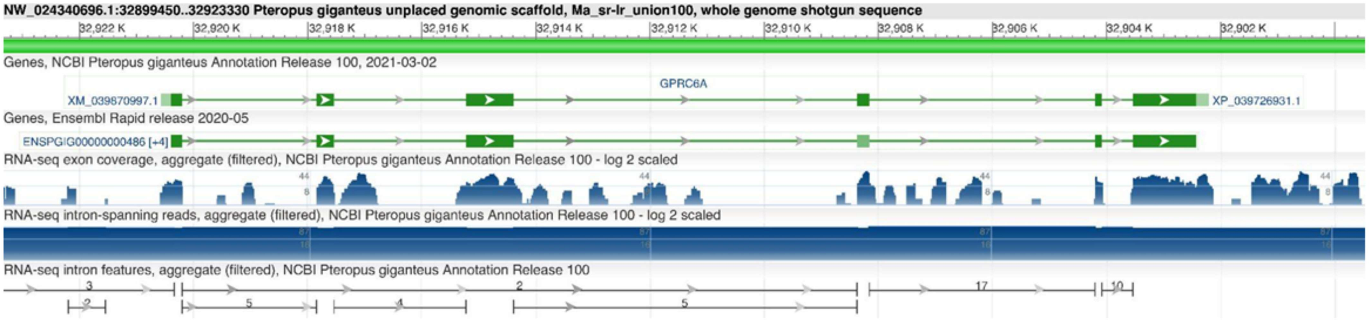


**Supplementary Figure S75. RNA-seq expression in *Pteropus giganteus:*** This figure displays the RNA-seq expression profile of *GPRC6A* in *Pteropus giganteus*, visualized using the NCBI Genome Data Viewer. Filtered aggregates of RNA-seq exon coverage, intron-spanning reads and intron features are shown as separate tracks.

S76

RNA-seq status of *GPRC6A* in *Rousettus aegyptiacus*

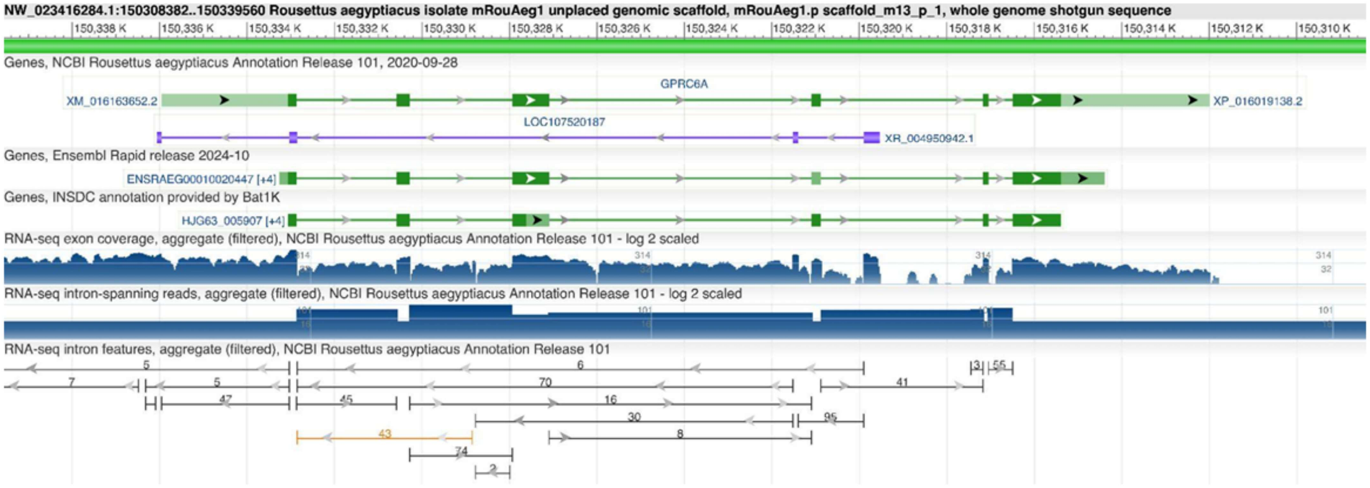


**Supplementary Figure S76. RNA-seq expression in *Rousettus aegyptiacus:*** This figure displays the RNA-seq expression profile of *GPRC6A* in *Rousettus aegyptiacus canadensis*, visualized using the NCBI Genome Data Viewer. Filtered aggregates of RNA-seq exon coverage, intron-spanning reads and intron features are shown as separate tracks.

S77

RNA-seq status of *GPRC6A* in *Sturnira hondurensis*

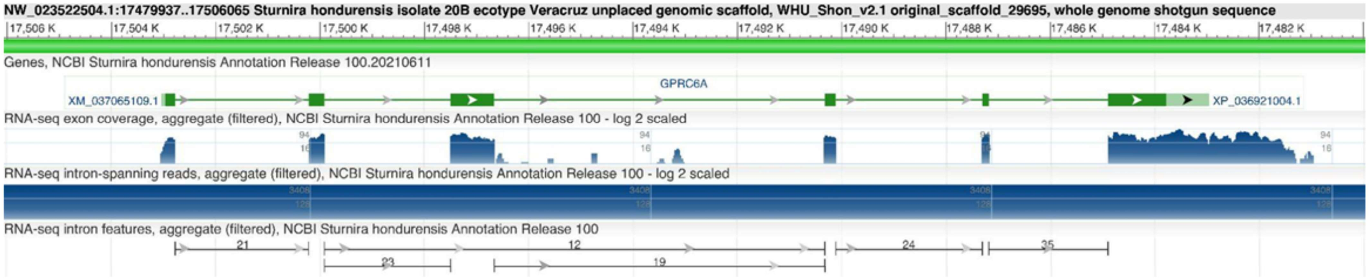


**Supplementary Figure S77. RNA-seq expression in *Sturnira hondurensis:*** This figure displays the RNA-seq expression profile of *GPRC6A* in *Sturnira hondurensis*, visualized using the NCBI Genome Data Viewer. Filtered aggregates of RNA-seq exon coverage, intron-spanning reads and intron features are shown as separate tracks.

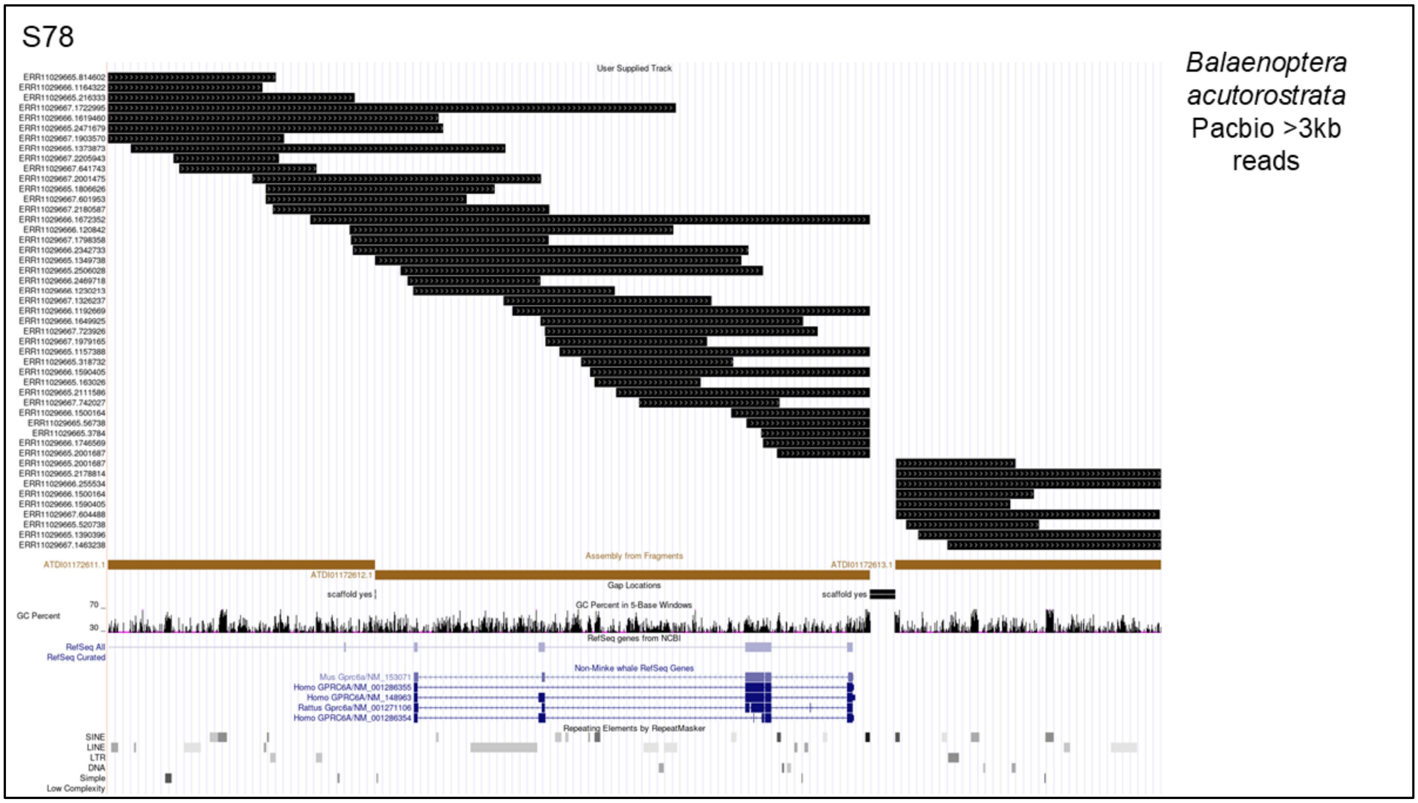


**Supplementary Figure S78. PacBio Read Coverage of the *GPRC6A* Region in *Balaenoptera acutorostrata***: This figure shows PacBio reads spanning the expected *GPRC6A* region in *Balaenoptera acutorostrata*. The annotations at the bottom represent the RefSeq gene model for *GPRC6A*. Reads longer than 3 kb cover the entire region, confirming that the assembly is intact and supporting the conclusion that the loss of *GPRC6A* in *Balaenoptera acutorostrata* is genuine.

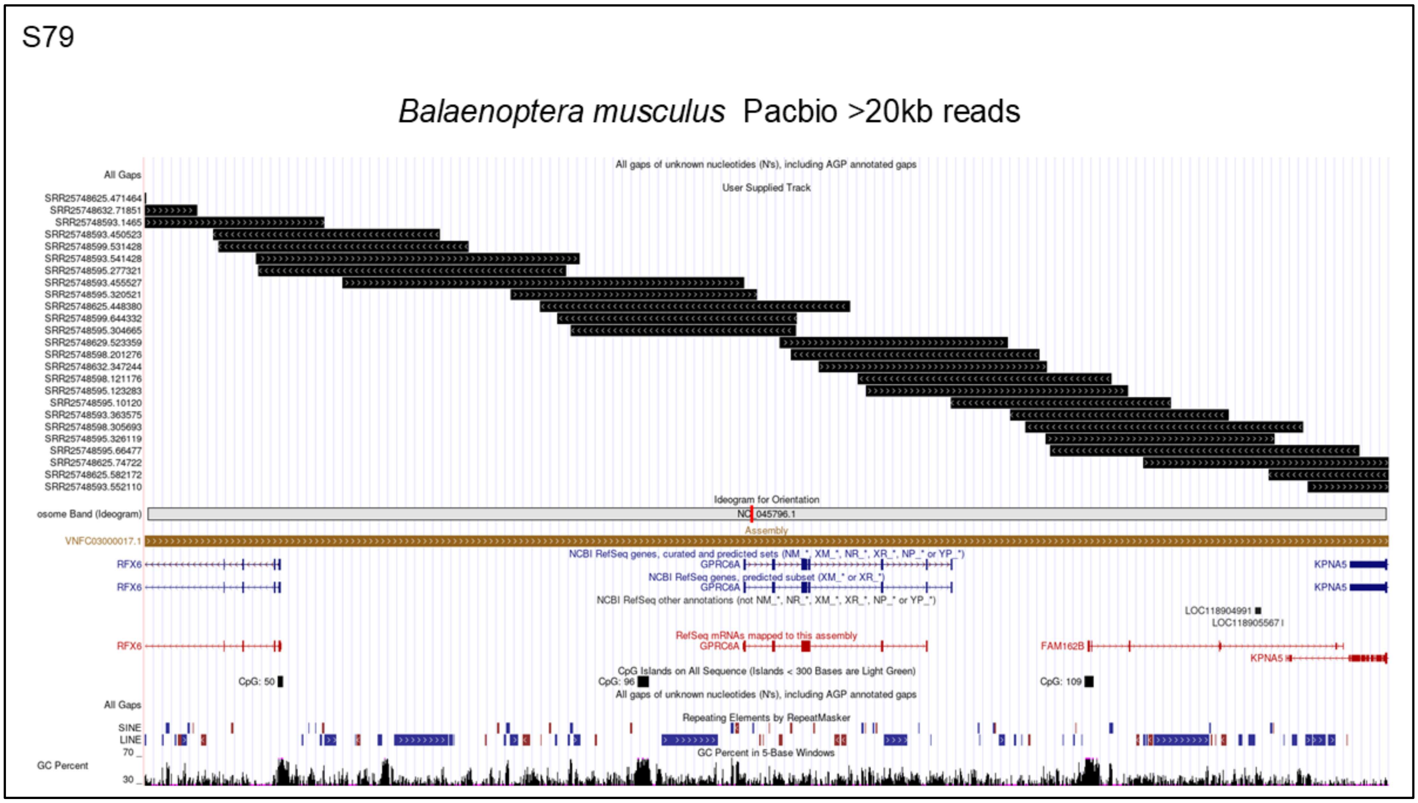


**Supplementary Figure S79. PacBio Read Coverage of the *GPRC6A* Region in *Balaenoptera musculus***: This figure shows PacBio reads spanning the expected *GPRC6A* region in *Balaenoptera musculus*. The annotations at the bottom represent the RefSeq gene model for *GPRC6A*. Reads longer than 20 kb cover the entire region, confirming that the assembly is intact and supporting the conclusion that the loss of *GPRC6A* in *Balaenoptera musculus* is genuine.

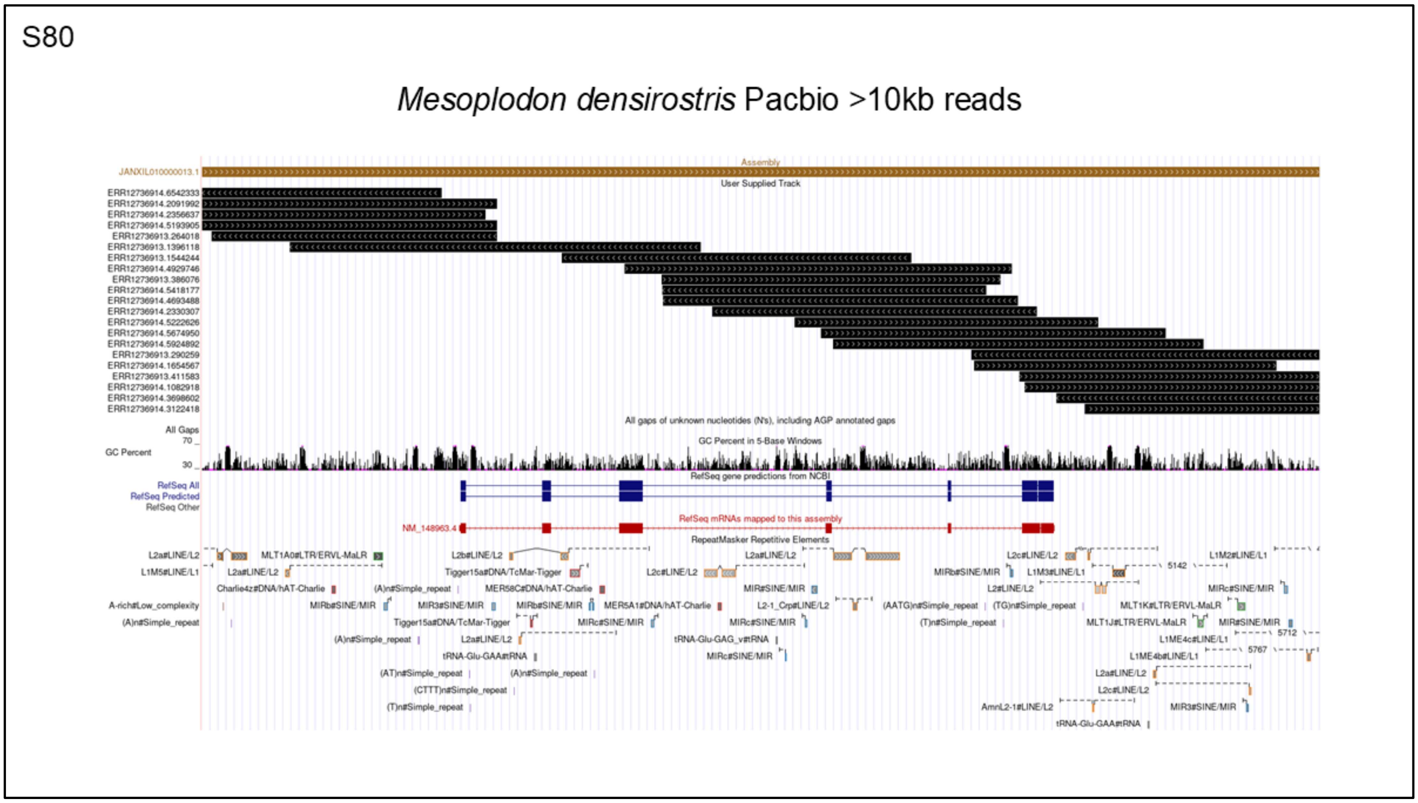


**Supplementary Figure S80. PacBio Read Coverage of the *GPRC6A* Region in *Mesoplodon densirostris***: This figure shows PacBio reads spanning the expected *GPRC6A* region in *Mesoplodon densirostris*. The annotations at the bottom represent the RefSeq gene model for *GPRC6A*. Reads longer than 10 kb cover the entire region, confirming that the assembly is intact and supporting the conclusion that the loss of *GPRC6A* in *Mesoplodon densirostris* is genuine.

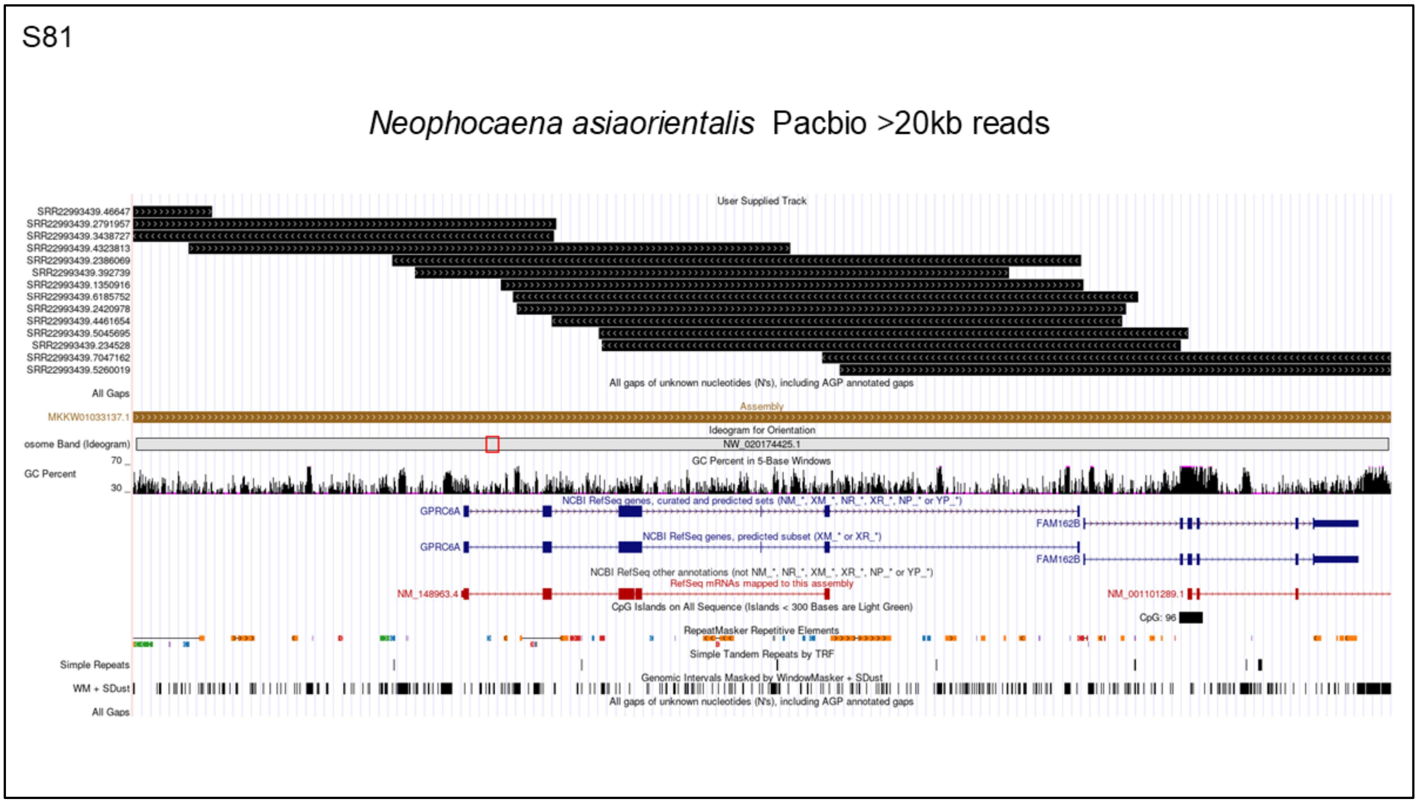


**Supplementary Figure S81. PacBio Read Coverage of the *GPRC6A* Region in *Neophocaena asiaorientalis*:** This figure shows PacBio reads spanning the expected *GPRC6A* region in *Neophocaena asiaorientalis*. The annotations at the bottom represent the RefSeq gene model for *GPRC6A*. Reads longer than 20 kb cover the entire region, confirming that the assembly is intact and supporting the conclusion that the loss of *GPRC6A* in *Neophocaena asiaorientalis* is genuine.

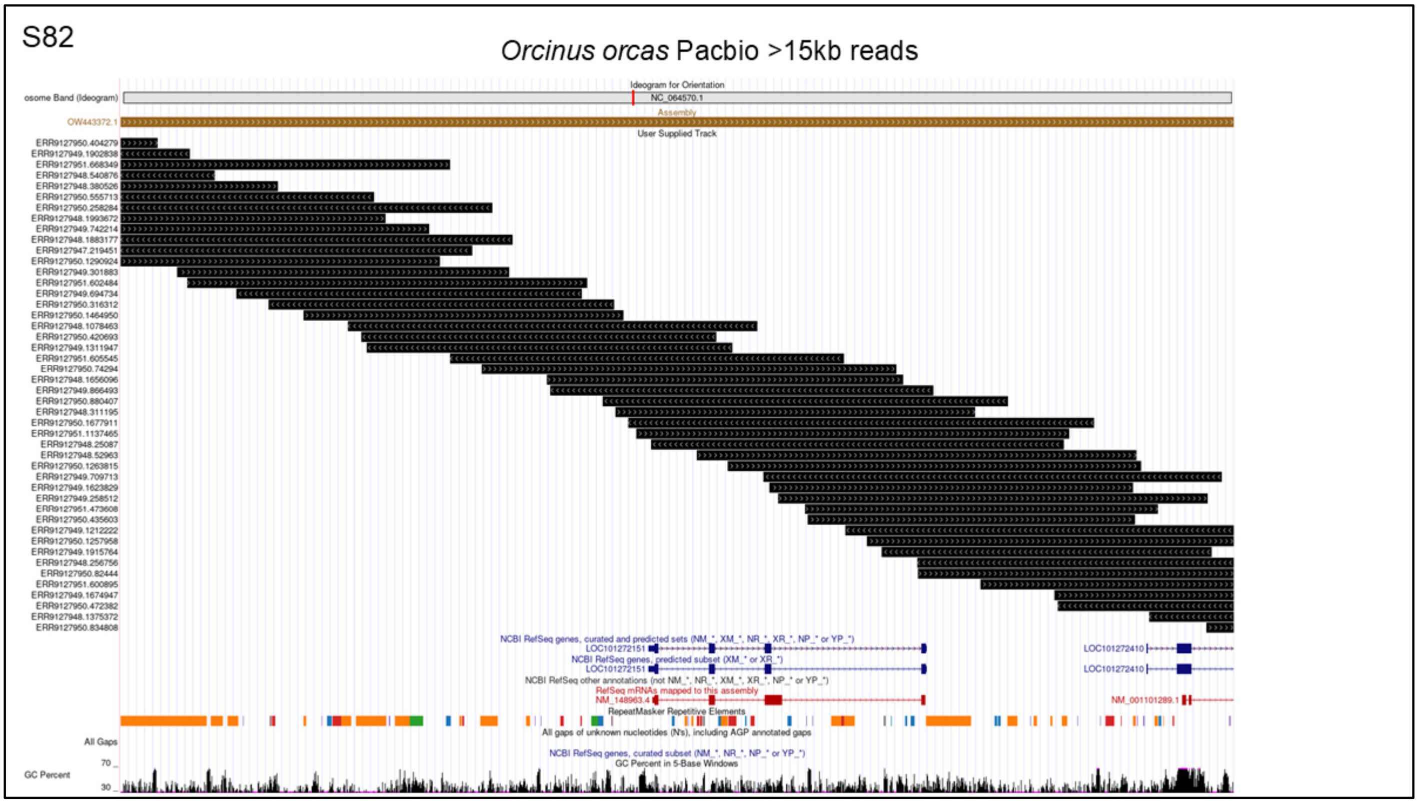


**Supplementary Figure S82. PacBio Read Coverage of the *GPRC6A* Region in *Orcinus orcas***: This figure shows PacBio reads spanning the expected *GPRC6A* region in *Orcinus orcas*. The annotations at the bottom represent the RefSeq gene model for *GPRC6A*. Reads longer than 15 kb cover the entire region, confirming that the assembly is intact and supporting the conclusion that the loss of *GPRC6A* in *Orcinus orcas* is genuine.

S83

*Tragulus javanicus* with *Sus scrofa* reference. Chain file.

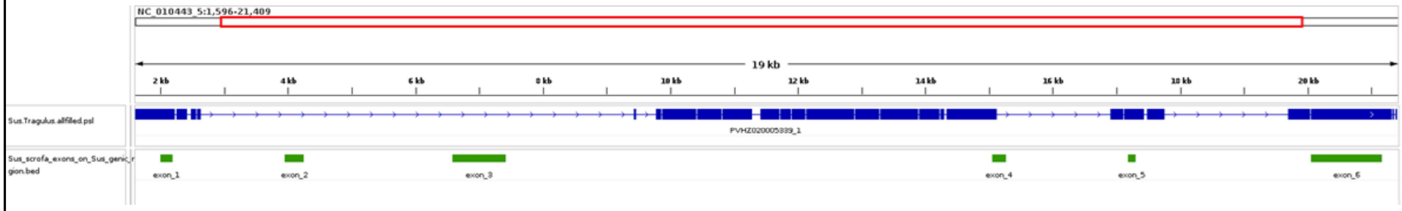


**Supplementary Figure S83:Chain file of *Tragulus javanicus* aligned with the genome of *Sus scrofa*** visualized in IGV. The chain file of *Tragulus javanicus* is shown in blue. The exons of the *GPRC6A* gene on the *Sus scrofa* genome are highlighted in green, indicating the alignment of *Tragulus javanicus* sequences with corresponding genomic regions of *Sus scrofa*.

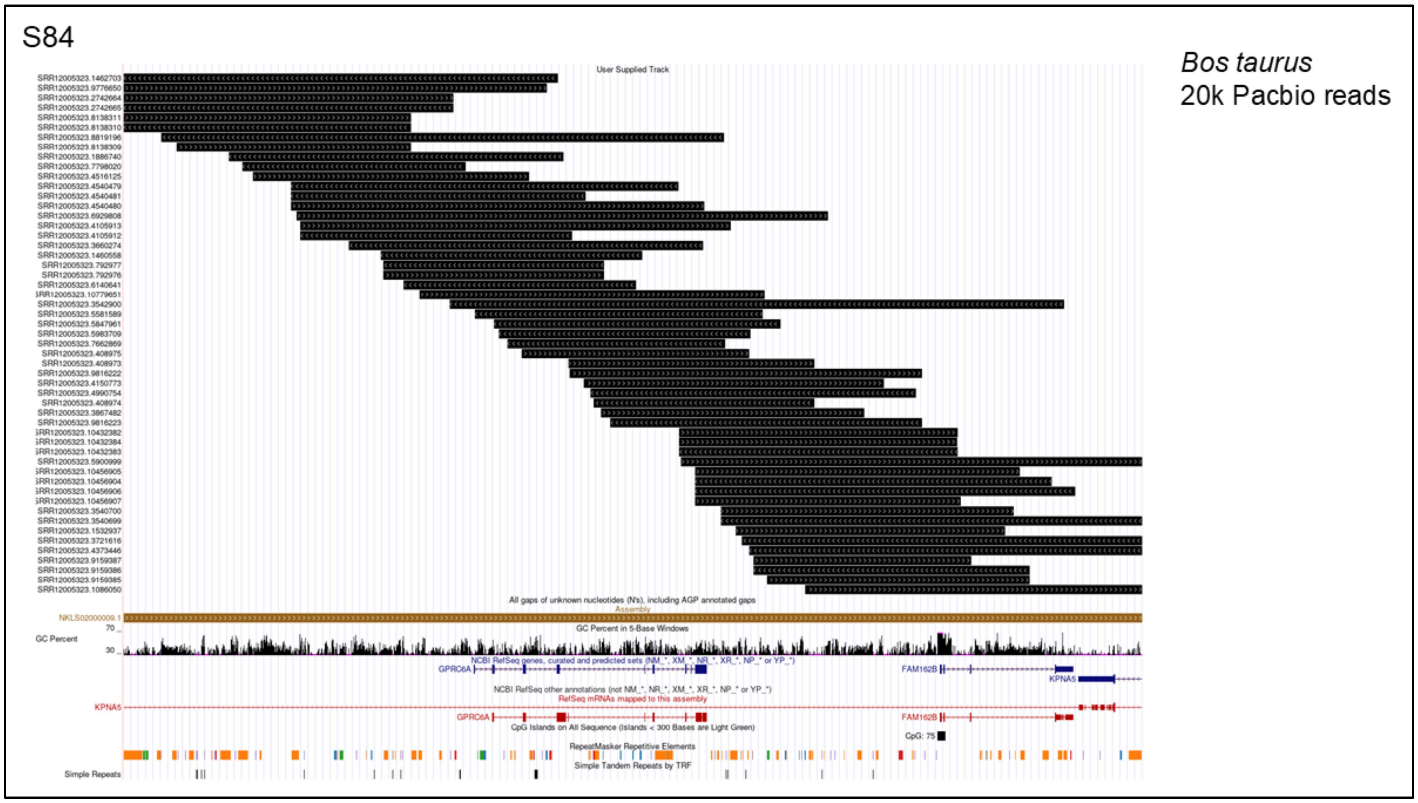


**Supplementary Figure S84. PacBio Read Coverage of the *GPRC6A* Region in *Bos taurus*:** This figure shows PacBio reads spanning the expected *GPRC6A* region in *Bos taurus*. The annotations at the bottom represent the RefSeq gene model for *GPRC6A* based on the *Homo sapiens* reference. Reads longer than 20 kb cover the entire region, confirming that the assembly is intact and supporting the conclusion that the loss of *GPRC6A* in *Bos taurus* is genuine.

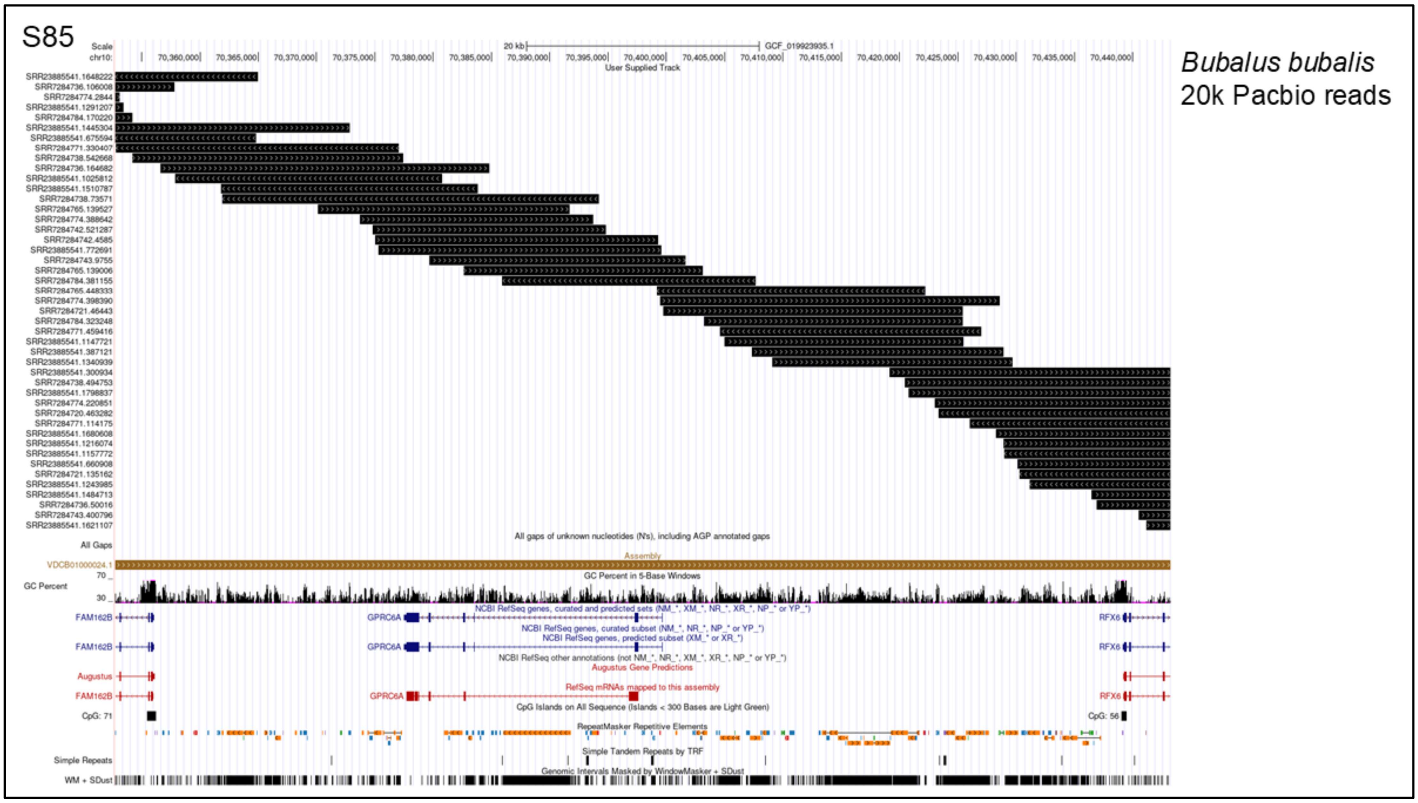


**Supplementary Figure S85. PacBio Read Coverage of the *GPRC6A* Region in *Bubalus bubalis*:** This figure shows PacBio reads spanning the expected *GPRC6A* region in *Bubalus bubalis*. The annotations at the bottom represent the RefSeq gene model for *GPRC6A*. Reads longer than 20 kb cover the entire region, confirming that the assembly is intact and supporting the conclusion that the loss of *GPRC6A* in *Bubalus bubalis* is genuine.

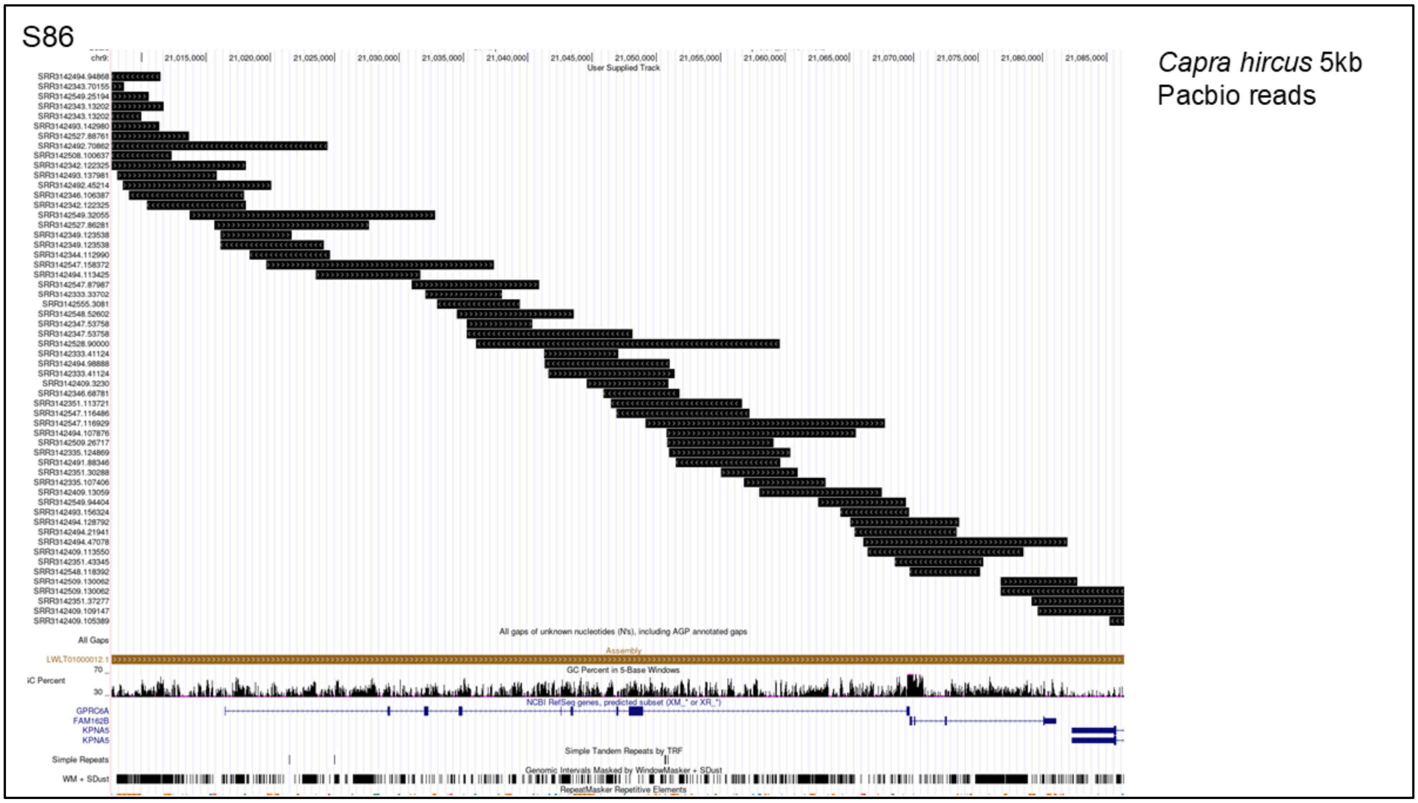


**Supplementary Figure S86. PacBio Read Coverage of the *GPRC6A* Region in *Capra hircus*:** This figure shows PacBio reads spanning the expected *GPRC6A* region in *Capra hircus*. The annotations at the bottom represent the RefSeq gene model for *GPRC6A*. Reads longer than 5 kb cover the entire region, confirming that the assembly is intact and supporting the conclusion that the loss of *GPRC6A* in *Capra hircus* is genuine.

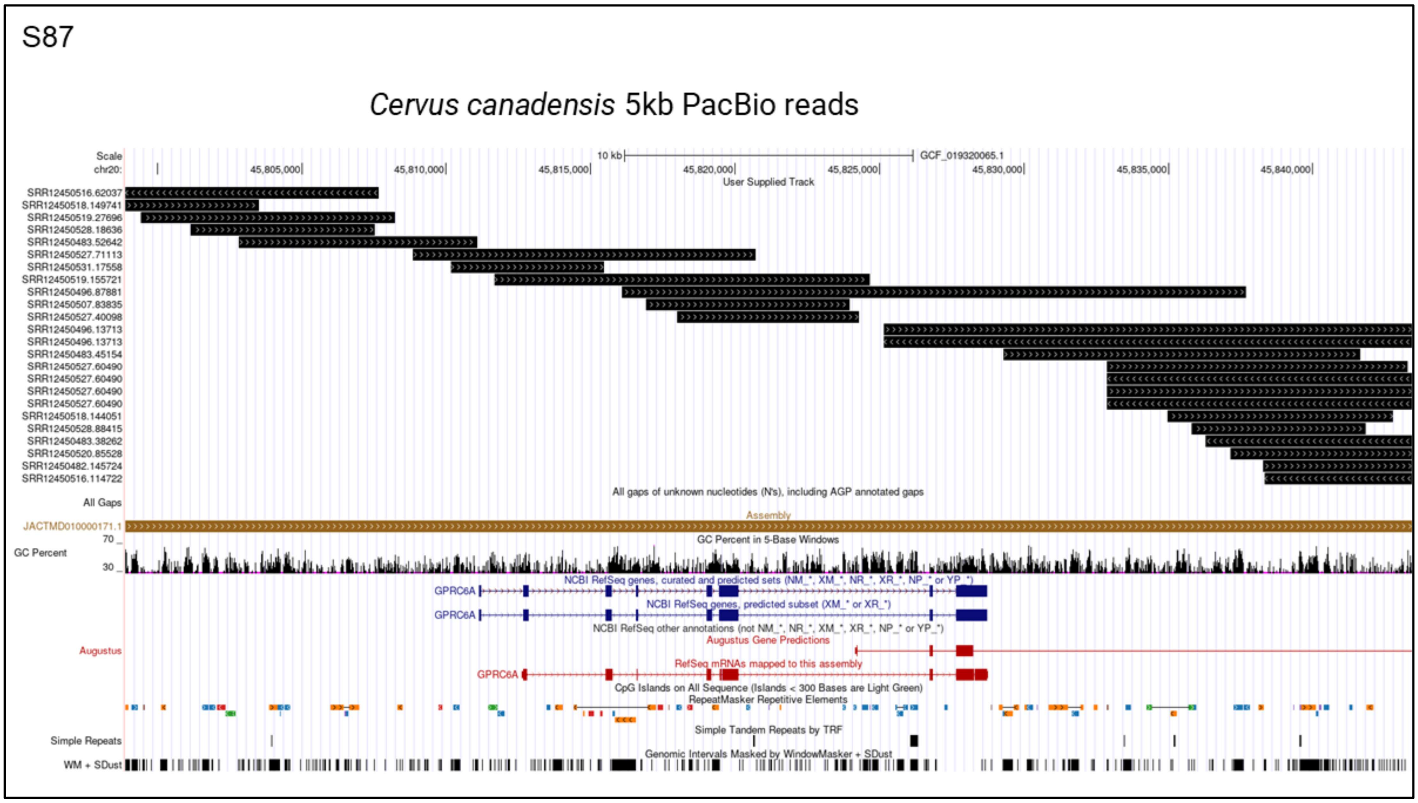


**Supplementary Figure S87. PacBio Read Coverage of the *GPRC6A* Region in *Cervus canadensis***: This figure shows PacBio reads spanning the expected *GPRC6A* region in *Cervus canadensis*. The annotations at the bottom represent the RefSeq gene model for *GPRC6A*. Reads longer than 5 kb cover the entire region, confirming that the assembly is intact and supporting the conclusion that the loss of *GPRC6A* in *Cervus canadensis* is genuine.

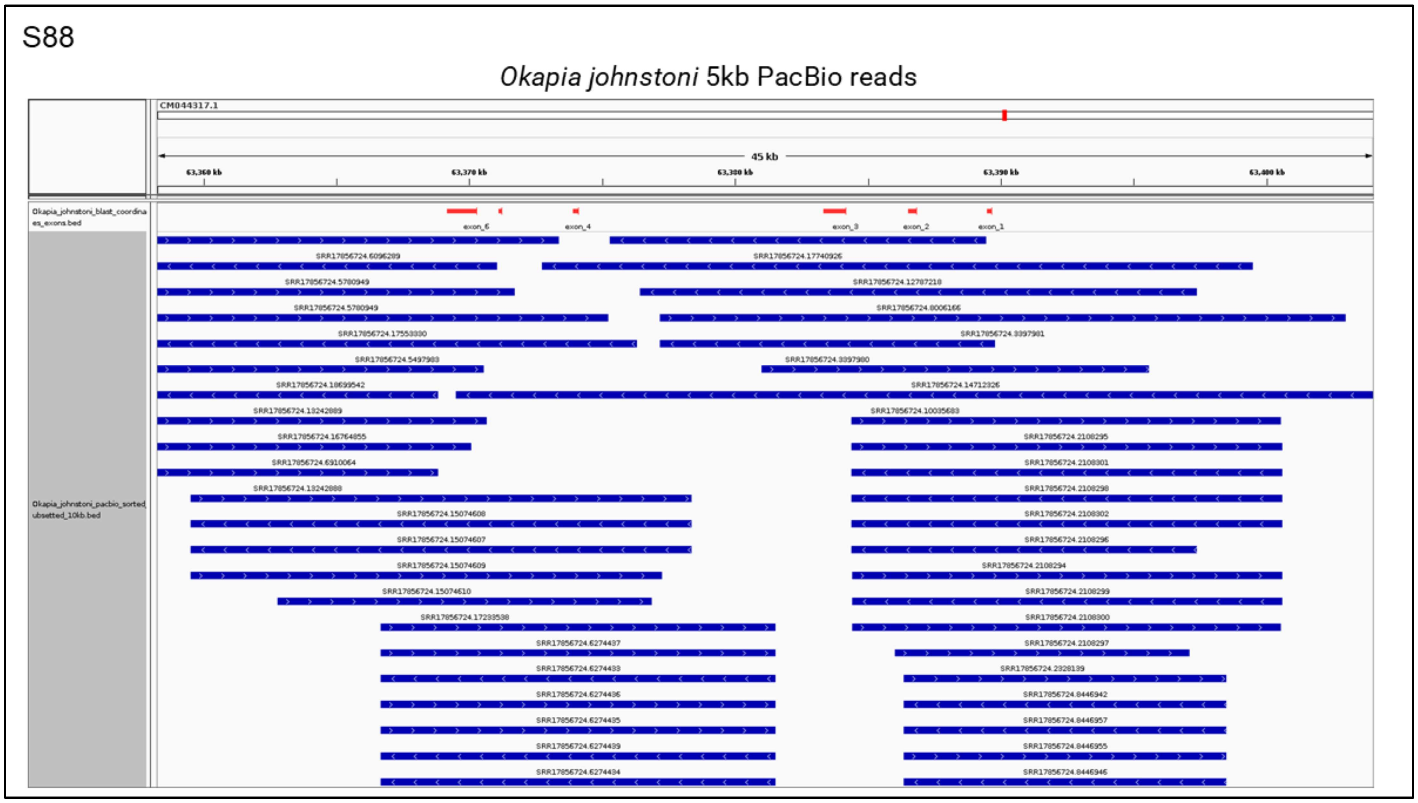


**Supplementary Figure S88. IGV Visualization of PacBio Reads Aligned to the *Okapia johnstoni* genome:** This figure presents an IGV visualization of PacBio reads longer than 5 kb from *Okapia johnstoni* aligned to the *Okapia johnstoni* genome. The putative locations of exons on the genome are highlighted in red. The complete coverage of this region by the reads indicates that the assembly is intact and accurate.

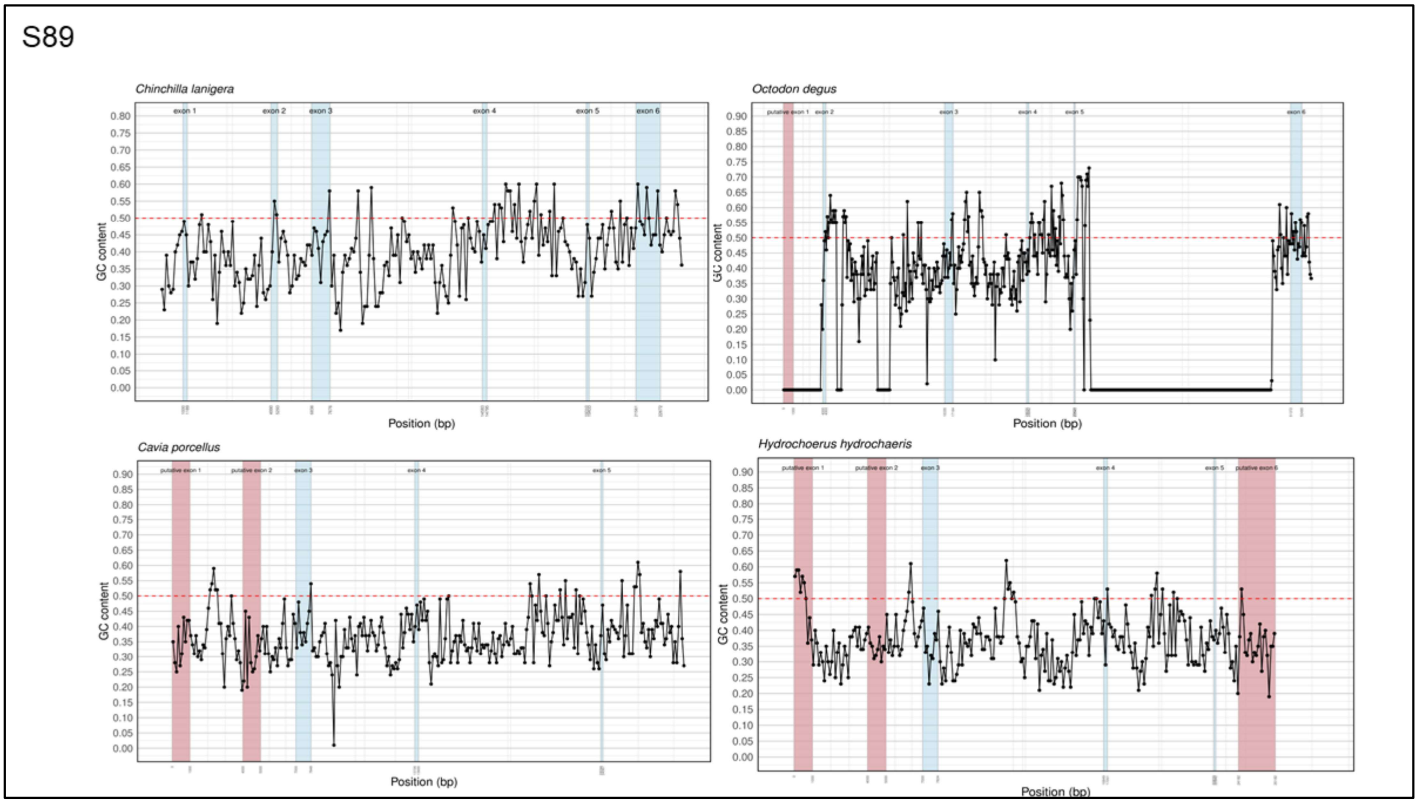


**Supplementary Figure S89. Figure showing GC content of four representative caviomorph rodents in 100 bp windows**: The x-axis represents the base pair (bp) positions, and the y-axis indicates the mean of GC base pairs within each 100 bp window. The exonic locations of the *GPRC6A* gene are highlighted in light blue while the putative location of the exons are in light red.

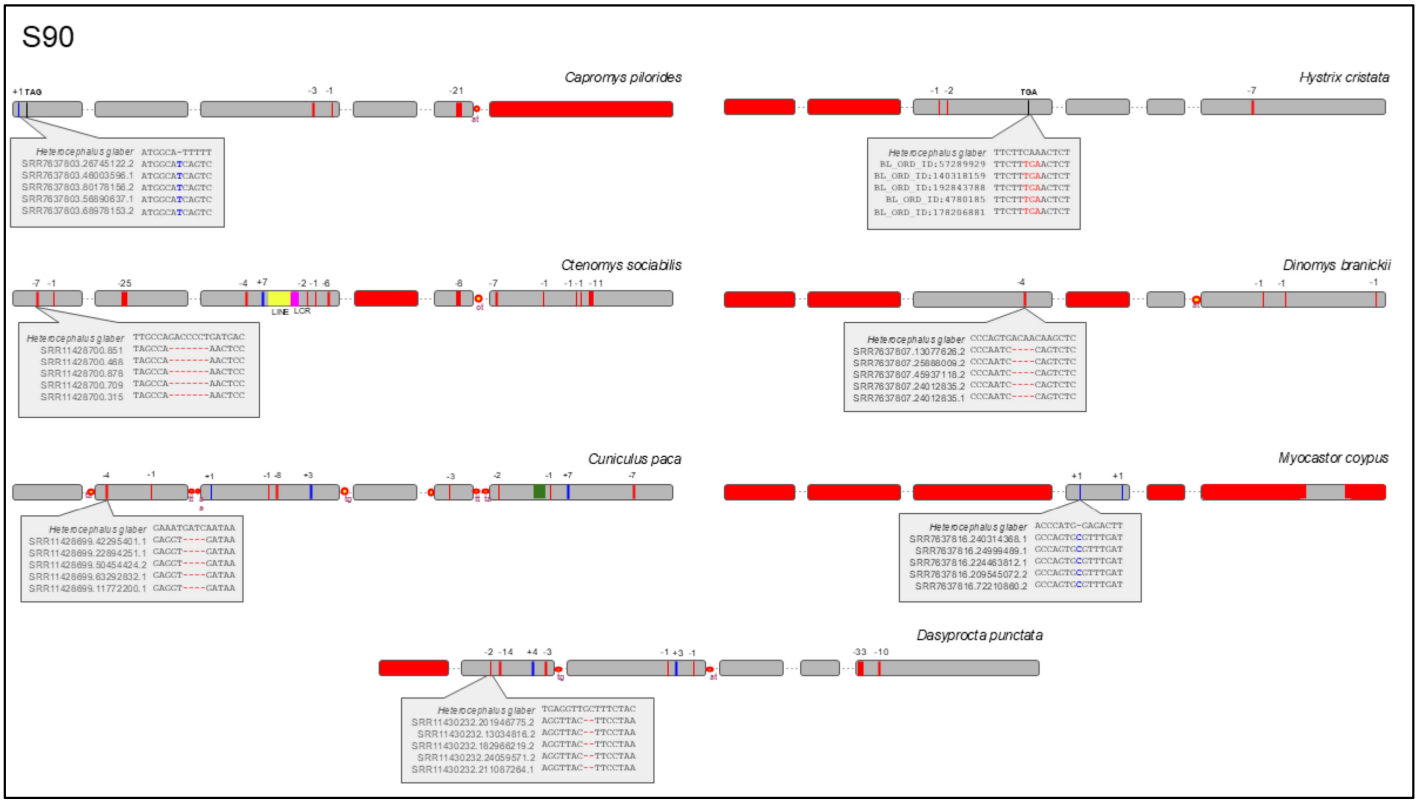


**Supplementary Figure S90: Visualization of Loss Events in *GPRC6A* Among Caviomorph Rodents:** This figure illustrates the identified loss events and the first occurrences of stop codons in the *GPRC6A* gene among caviomorph rodents. Exons are represented by grey boxes, while exons that could not be retrieved using BLAST are shown in red boxes. Insertions are depicted as blue bars, and deletions are indicated by red bars. Splice site disruptions are marked with red and yellow circles. Bright yellow represents LINE insertions, and magenta indicates low complexity repeats. Green bars signify simple repeats. The zoomed-in sections highlight Illumina SRA reads aligned to the *Heterocephalus glaber* reference genome, emphasizing frame-disrupting changes.

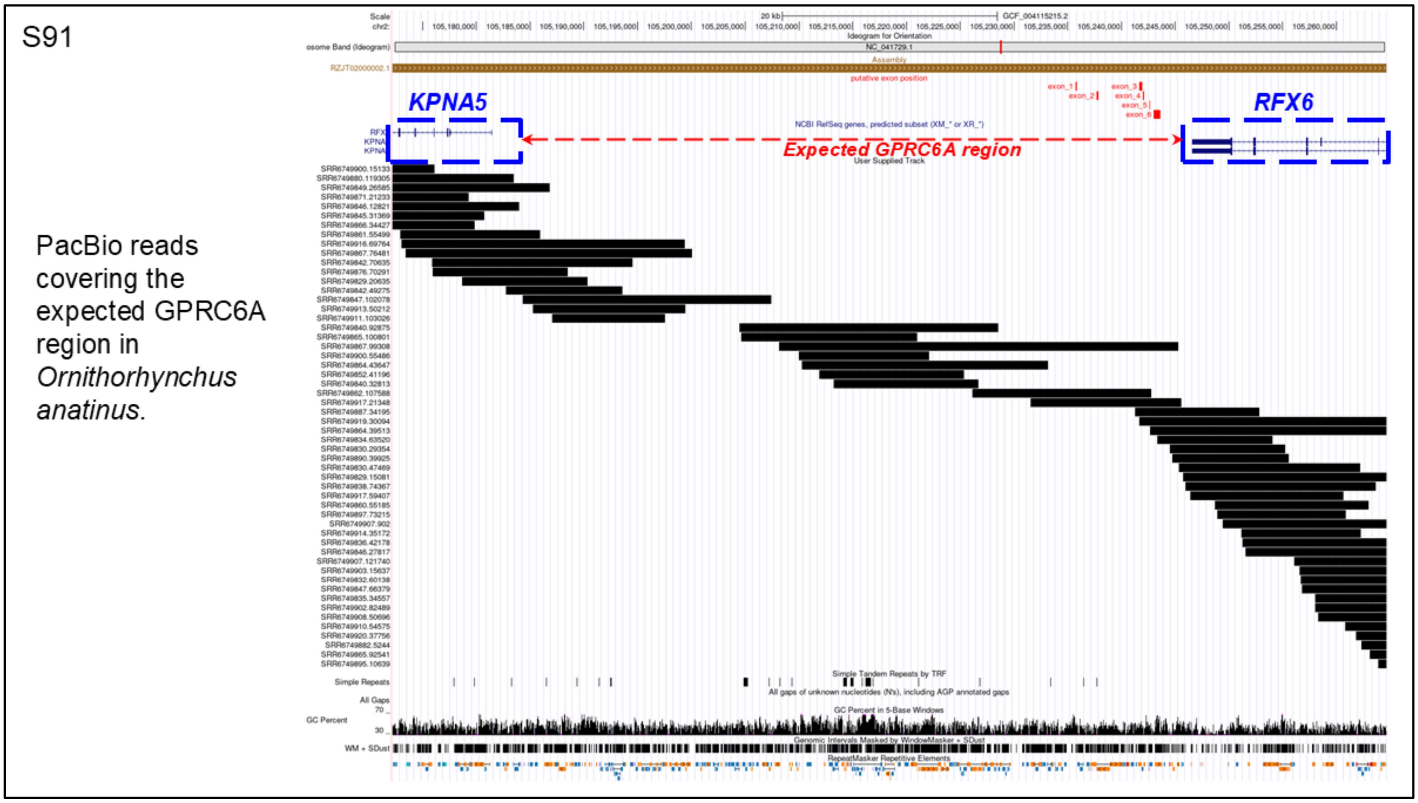


**Supplementary Figure S91. PacBio Read Coverage of the *GPRC6A* Region in *Ornithorhynchus anatinus***: This figure illustrates PacBio reads spanning the expected *GPRC6A* region in *Ornithorhynchus anatinus*. The two blue boxes indicate the presence of adjacent genes, while the red arrows mark the expected *GPRC6A* region. Reads longer than 10 kb cover the entire region, confirming that the assembly is intact and supporting the conclusion that the loss of *GPRC6A* in *Ornithorhynchus* is genuine.

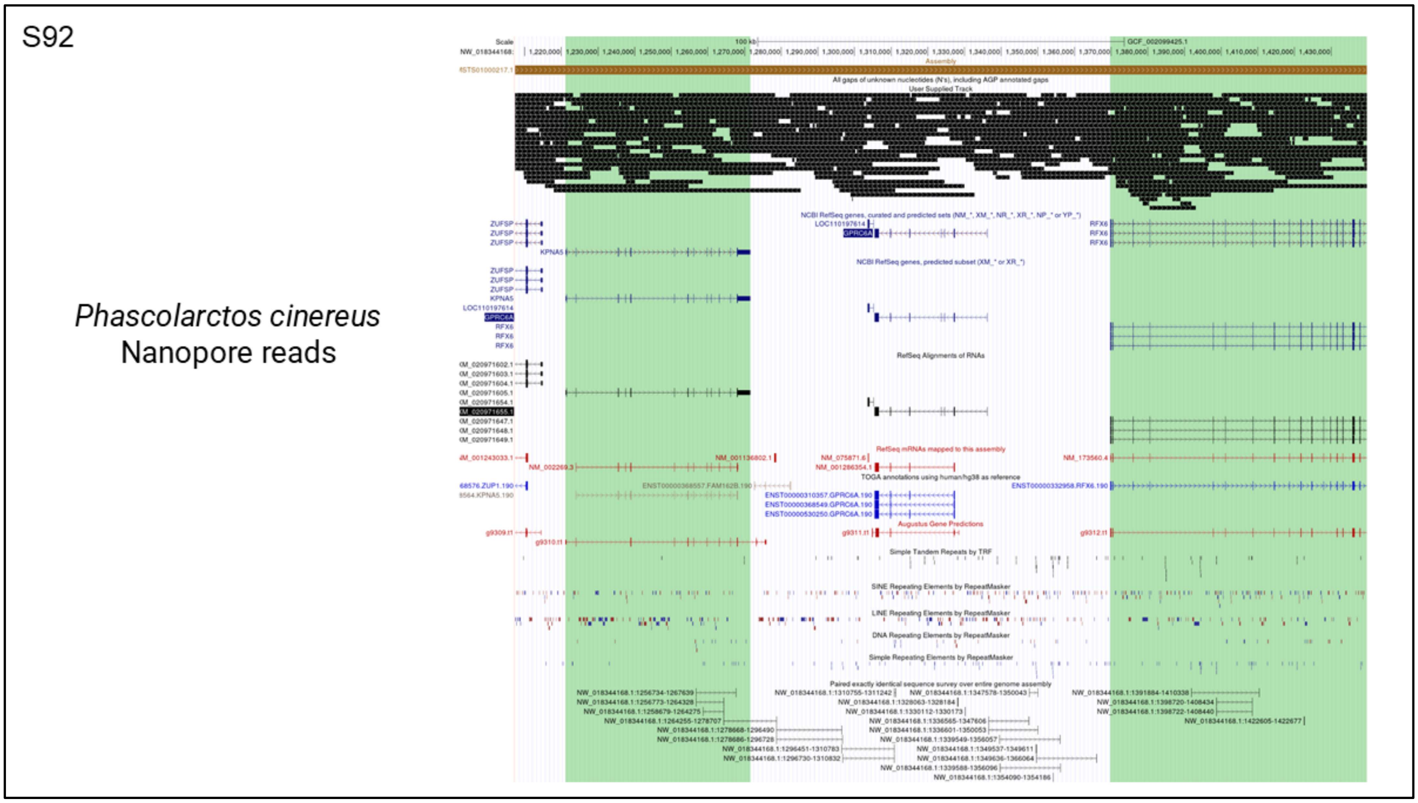


**Supplementary Figure S92. Nanopore Read Coverage of the *GPRC6A* Region in *Phascolarctos cinereus***: This figure shows Nanopore reads spanning the expected *GPRC6A* region in *Phascolarctos cinereus* visualized in UCSC genome browser. Reads are stacked in a compressed view. The annotations at the bottom represent the RefSeq gene model for *GPRC6A* based on the *Homo sapiens* reference. Reads longer than 5 kb cover the entire region, confirming that the assembly is intact and supporting the conclusion that the loss of *GPRC6A* in *Phascolarctos cinereus* is genuine.

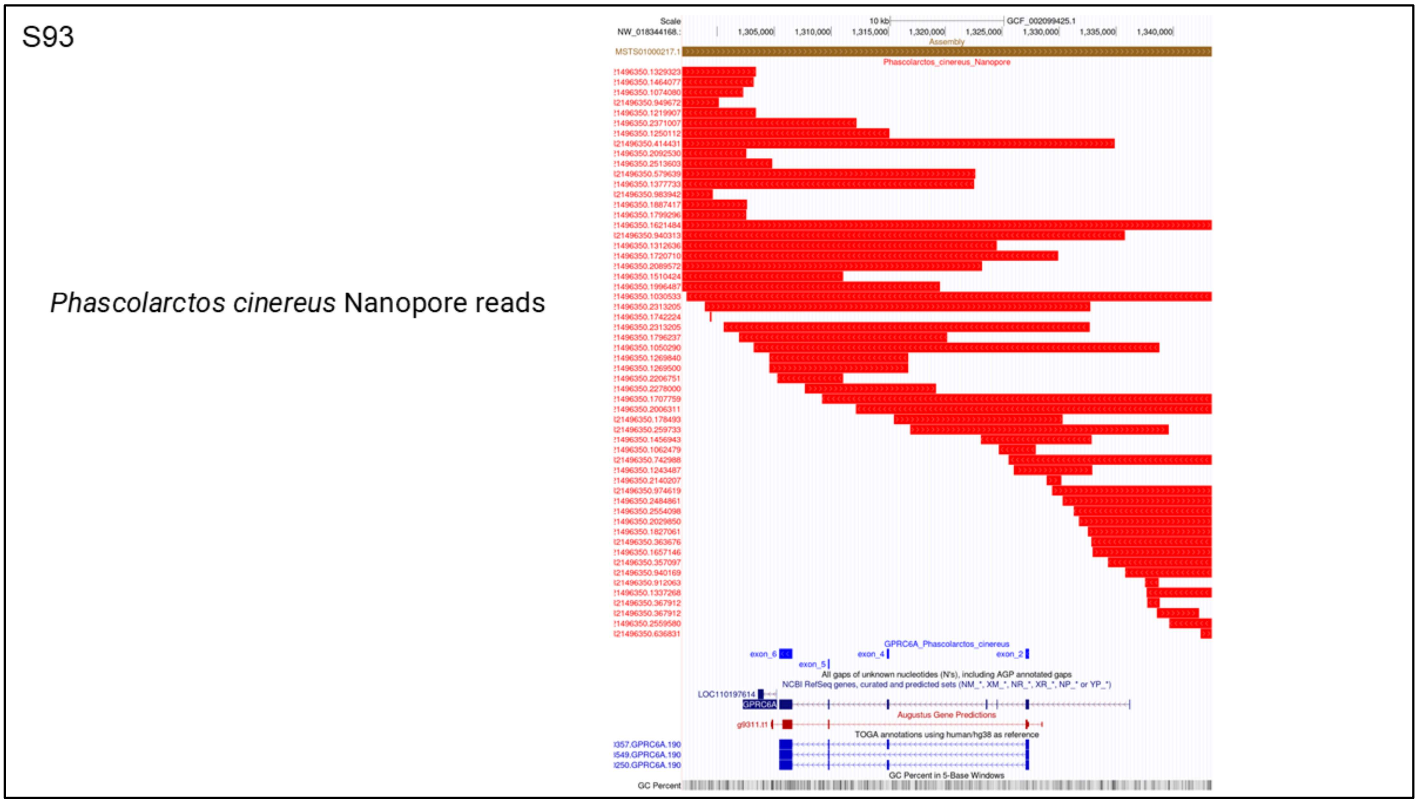


**Supplementary Figure S93. Nanopore Read Coverage of the *GPRC6A* region in *Phascolarctos cinereus***: This figure shows Nanopore reads spanning the expected *GPRC6A* region in *Phascolarctos cinereus* visualized in UCSC genome browser. Reads are stacked in an expanded view. The annotations at the bottom represent the RefSeq gene model for *GPRC6A*. Reads longer than 5 kb cover the entire region, confirming that the assembly is intact and supporting the conclusion that the loss of *GPRC6A* in *Phascolarctos cinereus* is genuine.

S94

*Phascolarctos cinereus* Chain file aligned with *Vombatus ursinus*

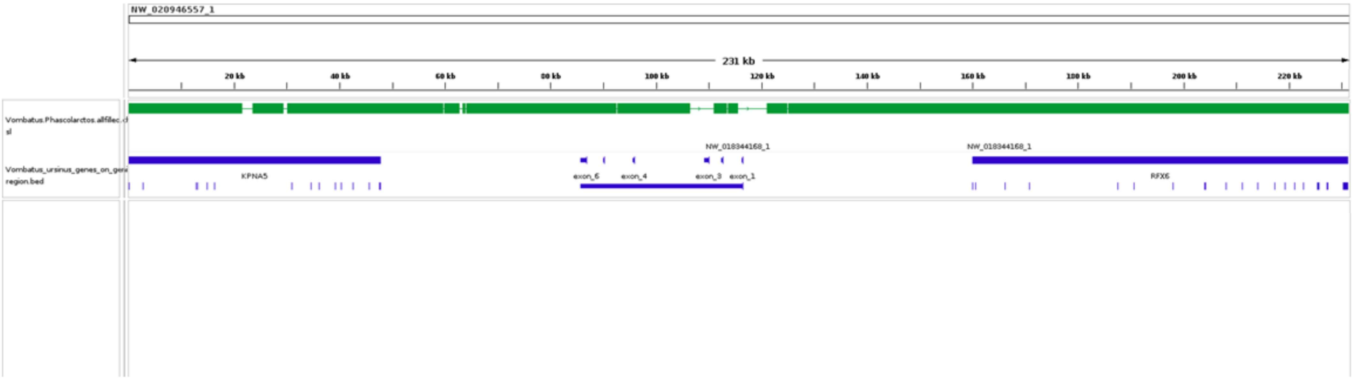


**Supplementary Figure S94:Chain file of *Phascolarctos cinereus* aligned with the genome of *Vombatus ursinus*** visualized in IGV. The chain file of *Phascolarctos cinereus* is shown in green. The exons of the *GPRC6A* gene on the *Vombatus ursinus* genome are highlighted in blue, indicating the alignment of *Phascolarctos cinereus* sequences with corresponding genomic regions of *Vombatus ursinus.*

S95

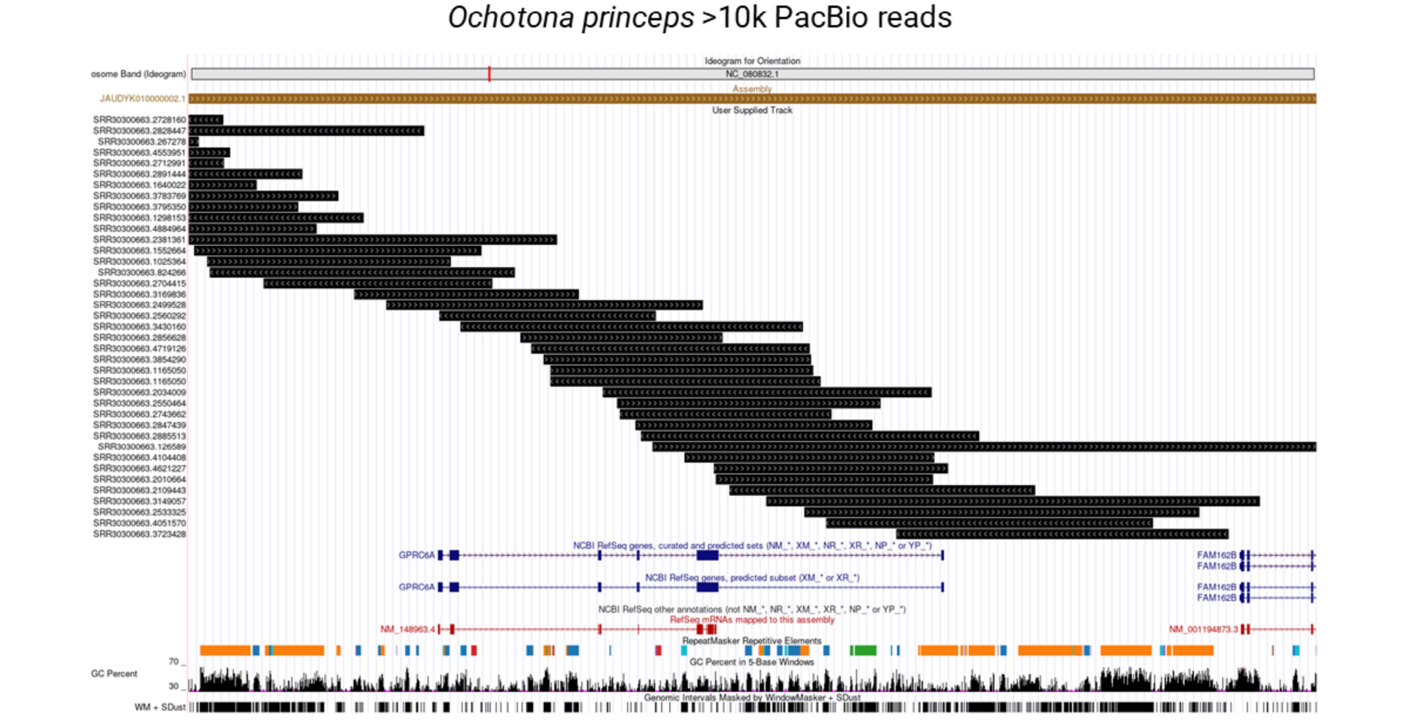


**Supplementary Figure S95. PacBio Read Coverage of the *GPRC6A* region in *Ochotona princeps***: This figure shows PacBio reads spanning the expected *GPRC6A* region in *Ochotona princeps* visualized in UCSC genome browser. The annotations at the bottom represent the RefSeq gene model for *GPRC6A*. Reads longer than 10 kb cover the entire region, confirming that the assembly is intact and supporting the conclusion that the loss of *GPRC6A* in *Ochotona princeps* is genuine.

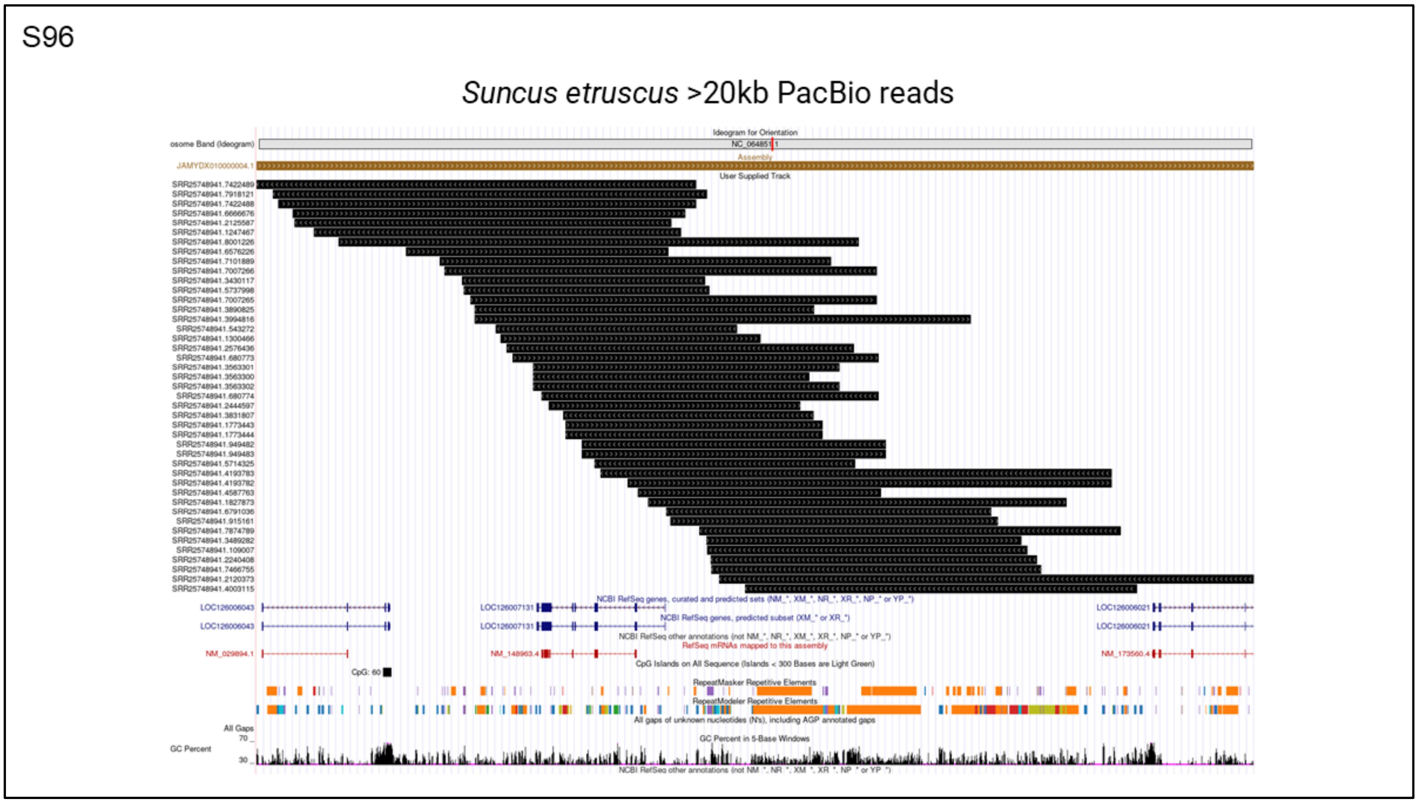


**Supplementary Figure S96. PacBio Read Coverage of the *GPRC6A* region in *Suncus etruscus***: This figure shows PacBio reads spanning the expected *GPRC6A* region in *Suncus etruscus* visualized in UCSC genome browser. The annotations at the bottom represent the RefSeq gene model for *GPRC6A*. Reads longer than 20 kb cover the entire region, confirming that the assembly is intact and supporting the conclusion that the loss of *GPRC6A* in *Suncus etruscus* is genuine.

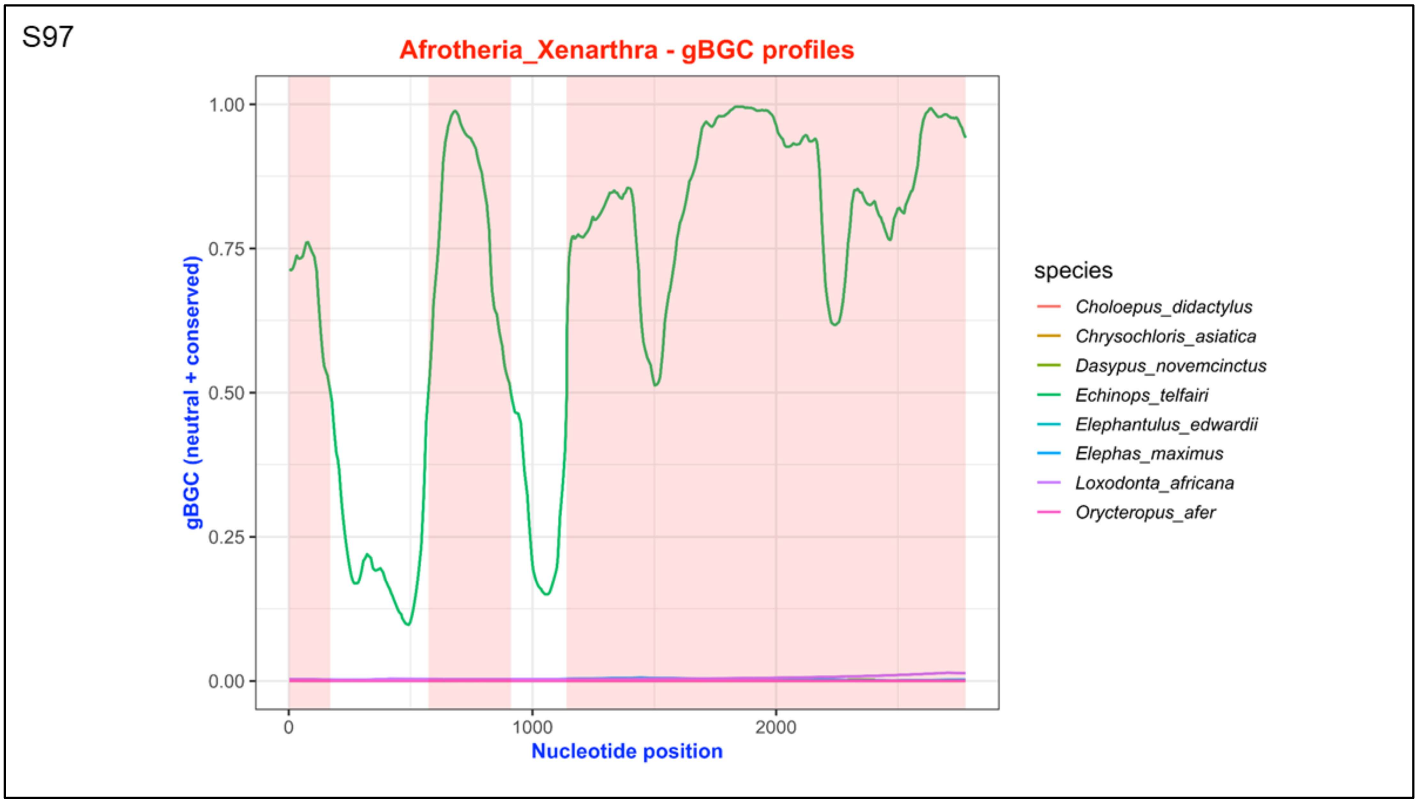


**Supplementary Figure S97. Per-base gBGC profiles of the groups Afrotheria and Xenarthra:** The X-axis represents nucleotide position, and the Y-axis depicts gBGC values. Each line illustrates the per-base gBGC pattern for an individual species.

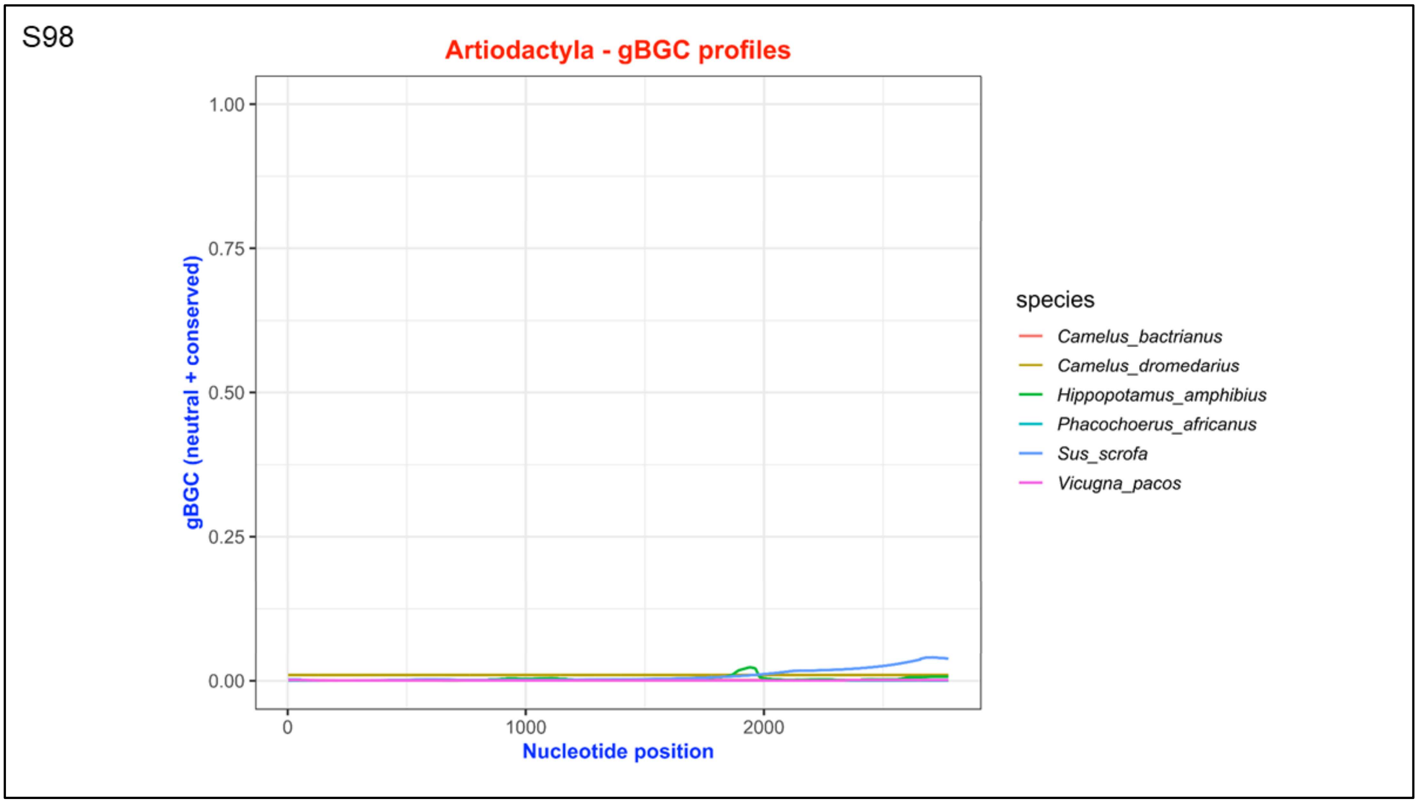


**Supplementary Figure S98. Per-base gBGC profiles of the group Artiodactyla:** The X-axis represents nucleotide position, and the Y-axis depicts gBGC values. Each line illustrates the per-base gBGC pattern for an individual species.

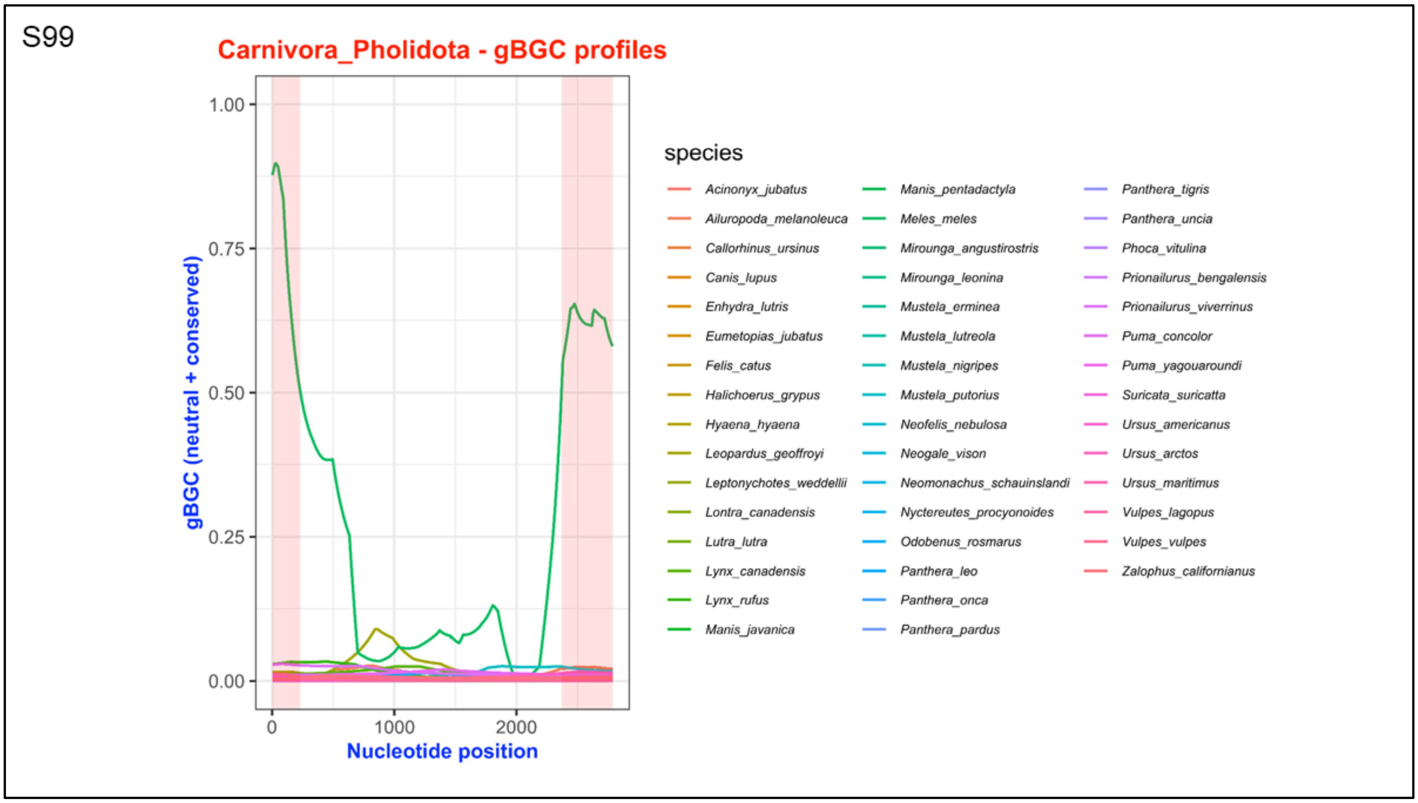


**Supplementary Figure S99. Per-base gBGC profiles of the groups Carnivora and Pholidota:** The X-axis represents nucleotide position, and the Y-axis depicts gBGC values. Each line illustrates the per-base gBGC pattern for an individual species.

S100

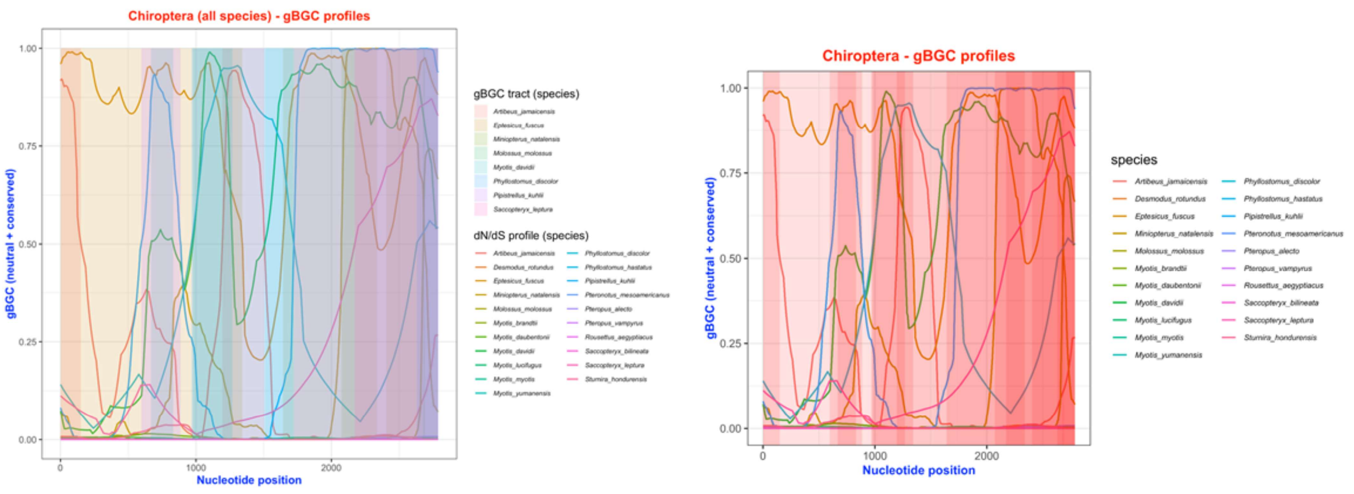


**Supplementary Figure S100. Per-base gBGC profiles of the group Chiroptera:** The X-axis represents nucleotide position, and the Y-axis depicts gBGC values. Each line illustrates the per-base gBGC pattern for an individual species.

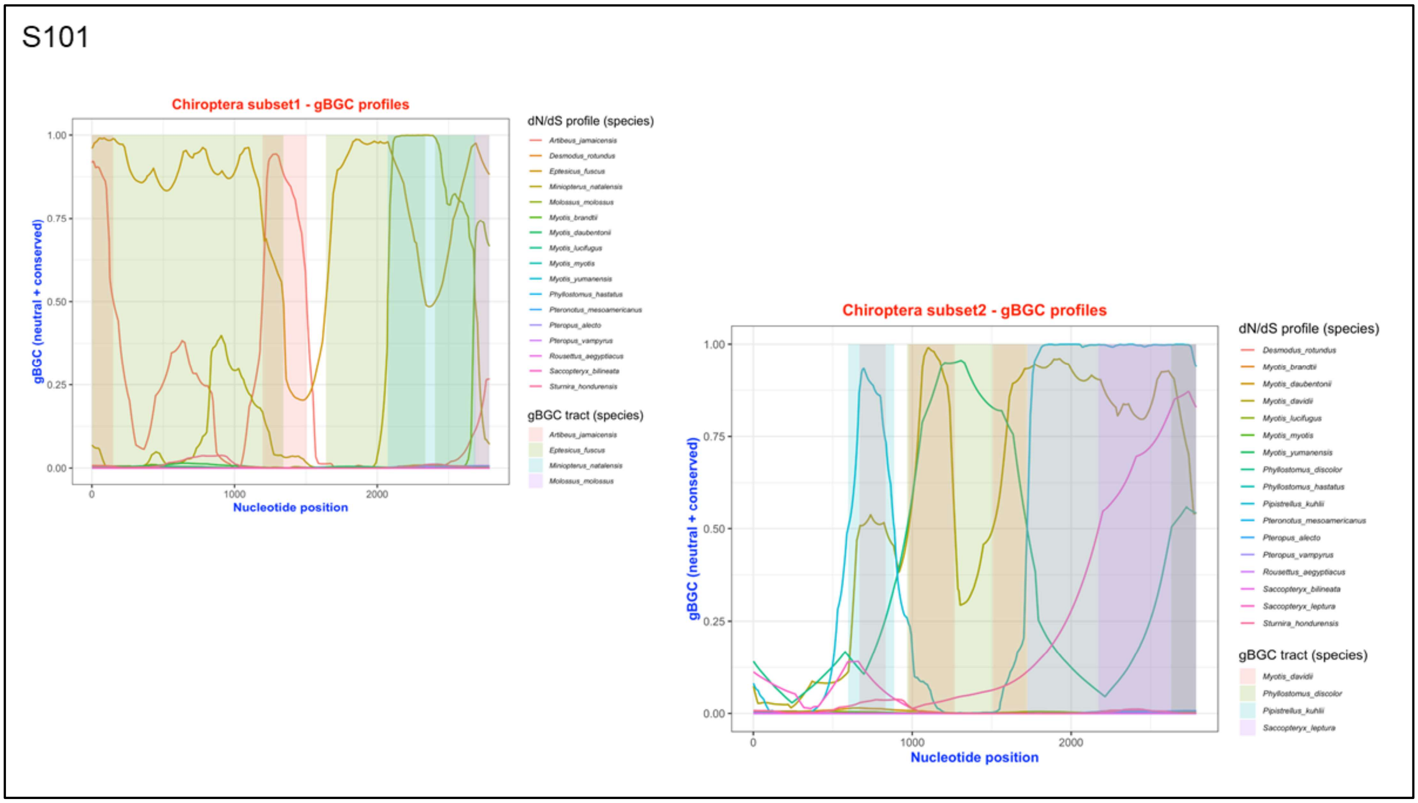


**Supplementary Figure S101. Per-base gBGC profiles of the group Chiroptera in subsets:** The X-axis represents nucleotide position, and the Y-axis depicts gBGC values. Each line illustrates the per-base gBGC pattern for an individual species.

S102

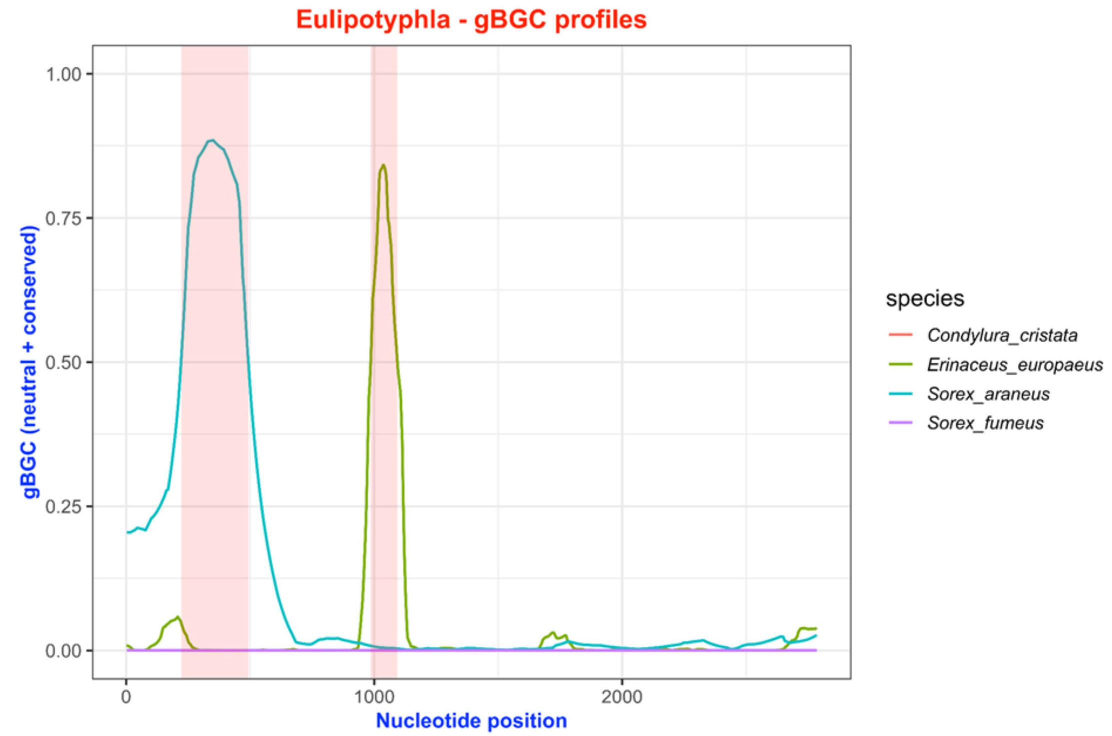


**Supplementary Figure S102. Per-base gBGC profiles of the group Eulipotyphla:** The X-axis represents nucleotide position, and the Y-axis depicts gBGC values. Each line illustrates the per-base gBGC pattern for an individual species.

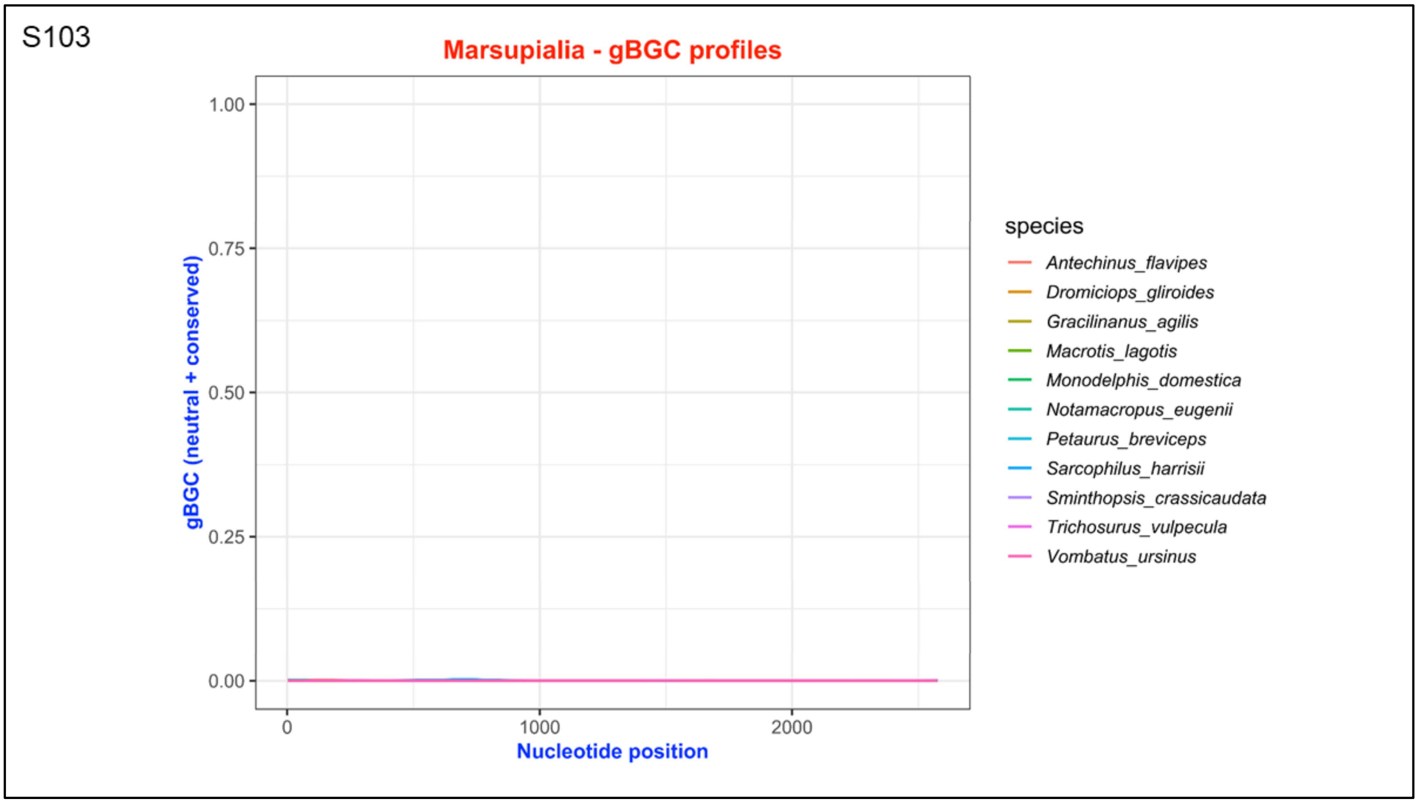


**Supplementary Figure S103. Per-base gBGC profiles of the group Marsupialia:** The X-axis represents nucleotide position, and the Y-axis depicts gBGC values. Each line illustrates the per-base gBGC pattern for an individual species.

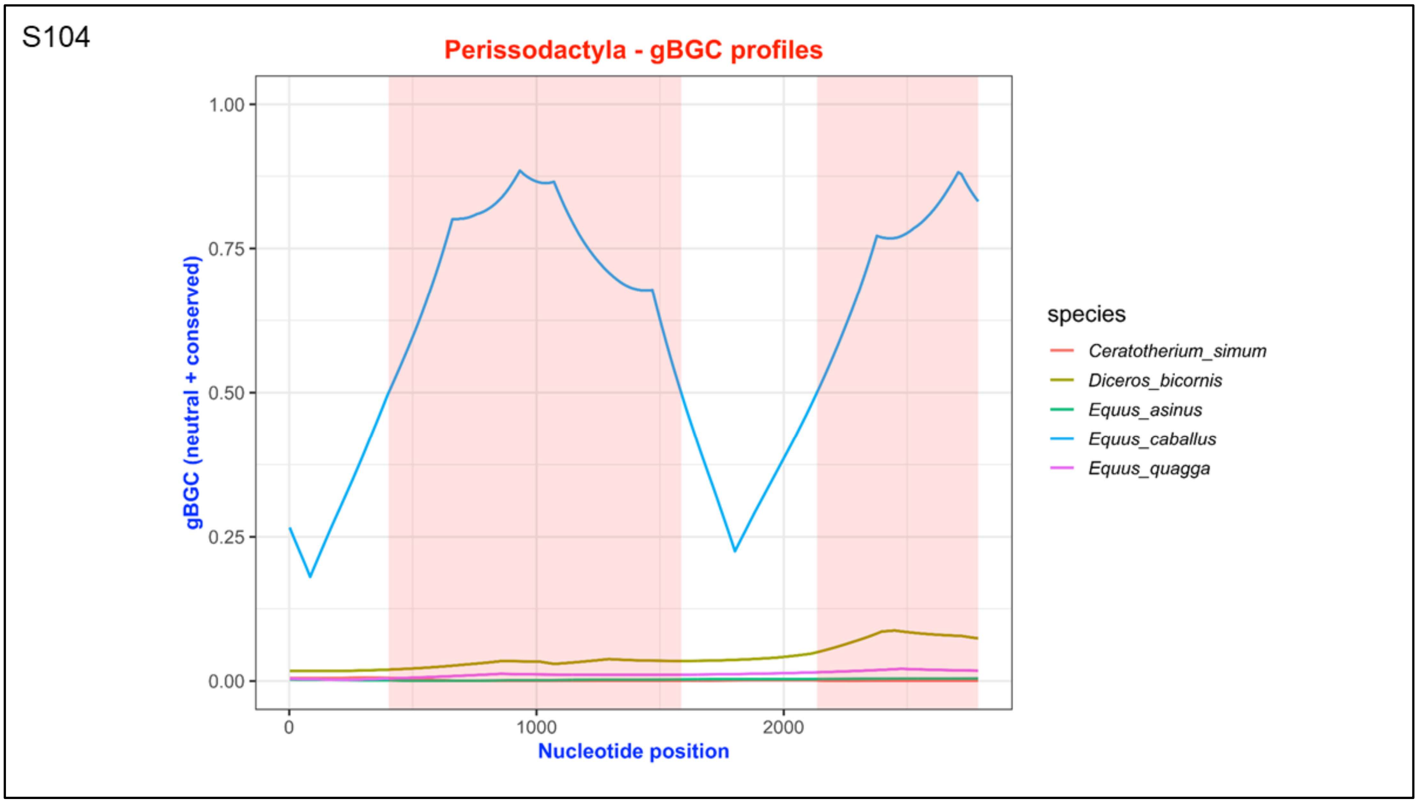


**Supplementary Figure S104. Per-base gBGC profiles of the group Perissodactyla:** The X-axis represents nucleotide position, and the Y-axis depicts gBGC values. Each line illustrates the per-base gBGC pattern for an individual species.

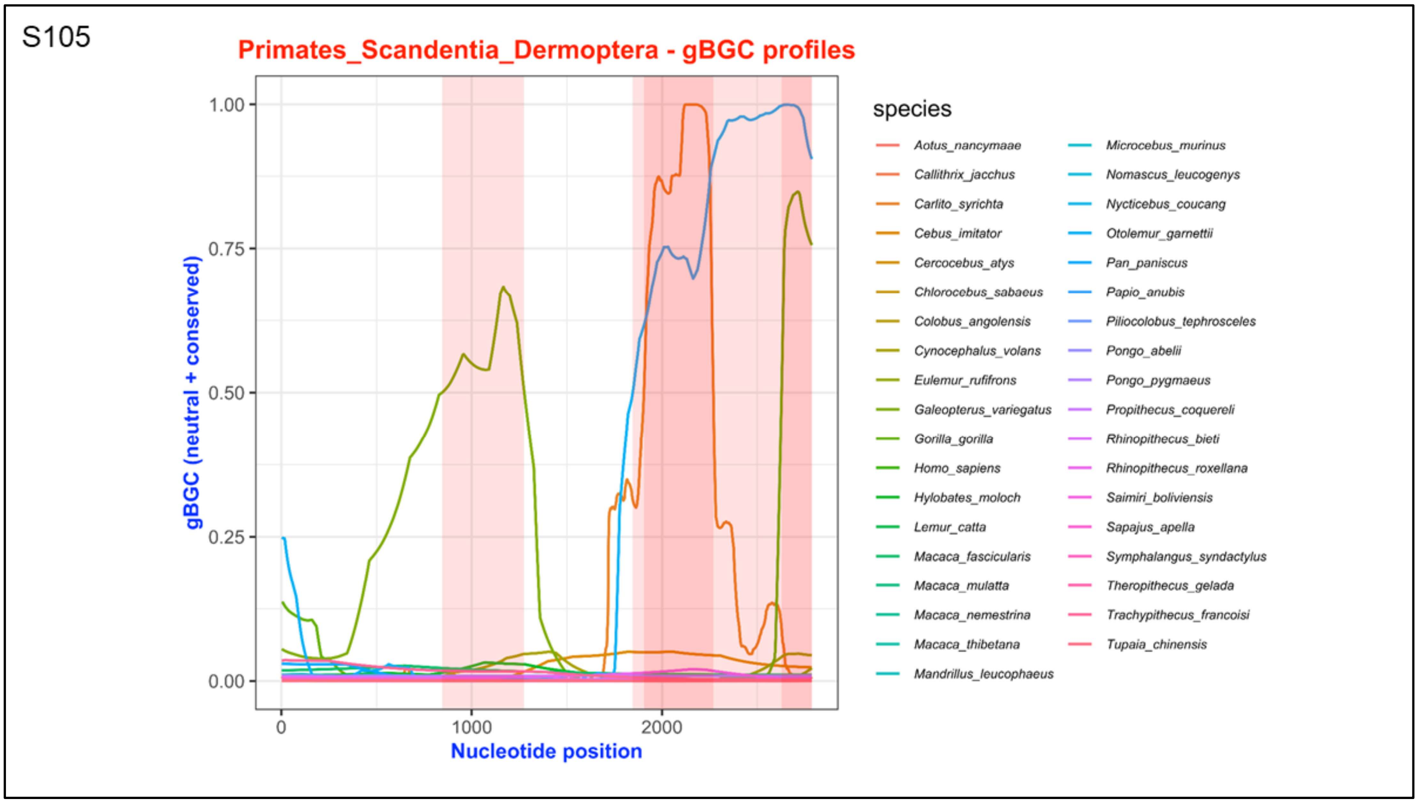


**Supplementary Figure S105. Per-base gBGC profiles of the groups Primates, Scandentia and Dermoptera:** The X-axis represents nucleotide position, and the Y-axis depicts gBGC values. Each line illustrates the per-base gBGC pattern for an individual species.

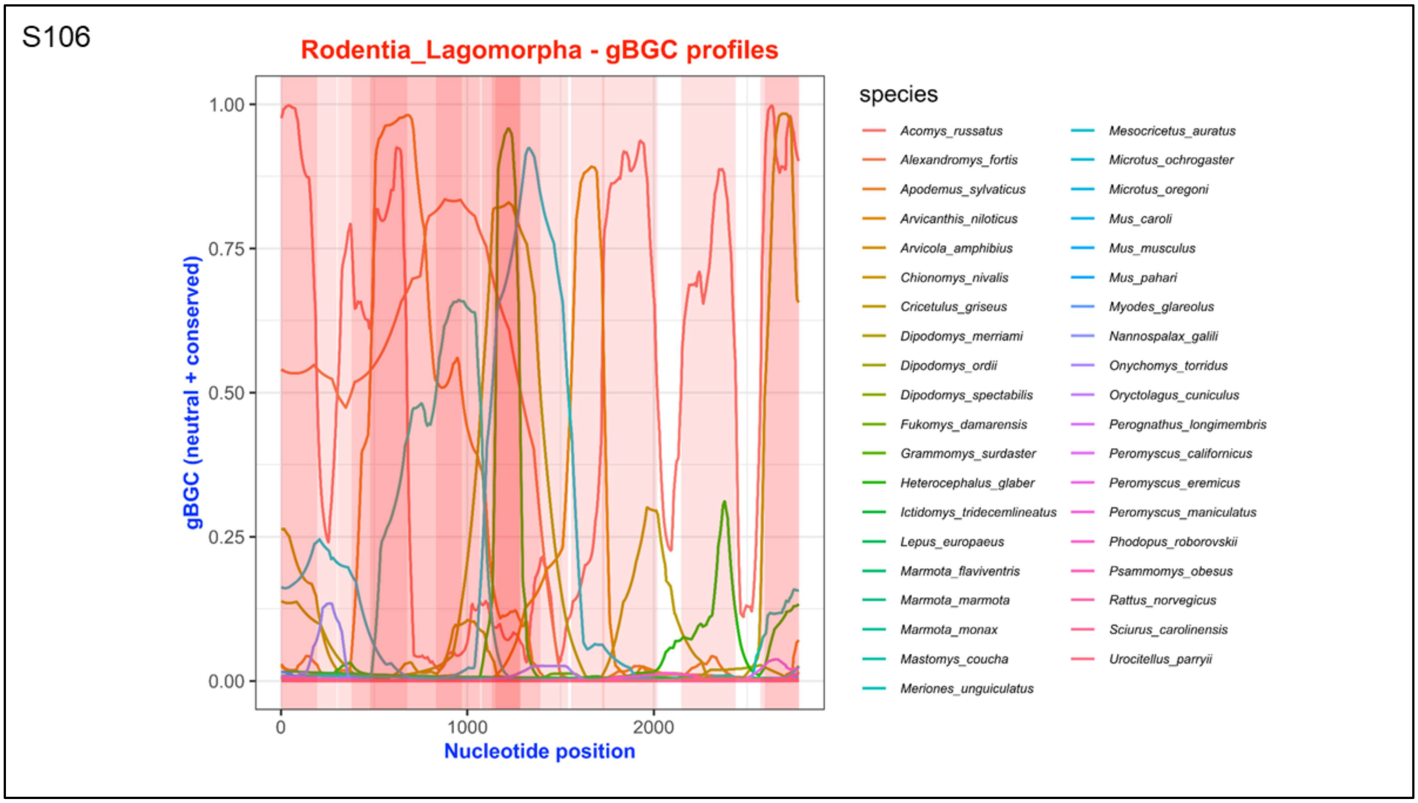


**Supplementary Figure S106. Per-base gBGC profiles of the groups Rodentia and Lagomorpha:** The X-axis represents nucleotide position, and the Y-axis depicts gBGC values. Each line illustrates the per-base gBGC pattern for an individual species.

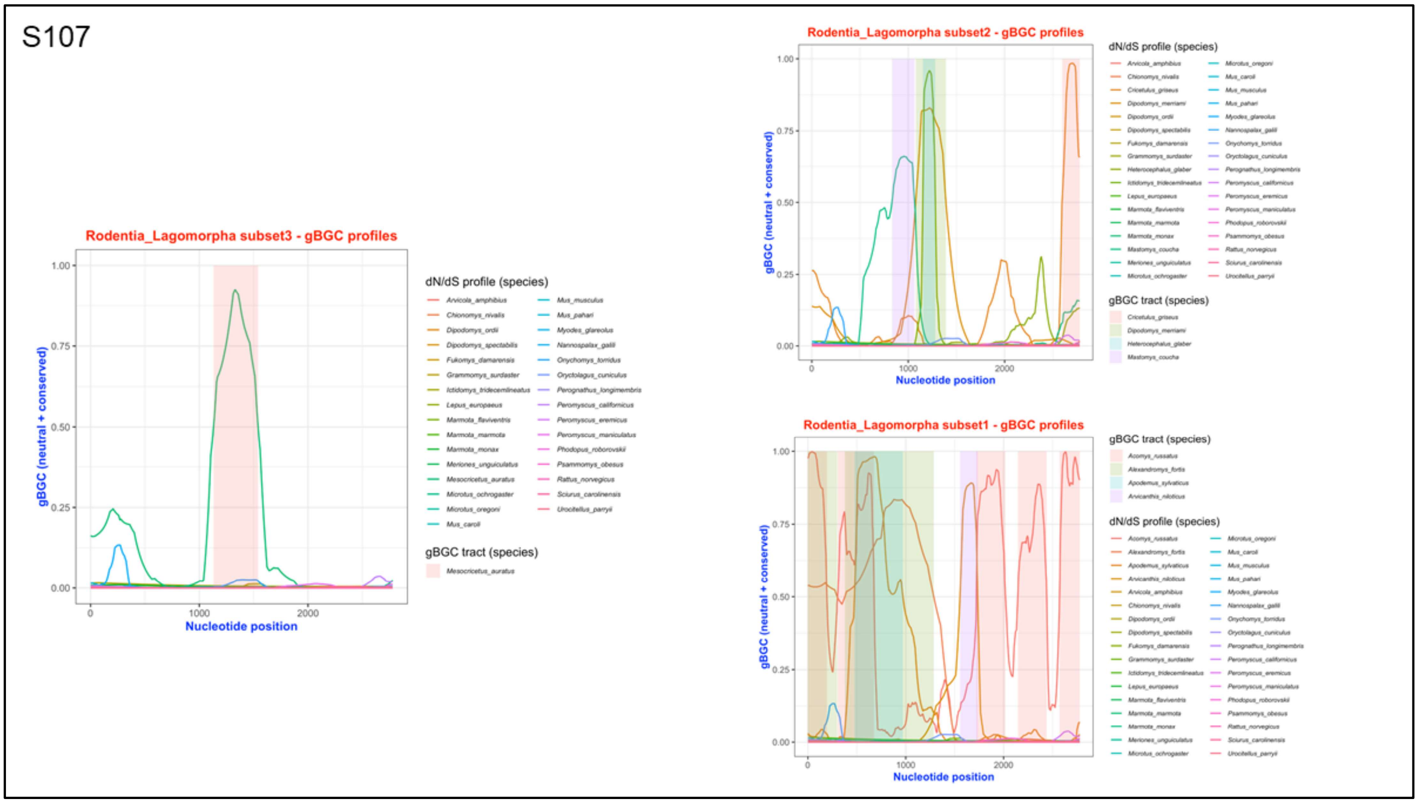


**Supplementary Figure S107. Per-base gBGC profiles of the groups Rodentia and Lagomorpha subsets:** The X-axis represents nucleotide position, and the Y-axis depicts gBGC values. Each line illustrates the per-base gBGC pattern for an individual species.

SX

Assembly verification in putative *GPRC6A* region from UCSC genome browser in
***Tachyglossus aculeatus***.

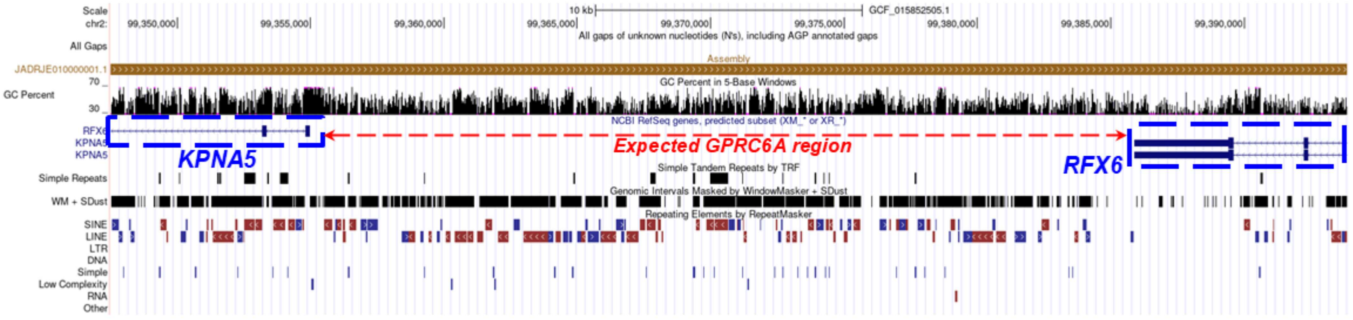


**Supplementary Figure S57. Assembly verification of *GPRC6A* region in *Tachyglossus aculeatus* in UCSC Genome browser**: This figure illustrates expected *GPRC6A* region in *Tachyglossus aculeatus* in the latest assembly of *Tachyglossus aculeatus* as available in UCSC genome browser. The two blue boxes indicate the presence of adjacent genes, while the red arrows mark the expected *GPRC6A* region. The figure confirms that the assembly is intact and supporting the conclusion that the loss of *GPRC6A* in *Tachyglossus* is genuine.
