## Supplementary Text for "Master of none: *GPRC6A* gene loss is more widespread than previously known"

### 1. Gene loss events in Sirenia

Sequence disintegration was observed in both representative species of Sirenia, the manatee species (*Trichechus manatus* and *Trichechus senegalensis*) and the dugong (*Dugong dugon).* Along with the shared frameshift mutation in the first exon, in manatee, a guanine (G) deletion in the third exon also resulted in stop codons across all three forward frames. In both species, the coding sequence of *GPRC6A* lacks a functional start codon. In the manatee, this deficiency is due to a substitution in which thymine (T) was replaced by guanine (G) at the second position of the start codon (ATG to AGG). In the dugong, the start codon was lost due to a mutation that changed the sequence from ATG to ACA, resulting in the absence of the start codon. A scan for alternative start codons within the 350 bp upstream region revealed that all resulting ORFs contained premature stop codons within the first 10% of the gene. In conclusion, the shared mutations observed in both manatee and dugong, leading to premature stop codons and the loss of a functional start codon, suggest the absence of a functional *GPRC6A* gene in these species.

### 2. **Loss in Caviomorpha**

In their GenBank annotation, we discovered gene loss in the three Caviomorph rodents labelled as “Low Quality proteins”: *Cavia porcellus, Chinchilla lanigera, and Octodon degus*. To better understand gene loss within the Caviomorpha group, we extended our analyses to include the Capybara (*Hydrochoerus hydrochaeris*), a closely related species to the Guinea pig. Remarkably, we identified shared instances of frame-disrupting changes in exon 3 in both the Capybara and Guinea pig, suggesting a conserved pattern of gene disruption within this taxonomic group.

To corroborate these findings and assess the extent of gene loss across the entire Caviomorpha clade, we examined all families with representative genomes in Caviomorpha, including Caviidae (*Cavia porcellus* and *Hydrochoerus hydrochaeris*), Chinchillidae (*Chinchilla lanigera*), Dasyproctidae (*Dasyprocta punctata*), Erethizontidae (*Erethizon dorsatum*), Octodontidae (*Octodon degus*), Ctenomyidae (*Ctenomys sociabilis*), Capromyidae (*Capromys pilorides*), Myocastoridae (*Myocastor coypus*), Cuniculidae (*Cuniculus paca*), and Dinomyidae (*Dinomys branickii*). Neither Echimyidae nor Abrocomidae had any representative genomes available in public databases. Genome blasts conducted across all representative genomes revealed widespread gene disintegration, except for Erethizontidae (*Erethizon dorsatum*), which yielded an intact ORF upon performing a blast search using *Heterocephalus glaber* as the query. SRA blast results further confirmed these gene disintegration events, supporting the observed patterns. Additionally, we identified repeat insertions in the third exon of *Ctenomys sociabilis*. These results strongly suggest gene disintegration is a prevalent and consistent phenomenon across the Caviomorpha clade.

We extended our analysis to the Phiomorpha group to assess the extent of gene loss within Hystricomorpha. We examined two representative species, the dassie rat (*Petromus typicus*) and the greater cane rat (*Thryonomys swinderianus*), belonging to the families Petromuridae and Thryonomidae, respectively. The gene is intact and functional in the naked mole-rat (*Heterocephalus glaber*), a species belonging to the Bathyergidae family. We found the gene to be intact in the Dassie rat. However, we found gene disintegration in the greater cane rat. A one-base pair deletion in greater cane rats resulted in a frameshift mutation, resulting in a stop codon in exon 2. Within exon 3, we found three frameshift mutations (two insertions and one deletion), which further disintegrate the gene.

We also examined the crested porcupine (*Hystrix cristata*), the most closely related species within the Hystricomorpha. In the crested porcupine, gene reconstruction proved challenging due to the lack of genomic BLAST hits for the first two exons. When using sequences from humans or naked mole-rat (*Heterocephalus glaber*) for exons 1 and 2, the reconstruction indicated a stop codon at the end of exon 3. However, using a shorter human isoform with a truncated exon 3 shifted the stop codon to exon 6. These findings suggest the presence of different isoforms, which complicates the reconstruction and raises questions about the gene's structure. Consequently, these inconsistencies lead us to consider the possibility of gene loss in the crested porcupine. Our analysis reveals widespread gene loss across Hystricomorphs, suggesting that the disintegration of the *GPRC6A* gene may be a lineage-specific phenomenon associated with ecological and dietary adaptations.

### 3. Disruption of Structural Motif and Domain in Koala

Genome BLASTs conducted on all available marsupial genomes confirmed the *GPRC6A* open reading frame (ORF) to be intact across species, with the notable exception of the koala (*Phascolarctos cinereus*), where we identified a specific loss of exons 1 and 3 (see **Fig. 6B**).

This absence of key exon regions is an unusual anomaly. Using the Wombat's sequence for exons 1 and 3, we were able to reconstruct the ORF for *GPRC6A* in koalas. To rule out potential assembly artefacts, we reconfirmed these deletions through short-read analysis and long-read Nanopore sequencing, all of which validated the absence of these specific regions. No in-frame stop codons were identified within the gene sequence.

Additionally, a chromosome alignment in chain format between the koala and the common wombat (*Vombatus ursinus*) revealed a single chain encompassing the remaining exons and introns of the *GPRC6A* gene and the syntenic region. This observation underscores the high level of conservation in this genomic region. To assess the consequences of the exon 1 and 3 deletions, we examined functional domains that coincide with the deleted exons, considering the remaining exons (exons 2, 4, 5 and 6) are intact up to the splice sites. The domain search of the common wombat *GPRC6A* unveiled the presence of a protein structural motif known as the PBP1_GPCA-like domain, which belongs to the Periplasmic Binding Protein type 1 (PBP1) superfamily. This ligand-binding domain serves as a recognition site for a broad spectrum of amino acids [https://doi.org/10.1016/0896-6273(93) 90269-W]. As expected, this domain exhibited full coverage in the common wombat, encompassing the first four exons, with the highest overlap of approximately 60% occurring within the third exon. In contrast, the deletion of exons 1 and 3 in the koala resulted in the whole PBP1_GPCA-like domain achieving only 22.75% coverage without any discernible binding sites, leading to its classification as a non-specific hit. Furthermore, we identified another domain termed the Nine Cysteines Domain of family 3 GPCR (NCD3G), which overlaps with the common wombat exon 3 by 68%. This domain contains several highly conserved Cysteine residues predicted to form disulfide bridges [https://doi.org/10.1016/j.tibs.2004.07.009, https://doi.org/10.1016/j.neuropharm.2010.07.007]. Moreover, our CD search revealed essential amino acid sites responsible for ligand binding (P183, L237) and dimer interface (A170, S171, V412) located within exon 3 in wombats but conspicuously absent in koalas (**Fig. 6B**). Collectively, these findings provide strong evidence supporting the loss of the *GPRC6A* gene through exon deletions in the koala.

### 4. Species labelled as "LOW QUALITY PROTEINS" in GenBank with intact ORF:

#### A. Assembly errors and annotation inconsistencies.

In the olive baboon (*Papio anubis)*, exon four initially contained a stop codon, as reported in the NCBI sequence (XM_003898267.5). However, subsequent genome BLAST analyses using newer assemblies (GCF_008728515.1 and GCA_008728515.2) revealed multiple insertions within exon four that were absent in the older genome GCF_000264685.3. The older genome's sequence, validated by SRA BLAST and alignment with the human sequence, showed no insertions or stop codons, indicating sequence integrity.

Similarly, the white-footed mouse (*Peromyscus leucopus*) exhibited an initial assembly error with a reported one-base pair deletion in its genome sequence (GCF_004664715.2) when compared to the naked mole-rat (*Heterocephalus glaber)* and mouse (*Mus musculus*). Subsequent validation through SRA BLAST identified a thymine (T) instead of a deletion. The sequence retrieved from SRA maintained an intact open reading frame (ORF) without any premature stop codon, affirming the gene's functionality despite the initial discrepancy.

Annotation errors may occur, such as the omission of exons later identified through BLAST searches, as seen in the Asian elephant (*Elephas maximus*) and the white rhinoceros (*Ceratotherium simum*). In both the Asian elephant and white rhinoceros, initial genomic annotations lacked the sixth exon. In the Asian elephant, the annotation (XM_049875552.1) did not include exon 6, but a BLAST search using the African elephant (*Loxodonta africana*) revealed the intact exon on the same scaffold, confirming an intact open reading frame (ORF). Similarly, in the white rhinoceros, the NCBI annotation (XM_004422388.2) missed exon 6, but subsequent analysis using horse (*Equus caballus*) identified the complete exon on a different scaffold, correcting the annotation error and confirming the presence of an intact ORF.

### B. Stop Codons and Potential Gene Functionality

In the genome of the Iberian mole (*Talpa occidentalis*), a stop codon was identified in the third exon. Using the European hedgehog (*Erinaceus europaeus*) isoform with a shorter third exon as a query (XM_007521031.2) in a BLAST search against the Iberian mole's genome, the entire corresponding sequence was retrieved. This indicates no conclusive evidence of gene loss in the Iberian mole. Despite the stop codon, the successful retrieval of the complete sequence suggests the gene is intact and may still be functional. Further studies are needed to determine the exact implications of the stop codon.

In the North American beaver (*Castor canadensis*), a stop codon was identified in the third exon, as confirmed through SRA BLAST analysis. However, using a human isoform

(NM_001286354.1) with a shorter third exon as a query revealed an intact open reading frame (ORF). This discrepancy suggests that the reported stop codon may be inaccurate or part of a longer isoform, while a shorter isoform may produce a functional protein. Further investigation is needed to determine the exact nature of this sequence variation and its implications for the gene's functional integrity in the North American beaver.

### 5. Phylogenetic analysis

In this study, we used two types of phylogenies: species and gene trees. All the species tree-based phylogenies were downloaded from the TimeTree website and reflect species trees inferred by earlier studies. Gene trees were generated using IQ-TREE 3 (Minh et al. 2020).

Generating genome-wide phylogenies for such a large number of species is computationally very expensive and beyond the scope of this study. Nonetheless, we explain the phylogenetic tree and the major nodes, including those with contested or ambiguous taxon placements. We used the orthofinder_synteny_supermatrix_maker.py script (available on our GitHub repository) to identify orthologous coding genes across the selected taxa. For each species, annotated genomes (genome fasta and gff files) were retrieved from NCBI using the datasets command-line interface. The longest protein isoform per gene was selected as the representative sequence. Reciprocal best BLAST (RBH) searches were conducted between a designated reference species and all others to define putative one-to-one orthologous groups, retaining only genes in ≥90% of taxa. To ensure high-confidence ortholog identification, blast hits were filtered to retain only pairs showing at least 25% amino acid identity and a minimum of 50% sequence coverage, calculated as the ratio of aligned length to the longer sequence. Gene pairs that failed these thresholds were discarded. For each orthogroup, protein sequences were aligned with MAFFT (Katoh and Standley 2013) (using the --auto strategy), and codon alignments were generated by alignment-guided back-translation of the corresponding CDS. Alignments were trimmed to remove poorly occupied columns (<60% non-missing sites) and sequences with >50% gaps, and genes shorter than 60 codons were excluded. The alignments were concatenated into a codon supermatrix with a corresponding partition file. Phylogenetic inference was performed with IQ-TREE (Minh et al. 2020) using ModelFinder (Kalyaanamoorthy et al. 2017) (-m TESTMERGEONLY, --merge-model, --merge-rate), followed by best-fit model selection (-m MFP) and ultrafast bootstrap replicates (-B 1000). Individual gene trees were estimated under the same models, concatenated into a gene-tree set, and used to compute site and gene concordance factors (--gcf, --scf) relative to the concatenated topology.

### 6. Selection analysis

#### A. Analyses of Selection Pressure

We examined the evolutionary pattern of *GPRC6A* in 166 mammalian species using three different approaches: hyphy RELAX, aBSREL, and PAML branch models. These analyses were conducted using both the gene tree and the species tree to determine whether the phylogenetic framework affected the results. For the majority of species, all methods suggested that *GPRC6A* has evolved under stable purifying selection, with no major shifts in selective pressure. However, a small number of lineages exhibited clear and statistically significant changes in selection strength, and in some cases, these patterns varied depending on the tree used.

#### B. RELAX

The RELAX analysis showed that most species did not deviate from the null expectation, meaning that selection on *GPRC6A* is generally conserved. However, a few species exhibited a relaxation of selection, where the selective pressure became weaker. This included *Felis catus*, *Heterocephalus glaber*, and *Oryctolagus cuniculus*, which showed relaxation in both the gene tree and species tree analyses. On the other hand, some species exhibited an intensification of selection, where the selective pressure became stronger. These included *Myodes glareolus*, *Urocitellus parryii*, *Phyllostomus hastatus*, *Neogale vison*, and *Homo sapiens*, and this signal was consistent across both phylogenies. In a few cases, the direction of selection depended on which tree was used. For example, *Sus scrofa* and *Aotus nancymaae* showed relaxation in the gene tree but intensification in the species tree, while *Microcebus murinus* showed relaxation only in the species tree. Similarly, *Sorex fumeus* and *Gorilla gorilla* showed intensification only when analysed using the species tree. These differences show that the direction of inferred selection change can sometimes be sensitive to the underlying phylogeny.

#### C. aBSREL

The aBSREL analysis tested whether particular branches experienced short bursts of positive selection. Across all species tested, no branches showed significant episodic positive selection.

This suggests that *GPRC6A* has not undergone rapid adaptive changes at specific evolutionary points, but rather has been shaped mainly by ongoing purifying selection across mammals.

#### D. PAML Branch Model

The PAML branch-model (model=M2) results supported the RELAX findings. In many species, the foreground ω was lower than the background ω, which indicates intensified purifying selection. The list of species with intensified selection included *Microcebus murinus, Otolemur garnettii, Tupaia chinensis, Arvicanthis niloticus, Microtus ochrogaster, Myodes glareolus, Molossus molossus, Sturnira hondurensis, and Neogale vison*. In contrast, positive selection ($\omega > 1$) was detected in *Sus scrofa* and *Felis catus* in the species tree analysis, and in *Sus scrofa, Ictidomys tridecemlineatus*, and *Felis catus* in the gene tree analysis. Meanwhile, relaxed purifying selection ($\omega < 1$ but fg $\omega >$ bg $\omega$) was detected in *Oryctolagus cuniculus* and *Aotus nancymaae* in both analyses, matching the direction detected by RELAX.
